# Molecular underpinnings of hornwort carbon concentrating mechanisms: subcellular localization of putative key molecular components in the model hornwort *Anthoceros agrestis*

**DOI:** 10.1101/2024.06.24.596764

**Authors:** Stephanie Ruaud, Svenja I Nötzold, Manuel Waller, Florian Galbier, Sabereh S. Mousavi, Mark Charran, Samuel Zeeman, Aurélien Bailly, Celia Baroux, Michael Hippler, Susann Wicke, Péter Szövényi

## Abstract

- Biophysical carbon concentrating mechanisms (CCMs) operating at the single-cell level have evolved independently in eukaryotic algae and a single land plant lineage, hornworts. An essential component for an efficient eukaryotic CCM is a pyrenoid whose biology is well-characterized in the unicellular green alga, *Chlamydomonas reinhardtii*. By contrast, pyrenoids and CCM are little understood in hornworts.
- Here, we investigate the molecular underpinnings and dynamics of hornwort pyrenoids. We do so by studying the subcellular localization of candidate proteins homologous to essential CCM genes in *C. reinhardtii* and assessing their mobility kinetics in the hornwort model *Anthoceros agrestis*.
- We provide evidence that an EPYC1 analog and the RuBisCO co-localize in the pyrenoid but pyrenoids seem less dynamic in *A. agrestis* than in *C. reinhardtii*. We further found that a carbon anhydrase homolog (CAH3) localizes to the pyrenoid, while an LCIB-like homolog is less intimately linked to the pyrenoid than in *C. reinhardtii*.
- Our results imply that the pyrenoid-based CCM of hornworts is characterized by a mixture of *Chlamydomonas*-like as well as hornwort-specific features which is in line with its independent evolutionary origin. Using these observations, we provide a first mechanistic model of hornwort CCM.

## Introduction

Increased photosynthetic efficiency is often achieved by concentrating and directly shuttling CO_2_ to RuBisCO, limiting its oxygenase activity (Parry *et al*., 2013; Kubis & Bar-Even, 2019). Carbon-concentrating mechanisms (CCMs) can either involve specialized tissues or be organized at the single-cell level. Vascular plants with C4 or CAM photosynthesis need complex structures involving multiple cell types and tissues that enable spatial and temporal separation of biochemical processes to increase photosynthetic efficiency (Osmond, 1978; Hatch, 1987). In contrast, cyanobacteria, several eukaryotic algae, and a single land plant lineage—hornworts—have evolved biophysical CCMs, operational at the level of a single cell (Barrett *et al*., 2021). An essential component for efficient biophysical CCM are pyrenoids and their analogs, the carboxysomes, in cyanobacteria, which are specialized compartments inside the chloroplast (Badger *et al*., 1998; Kerfeld & Melnicki, 2016; Barrett *et al*., 2021). Pyrenoids are subcellular structures containing highly concentrated RuBisCO that is scaffolded on a protein matrix (Mackinder *et al*., 2016; Itakura *et al*., 2019; Meyer *et al*., 2020). CCM is achieved by inorganic carbon transporters and carbonic anhydrases enabling shuttling and release of CO_2_ in the direct vicinity of RuBisCO, thereby reducing photorespiration and improving photosynthetic efficiency (Wang *et al*., 2015; Mackinder, 2017). Biomolecular mechanism as well as key molecular components of pyrenoid-based CCM has been intensively investigated in the unicellular alga, *Chlamydomonas reinhardtii* (Mackinder, 2017; Barrett *et al*., 2021; Crans & Jonikas, 2023).

While pyrenoids and biophysical CCM occur in a wide range of algal lineages, only a single lineage of land plants, the hornworts, have plastids containing pyrenoids that are capable of biophysical CCM (Villarreal & Renner, 2012; Li *et al*., 2017). The hornworts is one of three large clades of bryophytes thus more closely related to land plants than it is to algae such as *C. reinhardtii* (Harris *et al*., 2020). Hornworts are the only land plant lineage capable of biophysical CCM which is supported by multiple lines of evidence (Vaughn *et al*., 1990; Smith & Griffiths, 1996). Pyrenoid-forming hornworts often possess only a solitary, giant chloroplast per cell, like pyrenoid-bearing algae (Villarreal & Renner, 2012; Villarreal & Renzaglia, 2015; Li *et al*., 2017). In contrast, pyrenoid-free hornwort species have smaller plastids, present at higher copy numbers per cell – just like other land plants (Vaughn *et al*., 1992d; Frangedakis *et al*., 2023). Furthermore, the RuBisCO is concentrated in the pyrenoids in pyrenoid-bearing, whereas it is dispersed in the chloroplast stroma in pyrenoid-absent hornworts species, similarly to other land plants lacking pyrenoids (Vaughn *et al*., 1990, 1992d). Finally, isotope signature analyses of plant tissues also revealed that species with pyrenoids experience significantly higher 𝛿^13^C than pyrenoid-free hornworts and other bryophytes, providing evidence for CCM in hornwort pyrenoids (Smith & Griffiths, 1996; Hanson *et al*., 2002). Importantly, phylogenetic analysis showed that hornwort pyrenoids have been gained and lost repeatedly during the last 50 million years suggesting that the assembly of pyrenoids is potentially controlled by a few master regulators (Villarreal & Renner, 2012). Nevertheless, very little is known about the molecular basis of pyrenoid assembly and the molecular underpinnings of CCM in hornworts and it is currently unclear whether green algal and hornwort pyrenoids and CCM are regulated by convergent or divergent molecular mechanisms (Lafferty *et al*., 2023; Frangedakis *et al*., 2023).

While pyrenoids of *C. reinhardtii* and hornworts appear to be functionally homologous, they differ in multiple aspects. Pyrenoids of the alga *C. reinhardtii* are compact undissected structures occupying the middle of the large chloroplasts (Mackinder, 2017; Barrett *et al*., 2021). By contrast, pyrenoids of most hornworts are dissected into many subunits (several hundred) by stroma and thylakoid lamellae (Vaughn *et al*., 1992a; Li *et al*., 2017). In *C. reinhardtii*, pyrenoids are highly dynamic phase-separated fluid-crystal structures capable of quickly dissolving and reforming as a response to environmental change (Freeman Rosenzweig *et al*., 2017; Barrett *et al*., 2021). By contrast, hornwort pyrenoids appear to be more stable and do not immediately react to darkness or CO_2_ concentration changes (Ligrone & Fioretto, 1987; Li *et al*., 2020). Furthermore, the pyrenoid-based CCM is inducible in *C. reinhardtii* and only active under low CO_2_ concentrations. Preliminary experiments suggest that the hornwort CCM may not be inducible with low CO_2_ concentrations and CCM is active even under ambient CO_2_ concentrations (Hanson *et al*., 2002; Li *et al*., 2020). Finally, comparative genomic analyses indicate that only a few clear homologs of the *C. reinhardtii* essential CCM genes can be found in pyrenoid-containing hornwort genomes (Li *et al*., 2020). Collectively, these observations suggest that pyrenoids and their molecular biology may to some extent differ between *C. reinhardtii* and hornworts. Nevertheless, homologous and divergent properties of CCMs of hornworts and *C. reinhardtii* remain mainly unexplored.

To this end, we carried out experiments to provide the first insights into the pyrenoid-based CCM of the hornwort model *Anthoceros agrestis*. More specifically, we first identified hornwort genes homologous to essential CCM genes in *C. reinhardtii*. To narrow down the number of homologous candidates, we monitored expression of candidate genes across a range of CO_2_ conditions. We tested subcellular localization of our candidates by stably transforming *A. agrestis* plants with fluorescently labelled proteins. Finally, we investigated the mobility of putative pyrenoid components in hornworts using fluorescence recovery analysis after photobleaching (FRAP).

## Materials and Methods

### Strains and culture conditions

We cultured the hornwort *Anthoceros agrestis* (BONN or OXFORD strains) on Petri dishes containing KNOP medium (pH 5.8) (Waller *et al*., 2023) solidified with 7g/L Gelrite (Gelzan CM, G1910 Sigma). We maintained the plants by vegetative propagation (e.g., transferring small thallus fragments to new Petri dishes) in a growth chamber under 12/12h light/dark cycle (light intensity: 35 umol photons m-2 s-1), 60% humidity, and 21°C.

### Identifying *A. agrestis* genes homologous to *C. reinhardtii* CCM genes

To find *A. agrestis* genes homologous to essential CCM genes in *Chlamydomonas*, we run an Orthofinder v2.5.4 analysis with default options (Emms & Kelly, 2019) and using an extensive set of green algal, bryophyte (including hornworts), and vascular plant proteomes. The complete list of species can be found in Supplementary Table S1. We searched for orthogroups containing the following essential CCM genes in *Chlamydomonas*: RBMP1, RBMP2, SAGA1, SAGA2, EPYC1 (Meyer et al., 2020), HLA3, CAH3, CAH6, LCIB, LCIC, and LCIA (Mackinder, 2017; Emrich-Mills *et al*., 2021) (Supplementary Table S2). We inspected the orthogroups, refined their protein alignments manually, and generated maximum-likelihood phylogenetic trees using default options using the algorithm provided in Orthofinder v2.5.4 (Emms & Kelly, 2019). We also created an extended set of genes homologous to *C. reinhardtii* genes identified as part of the pyrenoid proteome by (Emrich-Mills *et al*., 2021) (Supplementary Table S2).

### Identifying a potential linker protein

To identify EPYC1 analogues (Mackinder *et al*., 2016; Wunder *et al*., 2018) in *A. agrestis*, a previously used method was adapted and applied to the complete annotated protein sequences of the *A. agrestis* [BONN] genome (Mackinder *et al*., 2016; Li *et al*., 2020). Briefly, XSTREAM (Newman & Cooper, 2007) was used with default settings, except the following (Min Word Match (i), 0.5; Min Copy No. (minC), 3; Min Period (minP), 40; Max Period (maxP), 80; Max Gaps (g), 25; Min TR Content, 0.8) to identify proteins with tandem repeats. The presence of transmembrane domains in repeat-containing proteins was assessed using TMHMM v. 2.0 (Krogh *et al*., 2001) and the disorder profile was calculated using VLXT (Romero *et al*., 2001).

### Monitoring transcription of candidates using RNA seq

Monitoring expression of candidate genes under a range of CO_2_ conditions for *A. agrestis* thalli was carried out using a LI-COR system in conjunction with hermetic chambers as described in (George *et al*., 2018). Mature thalli (from 2 to 4 months old) were employed, and each thallus, along with a portion of the medium, was transferred into the chambers. Experimental parameters were set at 90% relative humidity and 70 µmol×m^-2^s^-1^. Two chambers served as controls, containing only the medium. We used CO_2_ concentrations, spanning from very low to high CO_2_ conditions: 40, 150, 400, 800, and 2000 ppm using three biological replicates per concentration. Thalli were kept within the sealed chambers for a 24-hour period under a 12-hour light/12-hour dark cycle. The CO_2_ concentration remained stable throughout this 24-hour duration. Following the experiment, thalli were retrieved, weighed, transferred to RNA later (R0901 Sigma-Aldrich), and stored in the fridge at 4°C for one day. Subsequently, RNA later was removed, and the thalli were flash-frozen, rendering the tissue ready for subsequent RNA extraction. Total RNA extraction from the thalli was carried out using the Spectrum Plant Total RNA kit from Sigma Aldrich. The quality and concentration of the extracted RNA samples were assessed using a Bioanalyzer (Agilent 2100). Finally, stranded paired end sequencing was conducted by Novogene. For the analysis of *A. agrestis* transcriptome [BONN], Salmon (Patro *et al*., 2017) and DESeq2 (Love *et al*., 2014) were employed. The subsequent data analysis was executed using R (4.3.1), and the resulting plots were generated accordingly.

### Stable *Agrobacterium*-mediated transformation

To study the subcellular localization of selected hornwort candidate genes, we generated reporter lines expressing fluorescently labeled versions of the native coding sequence (CDS) of the candidate *A. agrestis* [BONN] gene driven by a strong promoter. We obtained stable genetic transformants applying a recently published protocol (Frangedakis *et al*., 2021; Waller *et al*., 2022) and using the *A. agrestis* BONN strain (Szövényi *et al*., 2015; Li *et al*., 2020).

In brief, we transformed one-month-old *A. agrestis* gametophytes using *Agrobacterium tumefaciens* (AGL1 or GV3101). After selection, we tested the localization of the following *A. agrestis* BONN proteins: EPYC1, LCIB, HLA3, CAH3, and RuBisCO small subunit (RSSU) (Supplementary Table S2; Supplementary Table S3). To drive the expression of the corresponding *A. agrestis* candidate genes, we used the following native promoters: Tip1;1, RSSU or LCIB (Frangedakis *et al*., 2021; Waller *et al*., 2022). Promoter sequences are given in the Sup_Info.docx file. The candidates were labeled using the following fluorescent proteins: eGFP (Cormack *et al*., 1996), mScarlet (Bindels *et al*., 2017) or mVenus (Kremers *et al*., 2006). We selected transformants using two rounds of selection on hygromycin. Plasmid constructs, their design, and sequences are available as supplementary material. Constructs were generated with loop assembly, using acceptor vectors and fluorescent tags of the OpenPlant toolkit (Sauret-Güeto et al., 2020): OP-033 CTAG-eGFP, OP-041 CTAG-mScarlet-I, OP044 CTAG-mVenus, OP-054 3TERM_Nos-35S.

### Subcellular localization of the CCM candidate proteins

To observe the localization of the labeled proteins, we pre-grow transformed plants for at least a month to reach an approximate thallus size of 1 cm in diameter. We excised small fragments of the gametophyte and placed them into a Lab-Tek chambered cover glass (#155361, Lab-Tek) together with a small piece of medium. For visualization, we used a multiphoton microscope Leica TCS SP8 MP with a water immersion objective lens (HC FLUOTAR L 25x/0.95 W VISIR). We used the confocal mode of the multiphoton microscope to detect eGFP fluorescence with an excitation laser (EX) wavelength of 476 nm and captured emitted fluorescence (EM) between 490-530 nm. For mScarlet and chlorophyll autofluorescence we used the parameters EX: 514 nm and EM: 590-620 nm, and EX: 458 nm, and EM: 675-800 nm, respectively (Figure 2A). We used the software LASX (version 3.5.7) to visualize and merge the images. For some imaging we also used a Leica TCS SPE DM5500Q confocal microscope with an ACS APO 40.0X1.15 OIL objective. Conditions for imaging were set as following EX: 488nm and EM: 490-600 nm for eGFP, EX: 488 nm and EM: 490-600 nm for mVenus, EX: 532 nm and EM: 630-800 nm for chlorophyll autofluorescence (Figure 2B, 2C, 2D, 2E).

### Fluorescence recovery analysis after photo bleaching (FRAP)

We carried out FRAP analysis to determine the mobility (diffusion) of EPYC1-eGFP and RSSU-mScarlet within the chloroplast of the hornwort species *A. agrestis* (Wunder *et al*., 2019; Atkinson *et al*., 2020). To accomplish this, we used a multiphoton microscope (Leica SP8 MP DIVE FALCON equipped with a FRAP module) with bleach and image recording settings described below. Experiments were performed on samples prepared from the transformant line carrying a labelled version of the EPYC1 analog and the RSSU encoding gene (p*-*AaTip1-EPYC1::eGFP -p-AaTip1-RSSU::mScarlet).

First, small fragments of the transformed thalli were propagated on a fresh selection plate. The following day, samples were mounted on microscopy slides. We then photobleached either mScarlet or eGFP with the following parameters: using a magnification of 25 with water immersion (NA 0.95), the field of view was set to 86.27 x 86.27 µm (in x and y, respectively, corresponding to a 26x zoom factor), and image format set to 512 x 512 pixels). A circular region of interest of x um diameter was defined for bleaching using the bleaching laser (WLL) set at 488nm or 514nm for eGFP or mScarlett respectively, set at 100% intensity, for x-y pulses over a duration of 5-25sec to achieve efficient bleaching) and a zoom factor of 5. After bleaching, the signal was recorded over 1-5 mins with 0.6-1.6 sec interval in the 590-620nm emission range for mScarlett and 1-4 mins for eGFP. Fluorescence recovery was measured over two regions of interest, one covering the bleached region (ROI1), one set in a neighbouring, non-bleached region (ROI2) for comparisons. Data were averaged and plotted using the FRAP module from the Leica software (LAS X Version 3.5.7.23225).

## Results

### Hornwort genes homologous to CCM-related genes of *C. reinhardtii*

Our Orthofinder analysis resulted in 67686 orthogroups, with an average size of 20.57 proteins (Supplementary Table S4). Of the 204 proteins reported to be associated with the pyrenoids or the CCM in *Chlamydomonas reinhardtii*, 56 formed orthogroups with hornwort proteins (Supplementary Table S2). For instance, the genes EPYC1, SAGA1, SAGA2, LCIA, and RBMP2 essential for CCM in *C. reinhardtii* do not have homologs in *Anthoceros agrestis*. For the LCIB-LCIC genes, we found a single homolog, while for HLA3, CAH3, and RBMP1, we identified several homologs in *A. agrestis*.

For the genes with more than one homolog, we built phylogenetic trees (Supplementary Figure S1) to determine the phylogenetic history of the proteins in the same orthogroup. For CAH3 we identified 7 homologs in *A. agrestis,* respectively. Of these only one gene was found in the same clade as CAH3 of *C. reinhardtii.* For RBMP1 and HLA3, we identified four and 17 homologs in *A. agrestis respectively*, but none of them occurred in the same clade as the RBMP1 and HLA3 proteins of *C. reinhardtii*. (Homologous gene ids are listed in Supplementary Table S2.

### The potential linker

From the 39981 input sequences, 12 proteins were identified following XSTREAM analysis (see XSTREAM_filtered.faa). Of those proteins, two (Sc2ySwM_117.2804.1 and Sc2ySwM_228.2813.1) contained predicted transmembrane domains. Of the remaining 10 sequences, 2 did not possess an oscillating disorder profile (Sc2SwM_362.1106.1 and Sc2SwM_369.518.1). 6 of the 8 remaining proteins contained duplicated ubiquitin-like sequences (see ubiquitin_like_alignment.txt). A representative sequence (Sc2ySwM_117.129.1) from the 6 was AlphaFold modelled along with the remaining 2 proteins (Sc2ySwM_228.417.1 and Sc2ySwM_344.1168.1). As predicted the ubiquitin-like sequence contained multiple ubiquitin-like domains (see AagrBONN_evmmodelSc2ySwM_1171291_b974d_unrelaxed_rank_001_alphafold2_ptm_m odel_3_seed_000). Of the other 2 sequences, Sc2ySwM_344.1168.1 contained a defined predicted 3D fold with homology to dehydrogenase folds (see AagrBONN_evmmodelSc2ySwM_34411681_bfa5f_unrelaxed_rank_001_alphafold2_ptm_m odel_2_seed_000.pdb and VAST results) whereas Sc2ySwM_228.417.1 showed no predicted 3D fold (see AagrBONN_evmmodelSc2ySwM_2284171_ebe1b_unrelaxed_rank_001_alphafold2_ptm_m odel_2_seed_000.pdb). Therefore, we selected the Sc2ySwM_228.417.1 gene model representing the best EPYC1 candidate for *A. agrestis* [BONN] (Supplementary Figure S2).

### Gene expression under variable CO_2_ concentrations to narrow down the list of candidates

Besides the LCIB and EPYC1 candidates, all other *C. reinhardtii* genes were part of large orthogroups and had many homologs in hornworts. We used transcriptomic data of whole gametophyte tissues under a wide range of CO_2_ concentrations to narrow down the list of our candidates (Supplementary Table S5). We found that none of the candidates reacted significantly to the variable CO_2_ concentrations (Fig. 1). This is in line with the assumption that CCM may be more constitutive in hornworts than in *C. reinhardtii* (Hanson *et al*., 2002; Li *et al*., 2017). Therefore, we assumed that bona fide candidates must be expressed under ambient, low and very low CO_2_ conditions in *A. agrestis*.

**Figure 1.**
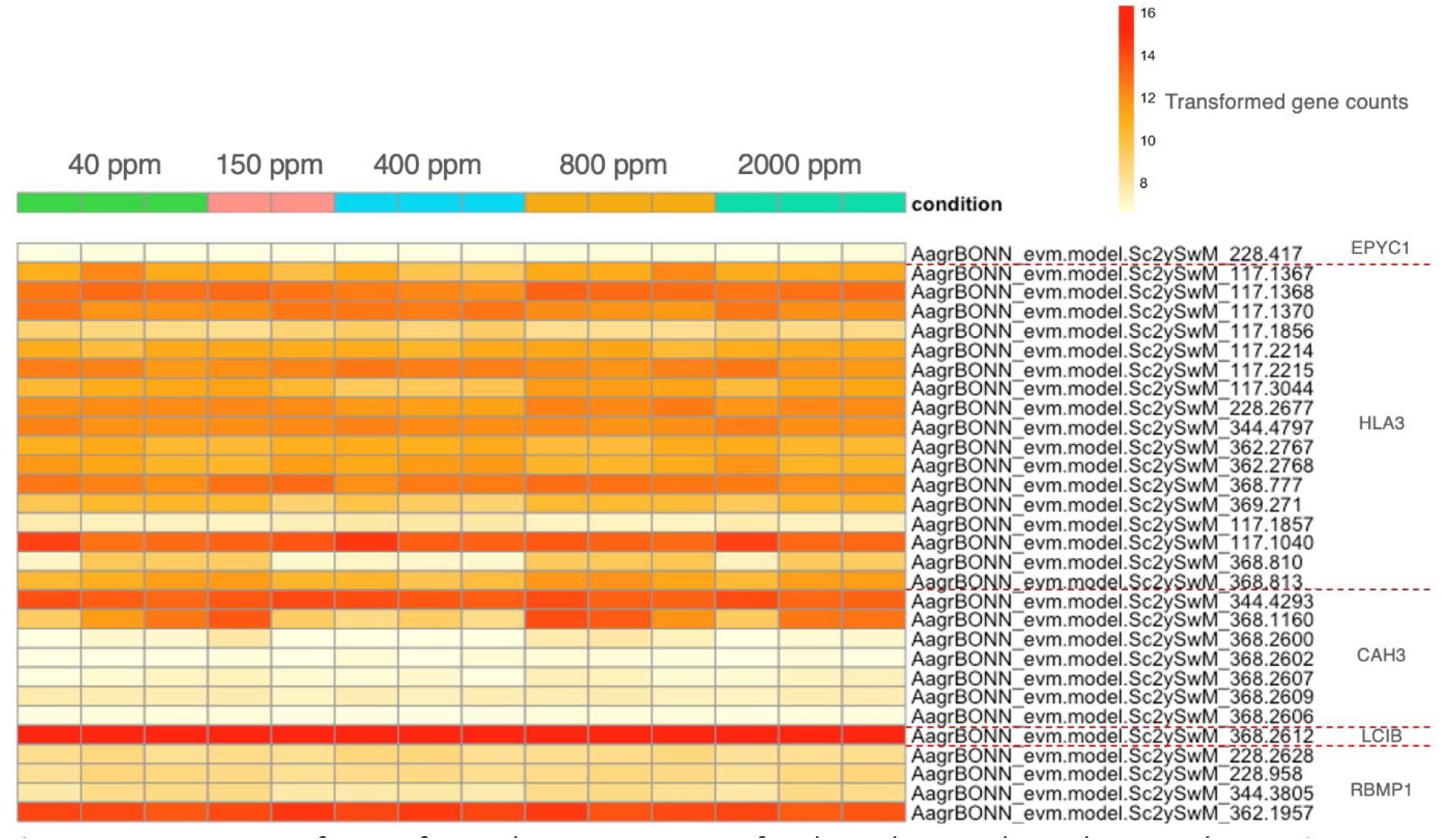
Heatmap of transformed gene counts of selected CCM homologs under various CO**_2_** concentrations in A. agrestis. Y axis: Hornwort genes homologous to essential CCM genes of Chlamydomonas reinhardtii. X axis: Five different **CO_2_** concentrations from 40 ppm (very low **CO_2_**) to 2000 ppm (very high **CO_2_**). Colored areas show transformed gene counts (using variance-stabilizing transformation (VST) on RNA-Seq data) for three (or two for 150 ppm) biological replicates.

**Figure 2.**
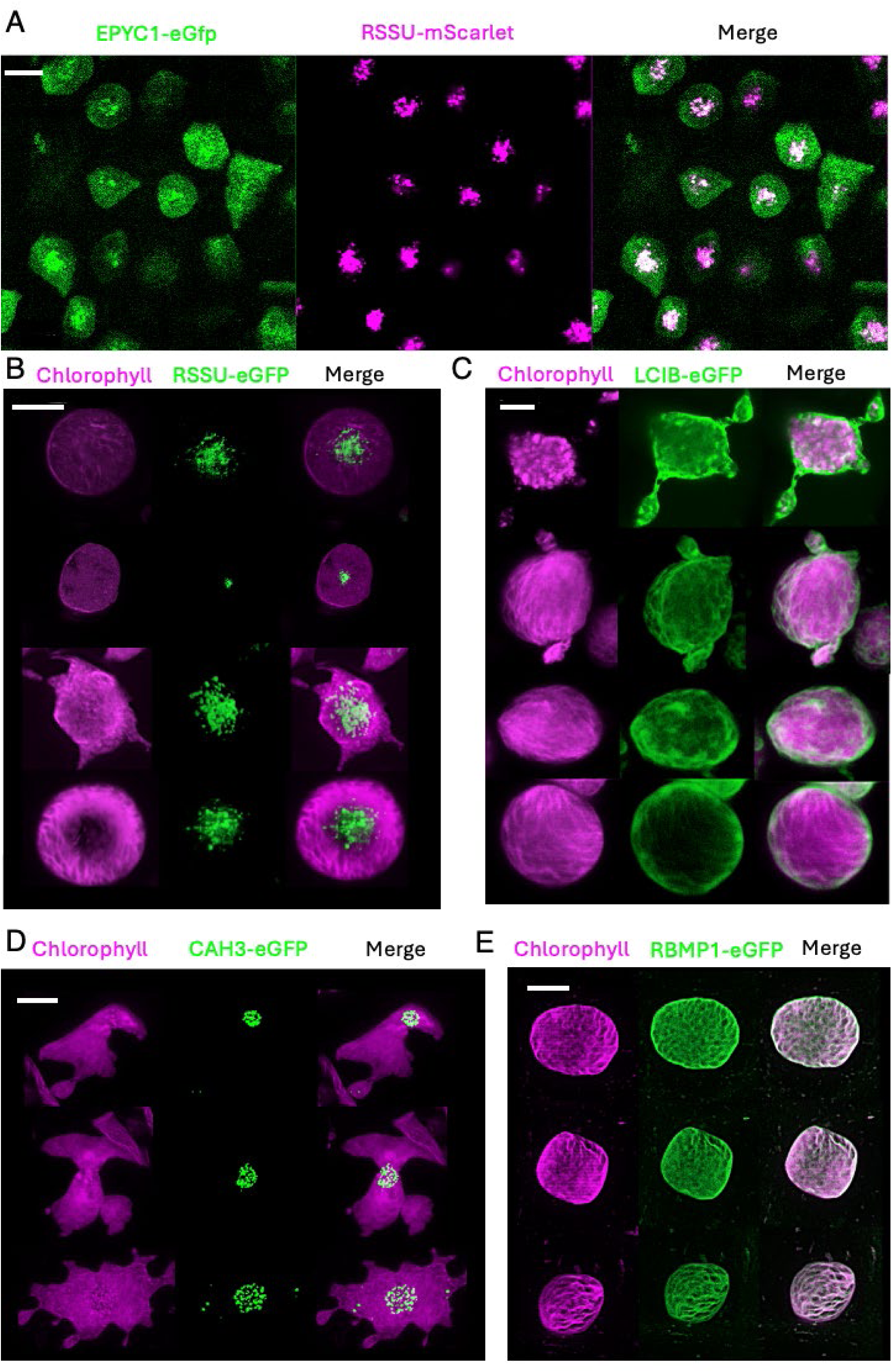
Localization of the EPYC1 analog and RuBisCO small subunit encoding gene (RSSU) in A. agrestis cells A. Confocal microscopy of the EPYC1 analog fused with eGFP (green) and the RSSU protein fused with mScarlet (magenta) both are driven by the strong AaTip1;1 promoter (Scale bar, 10 μm). Localization of RSSU B. Confocal microscopy of RSSU protein fused with eGFP (green) and driven by the strong promoter AaTip1;1, Chlorophyll autofluorescence is shown in magenta (Scale bar, 20 μm). The eGFP signal overlaps with the pyrenoid matrix. Localization of LCIB C. Confocal microscopy of LCIB protein fused with mVenus (green) and driven by the strong AaTip1;1 promoter. Chlorophyll autofluorescence is shown in magenta (Scale bar, 20 μm). The localization of the mVenus signal is ring-shaped, likely localized below the envelope of the chloroplast. Localization of CAH3 D. Confocal microscopy of CAH3 protein fused with eGFP (green) and driven by the promoter of the RSSU gene (pRSSU). Chlorophyll autofluorescence is shown in magenta (Scale bar, 10 μm). The eGFP signal overlaps with the pyrenoid matrix. Some granules are found in the chloroplast stroma. Localization of RBMP1 E. Confocal microscopy of RBMP1 protein fused with eGFP (green) and driven by the promoter of the RSSU gene (pRSSU). Chlorophyll autofluorescence is shown in magenta (Scale bar, 10 μm). The eGFP signal overlaps with the thylakoids.

For the HLA3 protein, we found that among the seventeen HLA3 candidate genes identified in *A. agrestis* eight proteins did not have a chloroplast transit peptide. We observed that these eight homologs were equally well expressed under most investigated CO_2_ concentrations. Therefore, these genes were not prioritized based on expression data (Fig. 1).

We found seven genes homologous to the CAH3 gene of *C. reinhardtii* of which all of them were detectable at the transcriptomic level (Supplementary Table S5, Fig. 1). By contrast, we observed that only a single CAH3 candidate (Sc2ySwM_228.4293.1) carried a predicted chloroplast signal peptide and this gene was present in the same clade as CAH3 of *C. reinhardtii*. Therefore, we identified this gene model as our best CAH3 homolog in *A. agrestis*. While the expression intensity of this gene did not respond to the variable CO_2_ concentration, it was the most strongly expressed among the potential CAH3 candidates.

We also identified four homologs of the RBMP1 gene of *C. reinhardtii* (Supplementary Table S5). Alignment the amino acid sequences of the *A. agrestis* RBMP1 homolog with that of *C. reinhardtii* was mainly restricted to the bestrophin domain. The short amino acid motif enabling the *C. reinhardtii* RBMP1 gene to bind to the RuBisCO was missing from the *A. agrestis* homolog (Supplementary Figure S3). Similarly to the other candidate genes, none of the RBMP1 candidates showed significant expression variation across the CO_2_ concentrations studied. Nevertheless, one RBMP1 homolog was much stronger expressed than the other candidates under all CO_2_ concentrations (Fig. 1) which were chosen for the subcellular localization study. Finally, this protein was the only one that carried a predicted chloroplast signal peptide.

### Protein localization of candidate CCM proteins in the hornwort cell

#### EPYC1 analog and RuBisCO

EPYC1 is known as a main scaffolding protein present in the pyrenoid matrix in the model organism *C. reinhardtii* (Mackinder *et al*., 2016; Freeman Rosenzweig *et al*., 2017; Wunder *et al*., 2018; He *et al*., 2020). We created double transformed plants carrying the eGFP-tagged EPYC1 candidate as well as an mScarlet-tagged version of a gene encoding the small subunit of the RuBisCO holoenzyme. Both genes were driven by the very same promoter to ensure balanced expression (Supplementary Table S3). We observed that the two candidates co-localized to the pyrenoid (Fig. 2A). While the RuBisCO was almost exclusively localized to the pyrenoids, the EPYC1 analog was concentrated in the pyrenoid but also showed considerable fluorescence throughout the entire chloroplast. We also created plants that were carrying only a fluorescently labelled protein encoding the small subunit of the RuBisCO holoenzyme.

Regardless of the promoter used, we could confirm that the RuBisCO was localized to the pyrenoid (Fig. 2B). Both the RuBisCO and the EPYC1 candidate formed small granules in the chloroplast which is in line with the dissected nature of the pyrenoids in *A. agrestis* (RSSU_3D_2.avi and RSSU_movie5.avi).

#### LCIB, CAH3, and RBMP1 homologs

The LCIB/LCIC protein complex surrounds the starch sheath around the pyrenoid and prevents CO_2_ leakage in *C. reinhardtii* (Miura *et al*., 2004; Wang & Spalding, 2006). We found that the LCIB candidate protein was closely associated with the chloroplasts in *A. agrestis*. The LCIB candidate forms a ring sitting below the chloroplast envelope (Fig. 2C, LCIB_3D.avi). The ring shaped LCIB is tightly associated with the chloroplast membrane and forms a thin layer. On some images, the LCIB layer appears to show thickenings.

The *C. reinhardtii* CAH3 is a carbonic anhydrase converting HCO_3_^-^ into CO_2_ thereby creating locally elevated CO_2_ concentration in the vicinity of the RuBisCO (Sinetova *et al*., 2012). Our findings indicate that the signal of the CAH3 homolog was predominantly localized in the pyrenoids (CAH_3D.avi), with a weak dispersed signal also observed in the chloroplast stroma. Interestingly, we also detected CAH3 signal in the chloroplast stroma, forming punctate structures (Fig. 2D).

RBMP1 and RBMP2 proteins are known to be primarily localized to the tubules and to the reticulate central regions of the *C. reinhardtii* pyrenoids. We found that the investigated RBMP1 homolog of *A. agrestis* occurred throughout the chloroplast mainly but not exclusively following the thylakoid network. Our observations suggest that the RBMP1 homolog is not concentrated in the pyrenoids but is rather associated with all thylakoid membranes throughout the chloroplast (Fig. 2E, RBMP1_movie2.avi).

### The *Anthoceros agrestis* pyrenoid is a relatively stable structure

We used Fluorescence Recovery Analysis after Photobleaching (FRAP) to evaluate the mobility of the RuBisCO and the putative EPYC1 analog in the hornwort pyrenoid. To do so, we used tissue fragments of *A. agrestis* transformants expressing either EPYC1-eGFP or a mScarlet-tagged small subunit of the RuBisCO holoenzyme. In contrast to the observations made in *C. reinhardtii*, we observed only a slight recovery of RuBisCO even after four minutes (Fig. 3A). Similarly, EPYC1 did not show significant recovery after four minutes (Fig. 3B). This suggests that both proteins are immobilized in the pyrenoid of *A. agrestis*.

**Figure 3.**
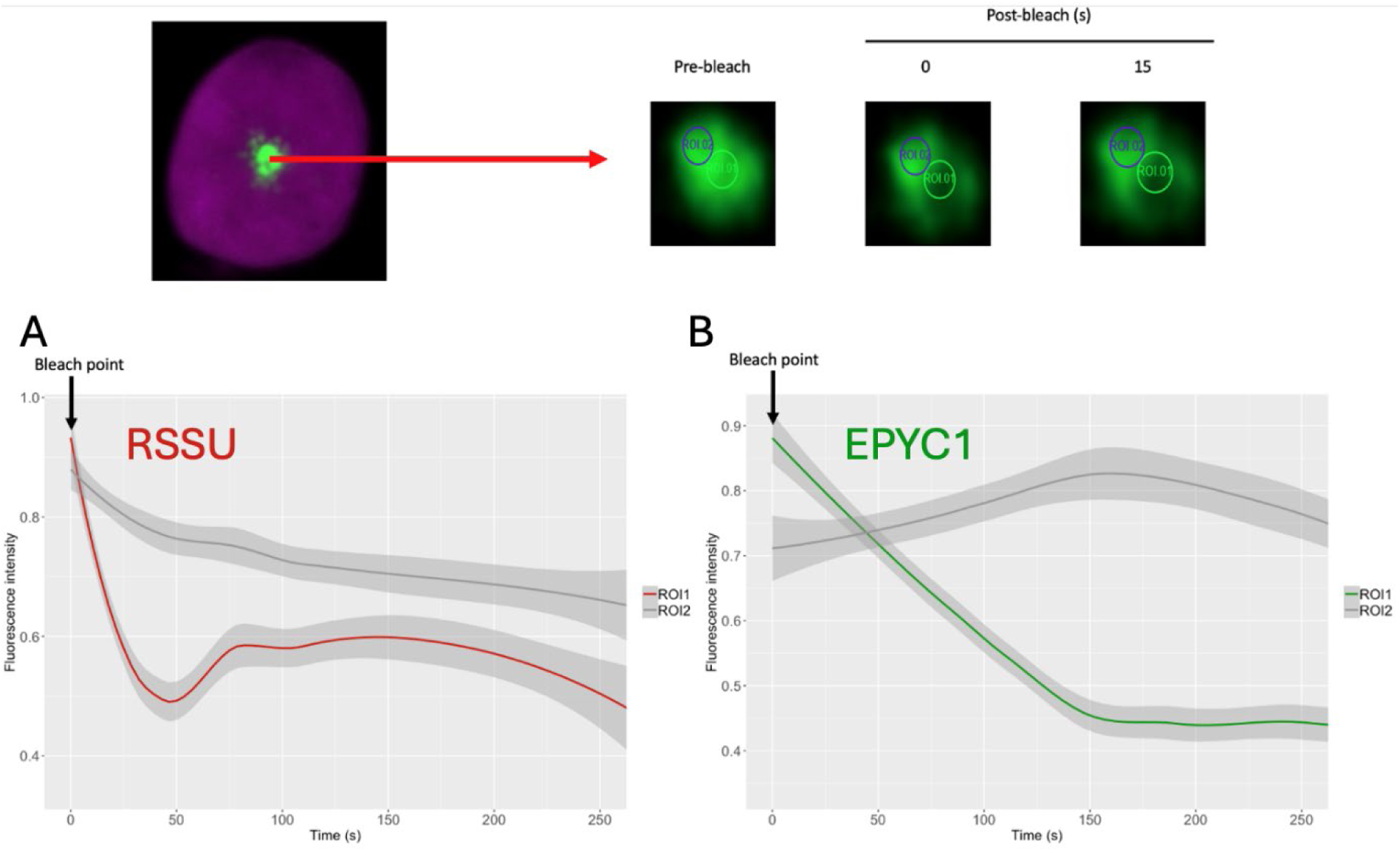
FRAP analysis of the RuBisCO and the EPYC1 analogs in A. agrestis. The top panel shows an example of a pyrenoid and the localization of the bleached (ROI1) and unbleached (ROI2, control) areas. A. RSSU tagged with mScarlet: red curve (left panel, ROI1). B. EPYC1 analog tagged with eGFP: green curve (right panel, ROI1). Curves show mean values of six measurements (biological replicates) and the standard error of the mean (gray envelope). Gray lines represent the unbleached control samples (ROI2). N = 4 for RSSU and N = 11 for EPYC1.

**Figure 4.**
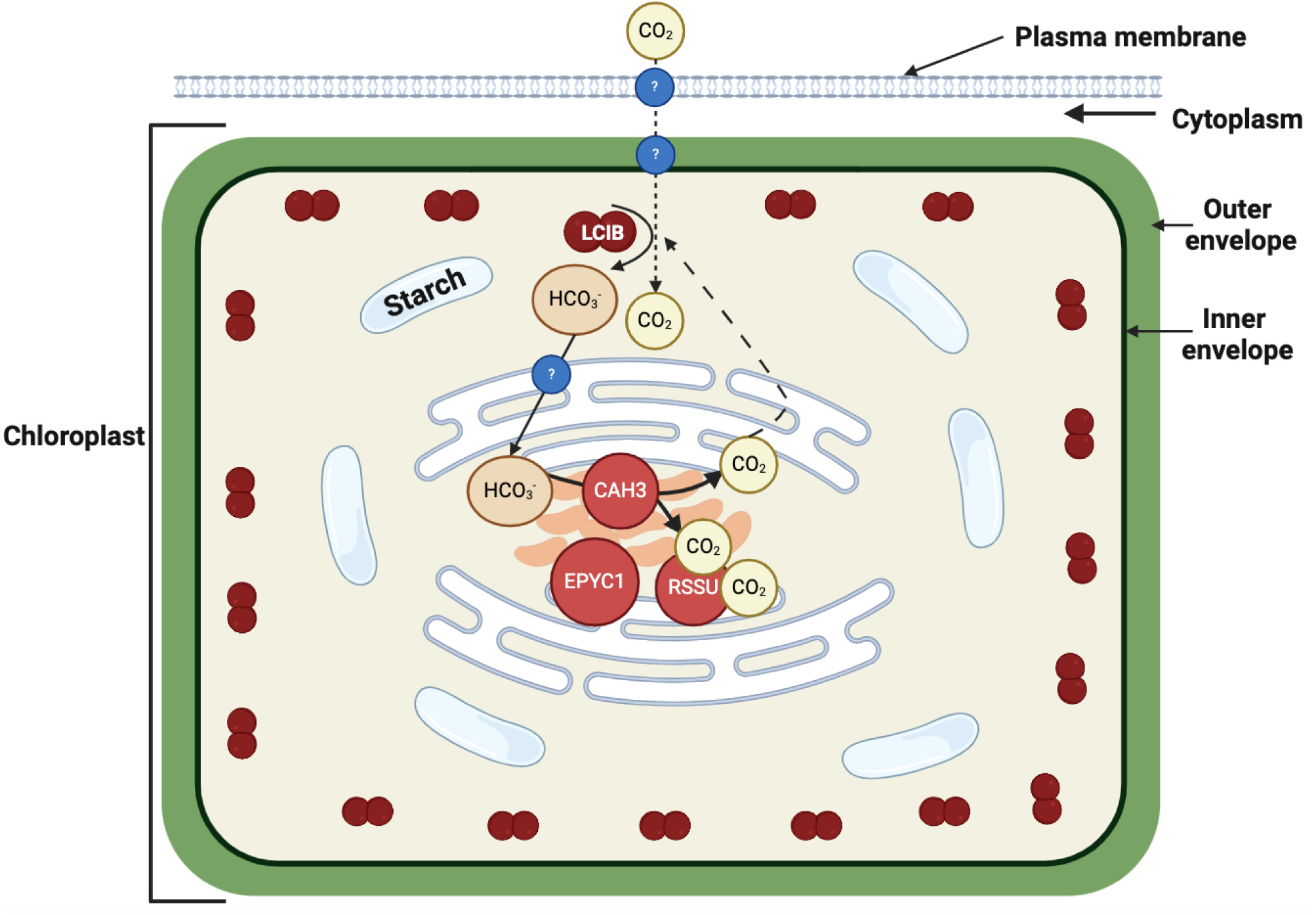
Schematic model of the hornwort CCM. The schematic representation assumes that inorganic carbon enters the cell and moves to the chloroplast as CO_2_ . The mechanism (diffusion/channels) through which CO_2_ enters the cell is unknown (see question marks). CO_2_ is hypothesized to be converted to HCO_3_-around or in the inner side of the chloroplast envelop by the LCIB homolog. HCO_3_-is transported via unknown channels/pumps (see question marks) into the thylakoids/channel thylakoids and converted into CO_2_ by the CAH3 homolog in the vicinity of RuBisCO. The leaking of unused CO_2_ is hypothesized to be captured by the LCIB homolog and converted back to HCO_3_-. Unknown channels/pumps and components of the system are labelled with questions marks.

## DISCUSSION

Pyrenoid-based carbon concentration mechanisms (CCM) are widely distributed across various groups of marine and freshwater algae and its molecular underpinnings extensively investigated in the model green alga *Chlamydomonas reinhardtii* (Giordano *et al*., 2005; Meyer & Griffiths, 2013; Fei *et al*., 2022; Wang *et al*., 2023). By contrast, very little is known about the molecular background of the pyrenoid-based CCM in hornworts, the single group of land plants in which pyrenoids occur (Hanson *et al*., 2002; Meyer *et al*., 2008; Li *et al*., 2017, 2020). Here we provide the first insight into the molecular underpinnings of hornwort pyrenoid-based CCM by assessing subcellular localization of its key components and studying the dynamics of pyrenoid recovery in the model hornwort *Anthoceros agrestis*. While our findings suggest that certain molecular components are likely shared between hornworts and *C. reinhardtii*, they also highlight divergent mechanisms concerning pyrenoid recovery and localization of key components.

### Pyrenoid-based CCM likely relies on a homologous set of genes in A. agrestis and C. reinhardtii

Phylogenetic analyses suggest that pyrenoids have originated independently in green algae and hornworts (Villarreal & Renner, 2012; Villarreal & Renzaglia, 2015; Li *et al*., 2017). Nevertheless, whether molecular underpinnings of pyrenoid-based CCM differ considerably between these two groups is unclear. In brief, key components of the pyrenoid-based CCM in the unicellular green alga *C. reinhardtii* transport HCO_3_^-^ from the extracellular environment to the pyrenoids, create locally elevated CO_2_ concentration in the pyrenoids by converting HCO_3_^-^ to CO_2_, recapture leaked CO_2_ at the pyrenoid periphery and maintain pyrenoid structure (Mackinder, 2017; Hennacy & Jonikas, 2020; Barrett *et al*., 2021). Our analysis implies that several key genes involved in the *C. reinhardtii* CCM have homologs in the hornwort genome suggesting similar underlying molecular mechanisms. For instance, we showed that a potential linker protein (EPYC1) is localized to the pyrenoid and co-occurs with the RuBisCO in *A. agrestis.* We also found that CAH3 and LCIB/LCIC homologs are localized to the pyrenoid and to the periphery of the chloroplast, respectively. Therefore, it is likely that the core molecular machinery driving pyrenoid-based CCM in hornworts relies on components homologous to the key CCM genes of *C. reinhardtii*. However, it is noteworthy that core CCM-like genes are constitutively present both on the mRNA (in this study) and protein level (Nötzold et al. 2024) in *A. agrestis* unlike in *C. reinhardtii* where CMM-related genes are usually upregulated under limited CO_2_ conditions (Mitchell *et al*., 2014; Adler *et al*., 2022). While CO_2_ concentration itself does not appear to be sufficient to induce expression of CCM-related genes in hornworts (Li *et al*., 2020), recent observations suggest that some of these genes may be induced by submersion and oxidative stress in a species-specific manner (Nötzold et al. 2024). Therefore, inducibility of CCM-related genes and CCM itself may vary across hornwort species and requires further attention.

Intriguingly, we found that most homologs were part of large gene families and hornwort genes usually formed clades with vascular plant genes lacking pyrenoid-based CCM. This suggests that CCM specialized function of homologous genes was likely acquired independently in green algae and hornworts by changing their subcellular localization and/or gene expression/regulation. This is similar to the observations made on the molecular underpinnings of C4 photosynthesis, a complex trait with over 50 independent evolutionary origins in various lineages of vascular plants (Schlüter & Weber, 2020). Therefore, improving photosynthetic efficiency is likely preferentially achieved by tinkering with the expression, localization and regulation of already existent genes. Finally, we also observed that some homologs were missing from the hornwort genomes. It is currently unclear whether the function of these genes is substituted by non-homologous hornwort genes or divergent structural organization of the hornwort and *C. reinhardtii* pyrenoids make them dispensable.

### Localization and potential CCM-related function of selected hornwort genes

We found that homologs of the CAH3 and LCIB/LCIC complex of *C. reinhardtii* are tightly co-localized with the pyrenoids and the chloroplast in *A. agrestis*. In *C. reinhardtii* the CAH3 gene catalyzes the conversion of HCO_3_^-^ to CO_2_ in the thylakoid lumen which then diffuses to the RuBisCO (Sinetova *et al*., 2012). Under ambient CO_2_ conditions CAH3 is mainly localized to the lumen of stroma thylakoids at the donor site of the PSII system (Terentyev *et al*., 2020). By contrast, at low concentration the majority of CAH3 moves to the thylakoids that cross the pyrenoid (tubules) lumen which are surrounded by RuBisCO (Sinetova *et al*., 2012). Finally, CAH3 was observed to spread out in the stroma in darkness while concentrated in the pyrenoid under moderate light conditions (Tirumani *et al*., 2014). The localization of CAH3 in *C. reinhardtii* is regulated by phosphorylation (Blanco-Rivero *et al*., 2012).

We identified a single *A. agrestis* CAH3 gene which had a chloroplast transport signal and confirmed that this protein is almost exclusively localized to the pyrenoid. We further observed that the CAH3 homolog occurs in well-delimited granule-like structures, which likely correspond to the subunits of the *A. agrestis* pyrenoid. These granules appear to spatially overlap with similar granular structures formed by the RuBisCO in the pyrenoids. Furthermore, we observed CAH3 filled vesicles in the chloroplast stroma that might originate from the Golgi and are transported into the chloroplast. Therefore, our observations suggest that the identified hornwort CAH3 candidate likely represents the functional homolog of the corresponding *C. reinhardtii* gene.

While subcellular localization of the CAH3 candidate suggests its central role in the pyrenoid-based CCM in hornworts, its function remains to be experimentally verified. Furthermore, it is unclear whether the CAH3 candidate is mainly localized to the lumen of the thylakoids traversing subunits of the *A. agrestis* pyrenoid or to the so-called channel thylakoids connecting grana thylakoids (Vaughn *et al*., 1992a; Barrett *et al*., 2021). Previous studies in *A. agrestis* showed that PSII is concentrated in the grana thylakoid membranes (Vaughn *et al*., 1992b). CAH3 is primarily associated with the donor site of PSII in *C. reinhardtii* under ambient CO_2_ conditions (Karlsson *et al*., 1998; Terentyev *et al*., 2019) which could potentially also be the case for *A. agrestis.* While this is likely, the possibility that CAH3 is localized to the channel thylakoids or thylakoids cannot be ruled out. It is also unknown whether subcellular localization of CAH3 is dependent on light/darkness and CO_2_ concentrations in *A. agrestis* as it was described for *C. reinhardtii* (Blanco-Rivero *et al*., 2012; Tirumani *et al*., 2014). Finally, experiments should be carried out to test the effect of its phosphorylation which may increase its activity and may contribute to its relocalization as seen in *C. reinhardtii* (Blanco-Rivero *et al*., 2012). Such observations would help to clarify the functional homology of CAH3 genes in *A. agrestis* and *C. reinhardtii*.

The LCIB/LCIC complex is an important component of the CCM in *C. reinhardtii* because LCIB deficient mutants can survive under very low CO_2_ and high CO_2_ conditions but not under low CO_2_ conditions (Miura *et al*., 2004; Wang & Spalding, 2006). In *C. reinhardtii* subcellular localization of LCIB/LCIC is dependent on light and CO_2_ conditions (Yamano *et al*., 2010). Under very low CO_2_ it forms a tight ring around the starch sheath of the pyrenoid, but it disperses to the chloroplast stroma under low or high CO_2_ (Yamano *et al*., 2010, 2022; Wang & Spalding, 2014a,b). LCIB has a dual role in the biophysical CCM, it can recapture leaking CO_2_ from the pyrenoid under very low CO_2_ conditions while it can facilitate CO_2_ uptake under low CO_2_ conditions. Finally, it can also actively pump in CO_2_ by converting inflowing CO_2_ to HCO_3_^-^ (Fei *et al*., 2022). These functions are achieved by its carbonic anhydrase activity converting CO_2_ into HCO_3_^-^ (Jin *et al*., 2016; Kasili *et al*., 2023).

Localization of the LCIB/LCIC homolog in *A. agrestis* appears to differ from that observed in *C. reinhardtii*. In particular, the LCIB homolog forms a thin ring-like layer likely localized under the chloroplast envelope. Therefore, LCIB is not in direct contact with the pyrenoid or with the starch granules. There are multiple non-exclusive explanations to this configuration. It can be hypothesized that the less intimate interaction with the pyrenoid and starch granules is a consequence of the divergent spatial organization of these components in *A. agrestis* and *C. reinhardtii*. In particular, the highly dissected nature of the pyrenoids and starch granules in the chloroplasts of *A. agrestis* might make direct and tight contact with the LCIB protein unfeasible (Vaughn *et al*., 1992c). Alternatively, divergent localization of the LCIB homolog in *A. agrestis* and *C. reinhardtii* may have functional significance. For instance, hornworts are thought to be rarely exposed to very low CO_2_ conditions thus inorganic carbon is likely taken up via CO_2_ diffusion and not as HCO_3_^-^. Under this scenario, LCIB may be important in converting inflowing CO_2_ into HCO_3_^-^ at the chloroplast membrane and feeding it into the pyrenoid containing stroma (Wang & Spalding, 2006; Jin *et al*., 2016; Tolleter *et al*., 2017; Fei *et al*., 2022). This hypothesis does not rule out the possibility that LCIB could function as a CO_2_ leakage barrier too. Taken together, subcellular localization of the LCIB homolog in *A. agrestis* suggests its involvement in the pyrenoid-based CCM, however, its exact molecular function remains to be clarified. Furthermore, it will be also necessary to investigate whether LCIB re-localization is triggered by CO_2_ concentration changes in hornworts as was observed for *C. reinhardtii* (Yamano *et al*., 2010, 2022; Toyokawa *et al*., 2020).

RBMP1 and RBMP2 are proteins thought to be important structural components of the *C. reinhardtii* pyrenoids anchoring the pyrenoid matrix (RuBisCO) to the pyrenoid tubules and to the reticulate region, respectively (Meyer *et al*., 2020). In *C: reinhardtii* both proteins were observed to contain short RuBisCO binding domains that are likely to be important for their recruitment to the pyrenoid and their stabilizing function. Besides this motif, both proteins contain a bestrophin domain. The structural role of RBMP1/2 is hypothesized but remains to be experimentally verified in *C. reinhardtii*. Both the RBMP1 and RBMP2 proteins are mainly localized to the pyrenoid in *C. reinhardtii*. By contrast, we found the hornwort RBMP1 homolog to occur throughout the chloroplast mainly along the thylakoid network without obvious concentration in the pyrenoid. Furthermore, we observed that the hornwort homolog of RBMP1/2 also contains a bestrophin domain, but it lacks the specific RuBisCO-binding motif. These, along with its localization, suggest it is unlikely to be involved in the structural stabilization of the hornwort pyrenoid but may be important in ion (HCO_3_^-^/chloride) transport (Adler *et al*., 2023; Nigishi *et al*., 2024).

### Hornwort pyrenoid assembly/disassembly is slow compared to C. reinhardtii

Pyrenoids of *C. reinhardtii* primarily contain RuBisCO molecules scaffolded up into a fluid-crystal by the EPYC1 protein (Mackinder *et al*., 2016; Freeman Rosenzweig *et al*., 2017; Wunder *et al*., 2018; He *et al*., 2020). In *C. reinhardtii* pyrenoids can rapidly assemble and dissolve (in the order of tens of seconds) depending on CO_2_, light conditions as well as hyperoxia (Yamano *et al*., 2010, 2022; Freeman Rosenzweig *et al*., 2017; Barrett *et al*., 2021). Pyrenoids form by liquid-liquid phase separation through a weak interaction between the RuBisCO and the EPYC1 scaffolding protein (Mackinder *et al*., 2016; Wunder *et al*., 2018; He *et al*., 2020). Assembly is mainly controlled by the phosphorylation status of EPYC1 (Turkina *et al*., 2006).

Our imaging and photobleaching experiments suggest that pyrenoid organization likely differs between *C. reinhardtii* and *A. agrestis*. We were able to identify an EPYC1 analog in the hornwort genome showing similar physical and chemical properties to the *C. reinhardtii* EPYC1 gene which colocalizes with the RuBisCO in *A. agrestis.* Nevertheless, it is unclear whether it actively interacts with the RuBisCO and whether it is involved in its scaffolding. First, we found the EPYC1 candidate to be lowly expressed under all CO_2_ concentrations investigated. This is in stark contrast to EPYC1 analogues described so far in which the scaffolding gene is always highly expressed under inductive CO_2_ conditions (Oh *et al*., 2023a; Barrett *et al*., 2024). While we started to carry out yeast two hybrid experiments to investigate the interaction between the EPYC1 candidate and the small or large subunits of RuBisCO, which could both contribute to binding based on recent work with pyrenoids in other algae and diatom systems (Oh *et al*., 2023a), our experiments led to inconclusive results due to autoactivation in the presence of the EPYC1 candidate. Therefore, currently we have no evidence that these two proteins interact in vivo and that pyrenoid formation in hornworts occurs in an analogous way to *C. reinhardtii*.

Pyrenoid assembly in *C. reinhardtii* is a rapid process which is well exemplified by the fluorescent recovery of pyrenoids within 10-14 seconds after photobleaching in plants carrying fluorescently tagged EPYC1 molecules (Freeman Rosenzweig *et al*., 2017; Wunder *et al*., 2018). Our photobleaching experiments show that RuBisCO fluorescence does not recover even four minutes after bleaching indicating its very low mobility. This is in stark contrast to the observations made in *C. reinhardtii* in vivo and in vitro indicating 50% recovery 40s after bleaching (Wunder *et al*., 2018). We made similar observations on the EPYC1 analog which is well known to show rapid fluorescent recovery (recovery half time is about 10 sec) in *C. reinhardtii* (Wunder *et al*., 2018). This suggests a considerably lower mobility of the RuBisCO as well as the potential EPYC1 candidate in *A. agrestis* compared to *C. reinhardtii*. Taken together, our results imply that pyrenoids may be less dynamic in *A. agrestis* than in *C. reinhardtii* and their assembly process may differ. It is possible that pyrenoids of *A. agrestis* may not form via liquid-liquid phase separation which is in line with their slow recovery. Alternatively, *A. agrestis* pyrenoids may form via liquid-liquid phase separation but the presence of thylakoids traversing the *A. agrestis* pyrenoids and creating hundreds of small pyrenoid subunits may results in low mobility of the RuBisCO and the EPYC1 candidates. The slow recovery of photobleached pyrenoids is in line with previous observations suggesting rigid pyrenoid structure in hornworts (Ligrone & Fioretto, 1987). Furthermore, taxonomic work implies that the presence/absence of pyrenoids is a fixed characteristic of hornwort species with no plastic response (Villarreal & Renner, 2012). Altogether, these data and other observations imply that hornwort pyrenoids are less dynamic under varied atmospheric CO_2_ conditions and more stable structures than those of *C. reinhardtii*. Nevertheless, the molecular basis of this difference is unknown and needs to be further explored. Furthermore, recent observations suggest that RuBisCO may also be of low mobility in other algal systems than *Chlamydomonas* (Oh *et al*., 2023b). Therefore, low mobility and fluorescent recovery of pyrenoid components may not be a unique feature of hornworts.

While pyrenoids may be less dynamic (de novo assembly/disassembly) in *A. agrestis* than in *C. reinhardtii,* recent observations suggest that the number and space occupied by pyrenoids in the chloroplast can rapidly change. In a companion study it was found that submersion in water efficiently induces pyrenoid formation in *A. agrestis* within the first 24 hours (Nötzold et al. 2024). By contrast, oxidative stress conditions, known to trigger CCM in *C. reinhardtii*, resulted in shape-shifting of plastids in hornworts, without notable effect on pyrenoids and CCM (Nötzoldt et al. 2024). Therefore, environmental responsiveness of pyrenoid formation is poorly understood and needs further scrutiny.

### A putative model of A. agrestis CCM

The data presented above enables us to provide a working hypothesis on pyrenoid-based biophysical CCM in hornworts. Our model assumes that hornworts rarely experience very low atmospheric CO_2_ concentrations. Therefore, inorganic carbon likely reaches the hornwort chloroplasts as CO_2_ via diffusion through the plasma membrane and the cytoplasm potentially facilitated by CO_2_ channels. Once at the chloroplast envelope, CO_2_ likely diffuses into the chloroplast stroma and its conversion to HCO_3_^-^ is catalyzed by the LCIB homolog forming a thin layer below the chloroplast envelope. This conversion is expected to create a CO_2_ gradient across the cell assuming that HCO_3_^-^ permeability of the chloroplast membrane is significantly lower than for CO_2_ (Tolleter *et al*., 2017). The exact route of HCO_3_^-^ within the chloroplast is unclear. Hornwort chloroplasts are made up of several hundreds of subunits separated by stroma thylakoids (Vaughn *et al*., 1992d). We hypothesize that HCO_3_^-^ is first transported to the thylakoids with a currently unknown mechanism because homologs of BST1-3 channels are missing from the hornwort genome (Mukherjee *et al*., 2019). Grana stacks of hornwort chloroplasts are also connected by channel thylakoids which may or may not play a role in the CCM (Vaughn *et al*., 1992d). HCO_3_^-^ in the thylakoids/channel thylakoids is converted to CO_2_. The hornwort CAH3 occurs in granules likely tightly associated with the pyrenoid subunits. The released CO_2_ is then used by the RuBisCO which is concentrated in the pyrenoid subunits. Because hornwort pyrenoids are not in direct contact with the starch granules, we assume that stroma thylakoids traversing the pyrenoids function as primary barriers to CO_2_ (Fei *et al*., 2022). The LCIB molecule at the periphery of the chloroplast may function as a secondary leakage barrier too. 3-Phosphoglyceric acid (3PGA) is then likely transported into the stroma and converted into starch.

### Unanswered and open questions

While our localization study suggests that certain mechanisms are shared between the pyrenoid-based CCM in hornworts and *C. reinhardtii*, detailed molecular underpinnings of the hornwort CCM remain to be clarified. The extension of our method to further candidate genes and targeted functional analyses of gene function (application of gene editing technics) will open the way to make significant advancement towards understanding the key components of the CCM in hornworts. A) Further experiments need to be carried out to clarify how inorganic carbon enters the cells and reaches the pyrenoids through various channels and pumps. This could be tackled by the phenotypic screening of many mutant plants. Large-scale forward genetic screens were successfully used in the unicellular alga *C. reinhardtii* to discover key components of the CCM machinery (Crans & Jonikas, 2023). B) It will also be necessary to screen further candidate genes for their localization and functional characterization. This will allow the discovery of key CCM components showing no homology to *C. reinhardtii* genes. C) Our study suggests that pyrenoids may be less dynamic in hornworts compared to *C. reinhardtii*. Nevertheless, further information on pyrenoid assembly, its dynamic and dependence on various environmental factors and the detailed structure of the *A. agrestis* pyrenoid must be understood. D) Further experiments must address the environmental responsiveness of various components of the hornwort CCM machinery to clarify its dynamic or constitutive nature compared to *C. reinhardtii*. E) Finally, the inducibility or constitutive nature of the hornwort CCM needs to be properly investigated and clarified.

## Supporting information

SupInfo

## Acknowledgements

This project was carried out as part of the Deutsche Forschungsgemeinschaft (DFG) priority program 2237: “MAdLand—Molecular Adaptation to Land: plant evolution to change” (http://madland.science), through which PS and SW received financial support (PSLJ1111/1; WI4507/9-1). Additional funding was received from the Swiss National Science Foundation (grant nos. 160004, 184826, and 212509 to PS); project funding through the University Research Priority Program “Evolution in Action” of the University of Zurich to PS and LW; a Georges and Antoine Claraz Foundation grant to AN, YY, SR, LW, MW, and PS; UZH Forschungskredit Candoc grant no. FK-19-089 and an SNSF Doc.Mobility Projekt grant no. P1ZHP3_200030 to MW, and FK-22-098 to SR; SSM was supported by a Swiss Excellence Postdoctoral fellowship.

We are very thankful to Luke MacKinder and James Barrett (University of York, UK) for providing information and plasmids related to this study as well as for Alistair J. McCormic and Aranzazú Díaz-Ramos (University of Edinburgh, UK) for providing information on their yeast two hybrid experiment.

## Author contributions

PS, SW, MH, SR designed the study. SR, MW, SSM, MC performed the plant transformations. SR, CB, MW, AB carried out the FRAP analysis, and microscopy. SN contributed to data interpretation of the data. SZ provided material, space, and resources. PS, SR, SN wrote the initial version of the manuscript, MH, PS, and SW critically reviewed it. All co-authors have read, revised and approved the final version of the manuscript.

## Supporting Information

Additional supporting information may be found in the online version of this article.

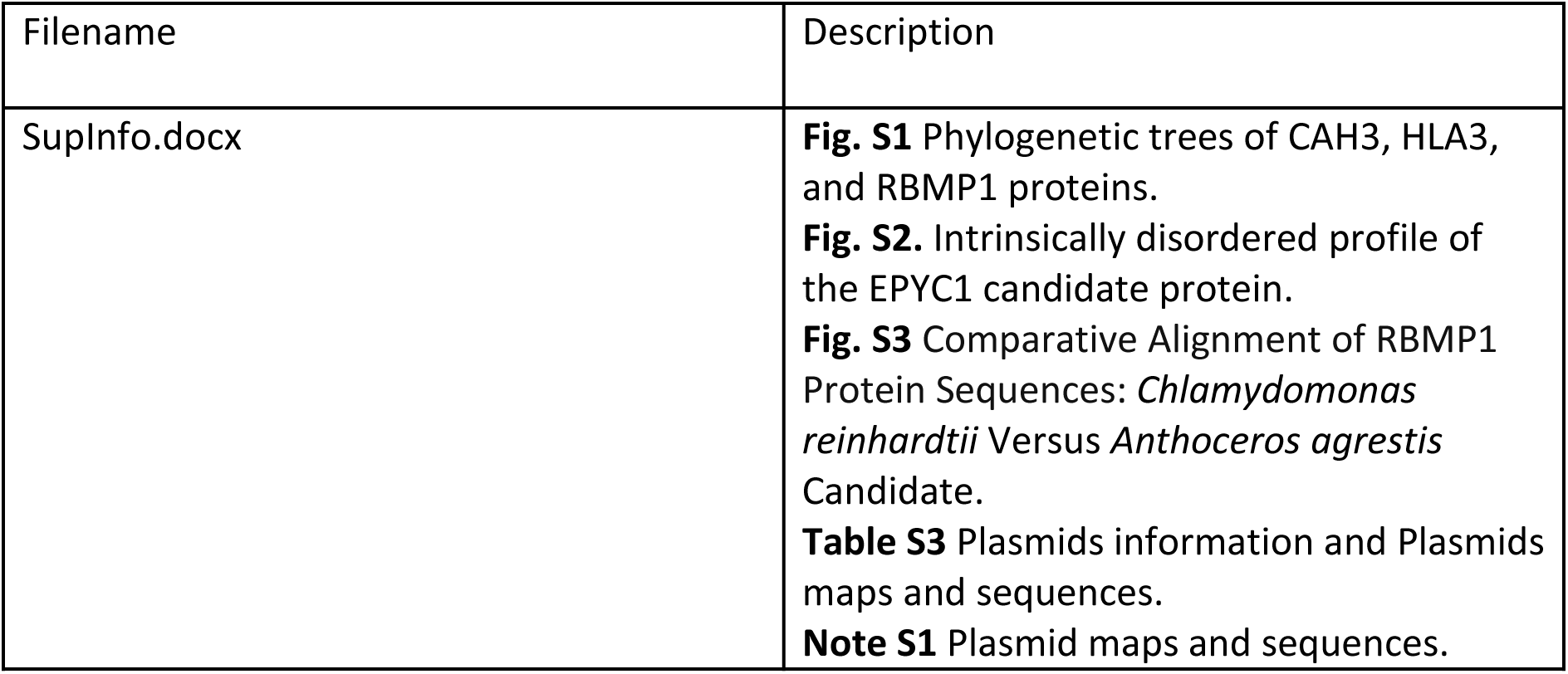

Additional supporting information (Additional files)

**Table S1** List of species and their proteomes analyzed with OrthoFinder

**Table S2** List of orthogroups and protein candidates

**Table S4** Orthogroups all species

**Table S5** Transformed gene counts of selected CCM homologs under various CO_2_ concentrations in *A. agrestis*

**RSSU_movie5.avi** 3D movie of the subcellular localization of the RuBisCO. **RSSU_3D_2.avi** 3D movie of the subcellular localization of the RuBisCO. **CAH_3D.avi** 3D movie of the subcellular localization of the CAH3 homolog. **LCIB_3D.avi** 3D movie of the subcellular localization of the LCIB homolog.

## Data availability

The raw sequences have been submitted to the NCBI Sequence Read Archive (SRA) and associated with the BioProject number PRJNA574453 (under 150, 400, and 800 ppm), and PRJNA1108481 (under 40, and 2000 ppm).

All the plasmids used in this article can be found here: https://figshare.com/s/0ea59a40c6a1dd34e2ca

Data about potential linker analysis is found here: https://figshare.com/s/54f5f37cf61cf9924458

