## Supplementary material for "Molecular underpinnings of hornwort carbon concentrating mechanisms: subcellular localization of putative key molecular components in the model hornwort *Anthoceros agrestis*": SupInfo

**Fig. S1. Phylogenetic trees of CAH3, HLA3, and RBMP1 proteins.**

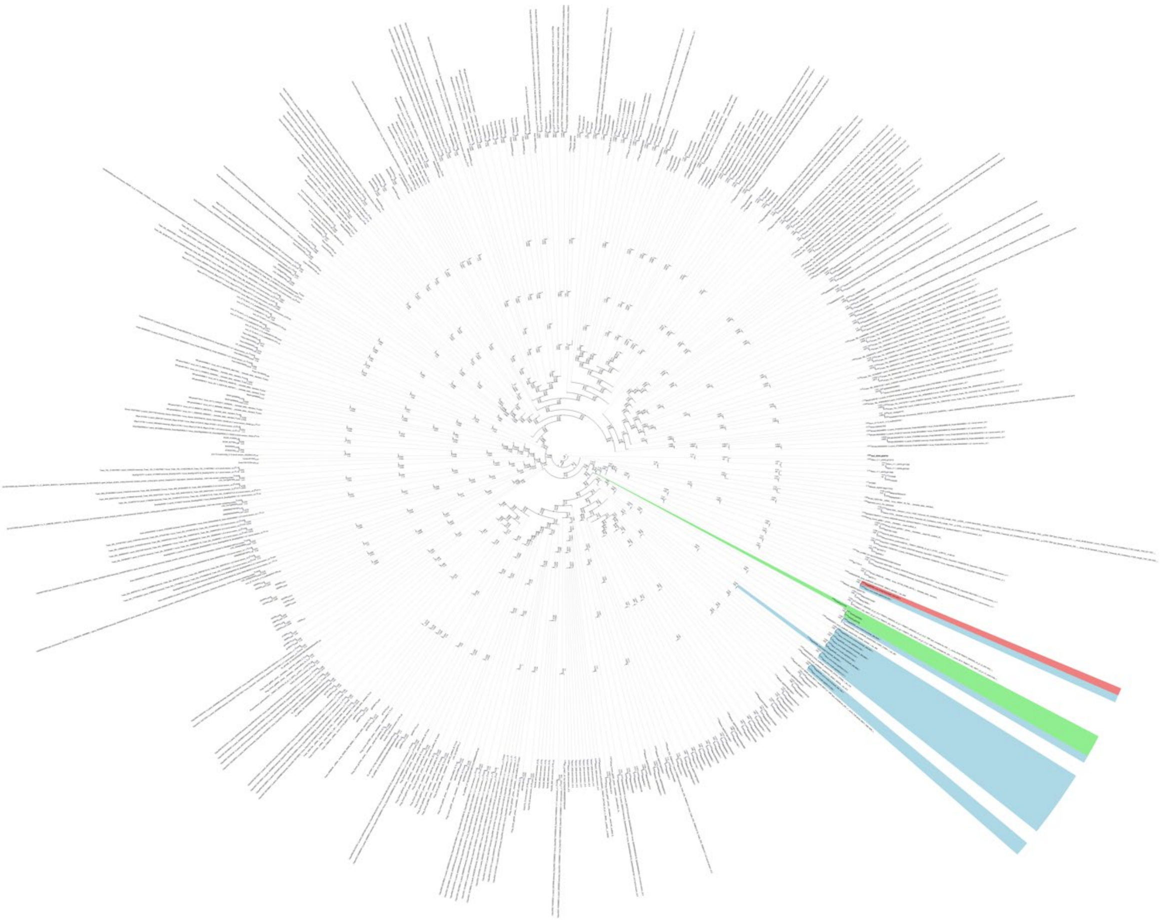

**Phylogenetic tree of CAH3 homologs.** *Chlamydomonas reinhardtii* gene ids are shown in green, those of *Anthoceros* species in blue. Gene ids highlighted in red refer to the selected candidate from *Anthoceros agrestis* BONN.

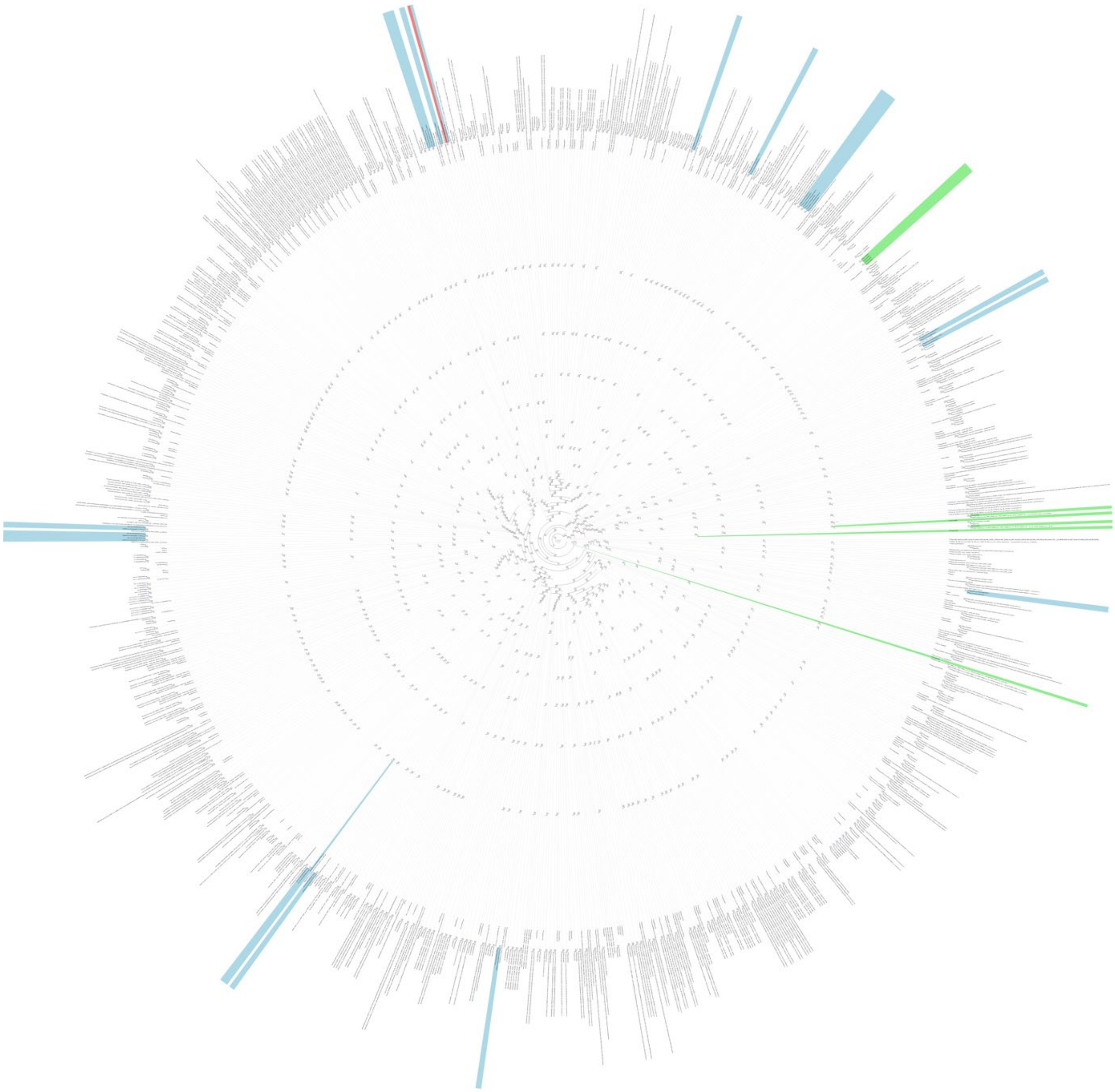

**Phylogenetic tree of HLA3 protein.** *Chlamydomonas reinhardtii* gene ids are shown in green, those of *Anthoceros* species in blue. Gene ids highlighted in red refer to the selected candidate from *Anthoceros agrestis* BONN.

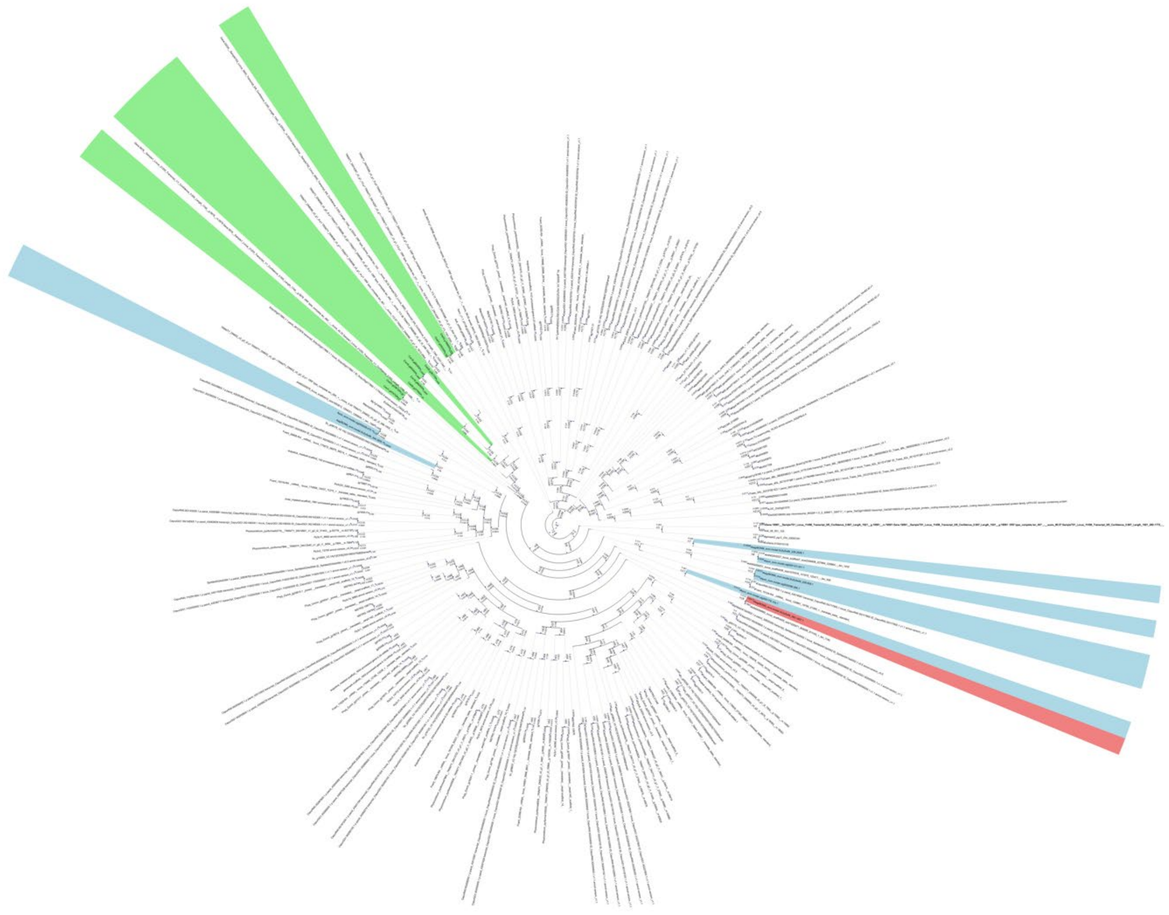

**Phylogenetic tree of RBMP1 protein***. Chlamydomonas reinhardtii* gene ids are shown in green, those of *Anthoceros* species in blue. Gene ids highlighted in red refer to the selected candidate from *Anthoceros agrestis* BONN.

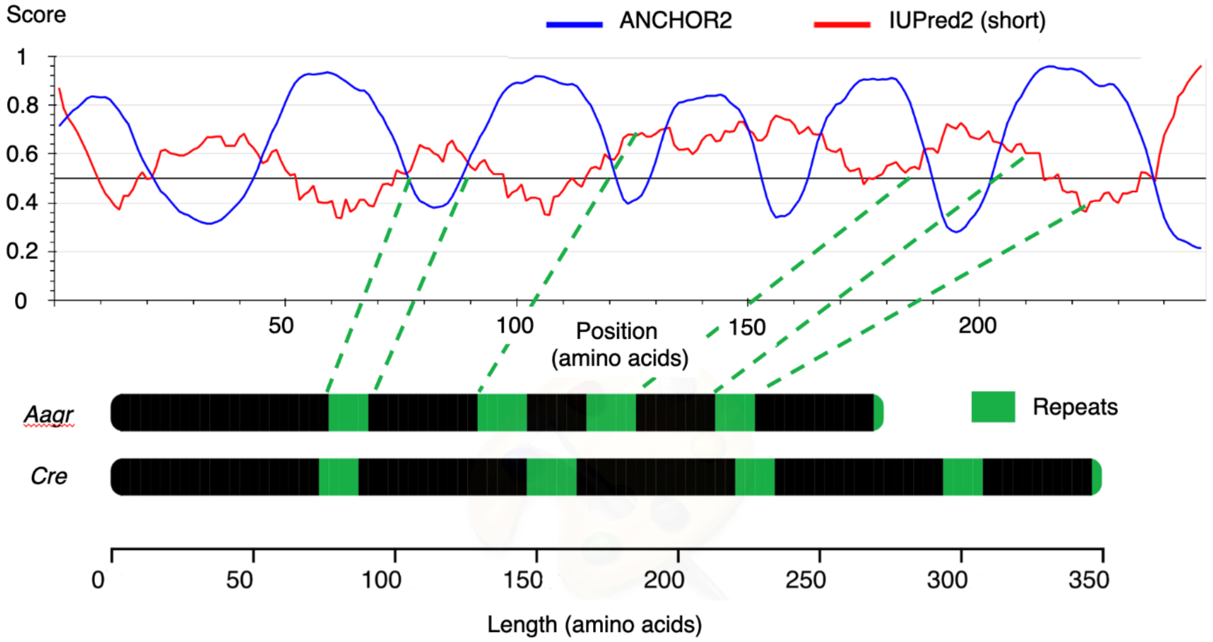


**Fig. S2. Intrinsically disordered profile of the EPYC1 candidate protein.**
The ANCHOR2 and IUPred2 scores have been obtained using the web interface IUPred2A. The IUPred2 score indicates the level of the intrinsically disordered profile of an amino acid sequence while the ANCHOR2 score indicates if a region is predicted to be involved in protein-protein interactions. The black and green bars show the protein sequences of EPYC1 in *Chlamydomonas reinhardtii* and the EPYC1 candidate protein in *Anthoceros agrestis*, the green blocks indicate the positions of the repeats.

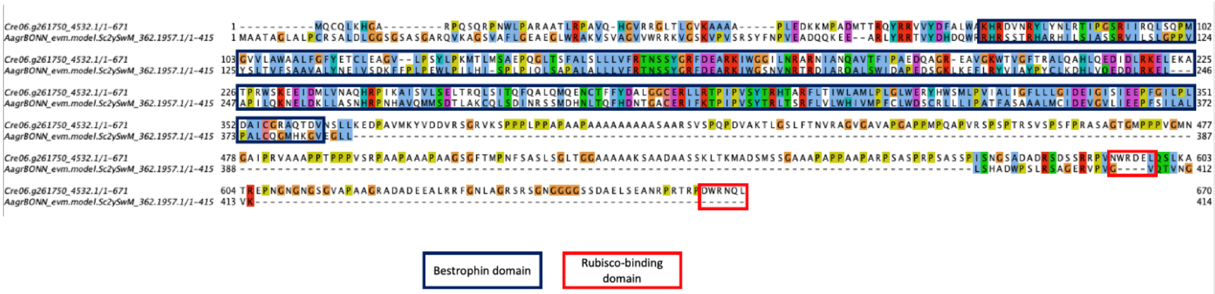


**Fig. S3. Alignment of RBMP1 protein sequences: *Chlamydomonas reinhardtii* versus *Anthoceros agrestis* BONN.**

**Table S3. Plasmid maps, sequences and their notations.** Plasmid constructs that were used to create stable transformed lines of *A. agrestis* [BONN].

| *A. agrestis* [BONN] gene ID | *C. reinhardtii* gene symbol | Construct description | Plasmid map |
|---|---|---|---|
| AagrBONN_evm.model.Sc2ySwM_228.417.1 | EPYC1 | *p-AaTip1::EPYC1-eGFP* and *p-AaTip1::EPYC1-eGFP* -p-AaTip1::*RSSU-mScarlet* | SR_L2_Tip1_EPYC_eGFP-sequence and SR_L2_Tip1_EPYC_eGFP_RSSU_mScarlet-sequence.pdf |
| AagrBONN_evm.model.Sc2ySwM_368.2612.1 | LCIB | *p-AaTip1::LCIB-mVenus* | SR_L2_Tip1_LCIB_mVenus-sequence.pdf |

| | | | |
|---|---|---|---|
| AagrBONN_evm.model.Sc2ySwM_117.2215.1 | HLA3 | *p-RSSU::HLA3-eGFP* | SR_L2_pRSSU_HLA_GFP-sequence.pdf |
| AagrBONN_evm.model.Sc2ySwM_344.4293.1 | CAH3 | *p-RSSU::CAH3-eGFP* | SR_L2_pRSSU_CAH_GFP-sequence.pdf |
| AagrBONN_evm.model.Sc2ySwM_344.2836.1 | RSSU | *p-AaTip1::RSSU-eGFP* | SR_L2_Tip1_RSSU_eGFP-sequence.pdf |
| AagrBONN_evm.model.Sc2ySwM_362.1957.1 | RBMP1 | *p-RSSU::RBMP1-eGFP* | SR_L2_pRSSU_RBMP_eGFP-sequence.pdf |

**Note S1: Plasmid maps and sequences**

10/01/2023 13:50:07

SR_L2_Tip1_EPYC_eGFP_RSSU_mScarlet (16759 bp)

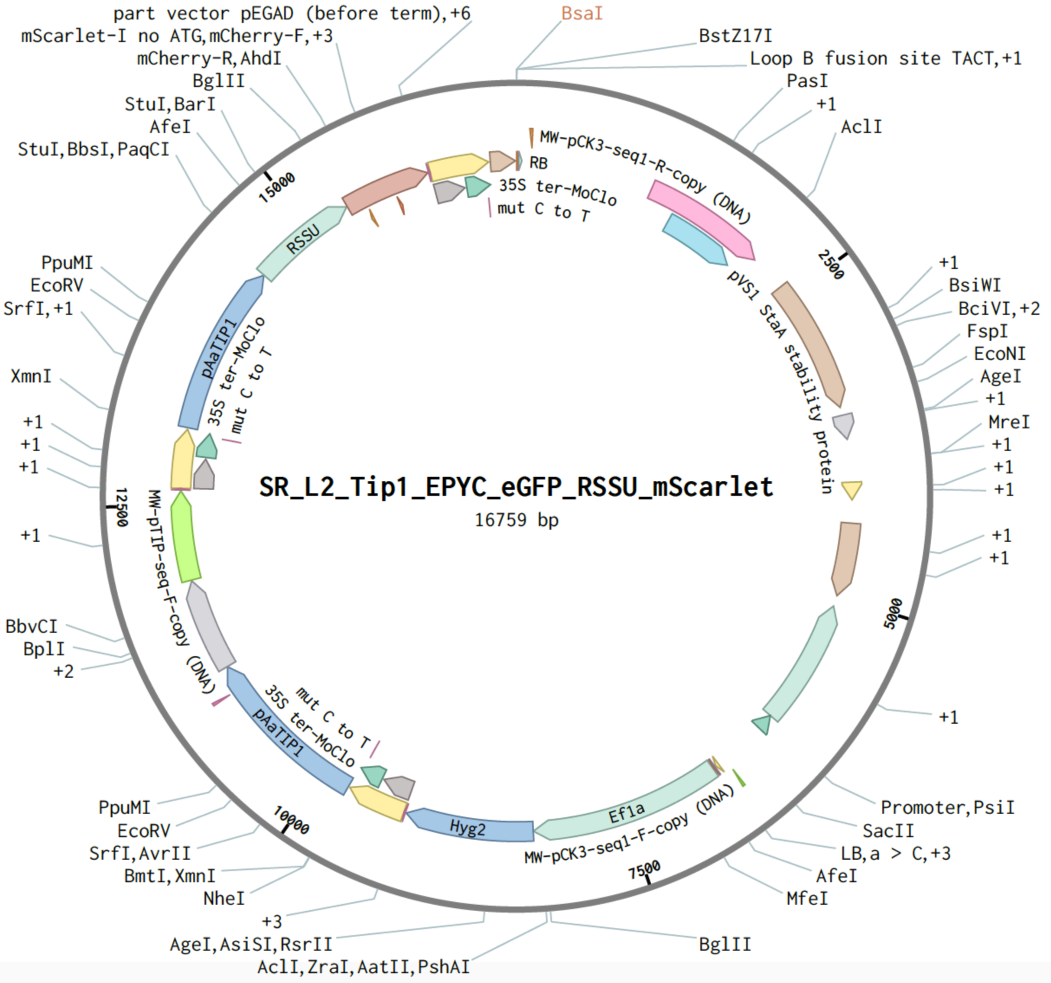

10/01/2023 13:50:21

### SR_L2_Tip1_RSSU_eGFP (13653 bp)

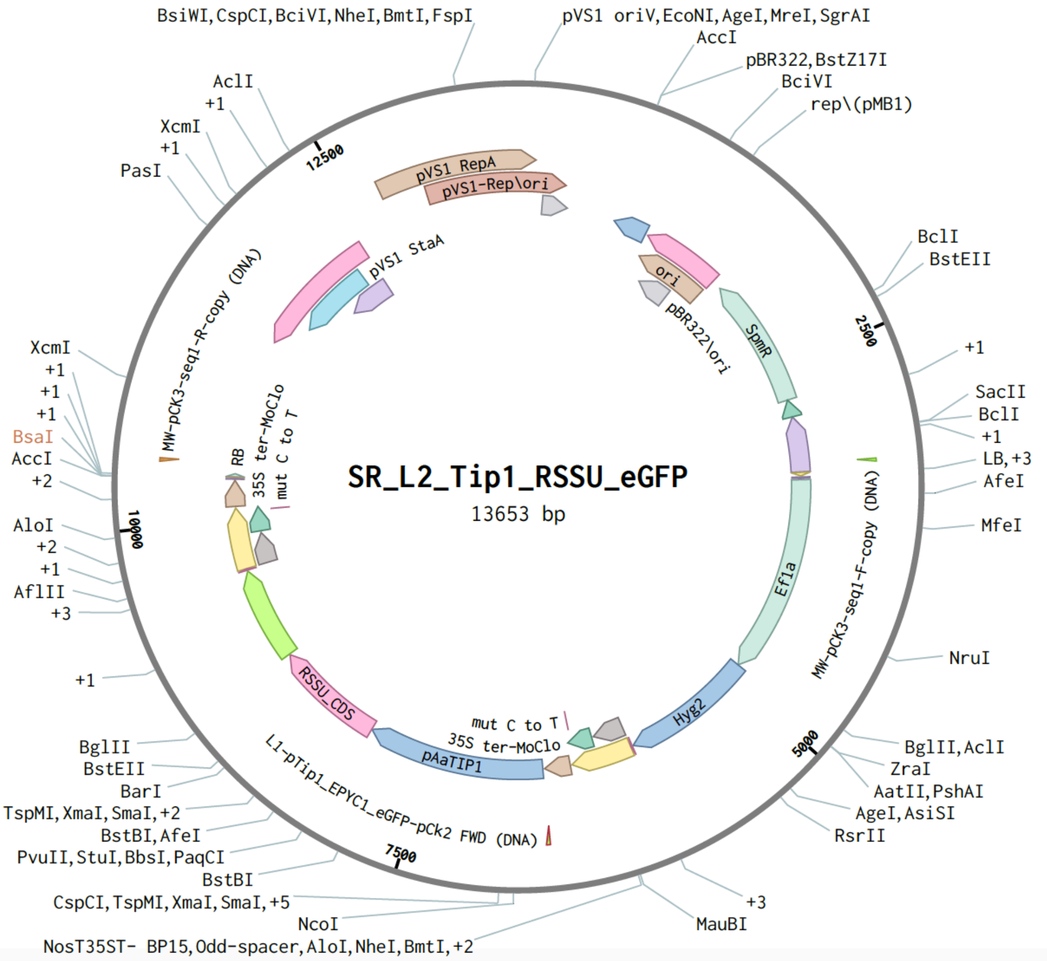

10/01/2023 13:50:14

### SR_L2_Tip1_LCIB_mVenus (14067 bp)

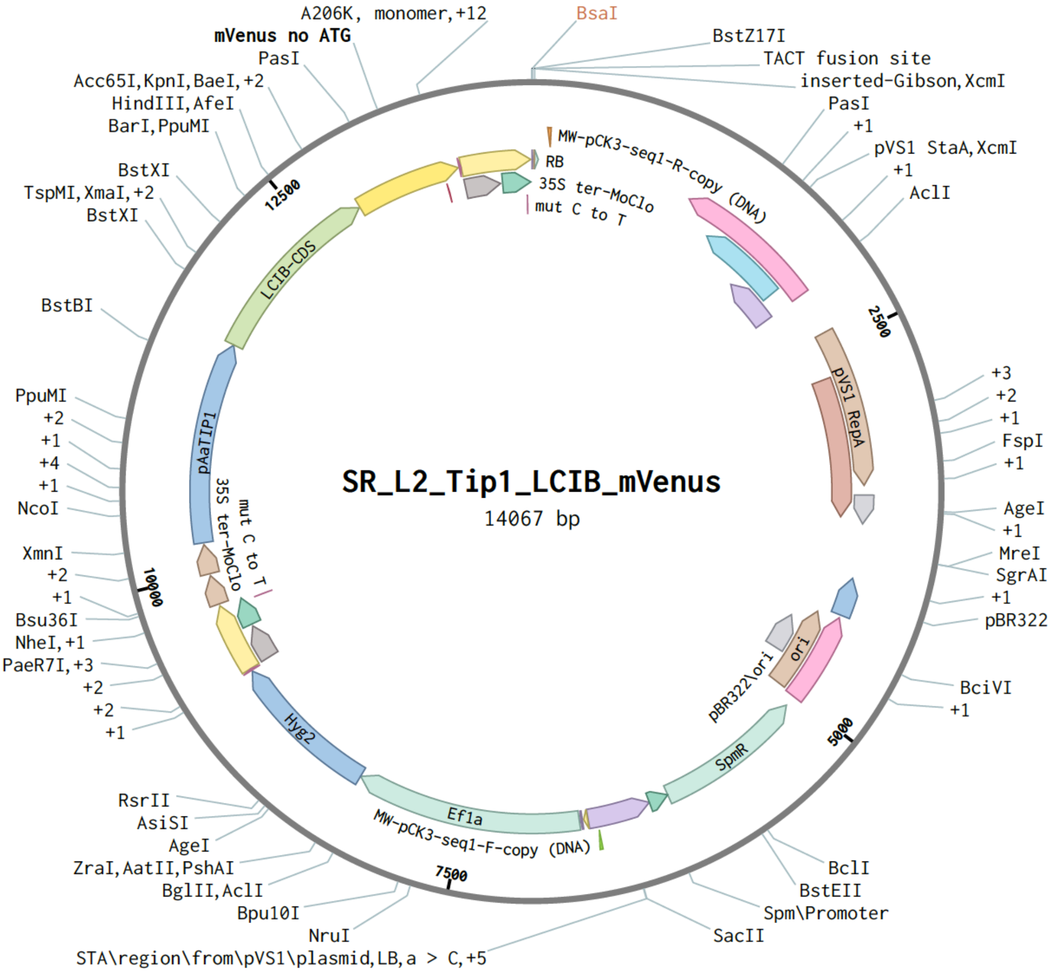

30/01/2023 13:31:48

### SR_L2_pRSSU_CAH_GFP (13867 bp)

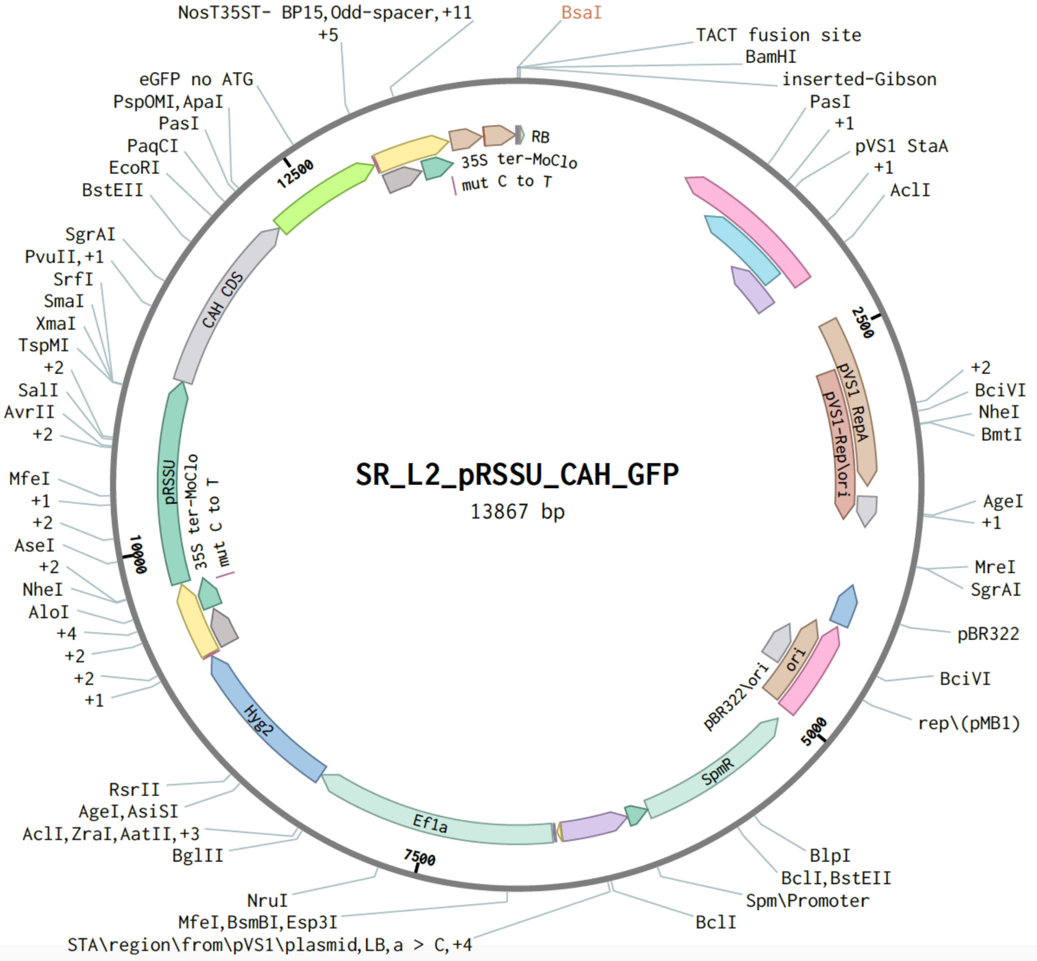

20/03/2024 15:56:32

### SR_L2_pRSSU_RBMP_GFP (13954 bp)

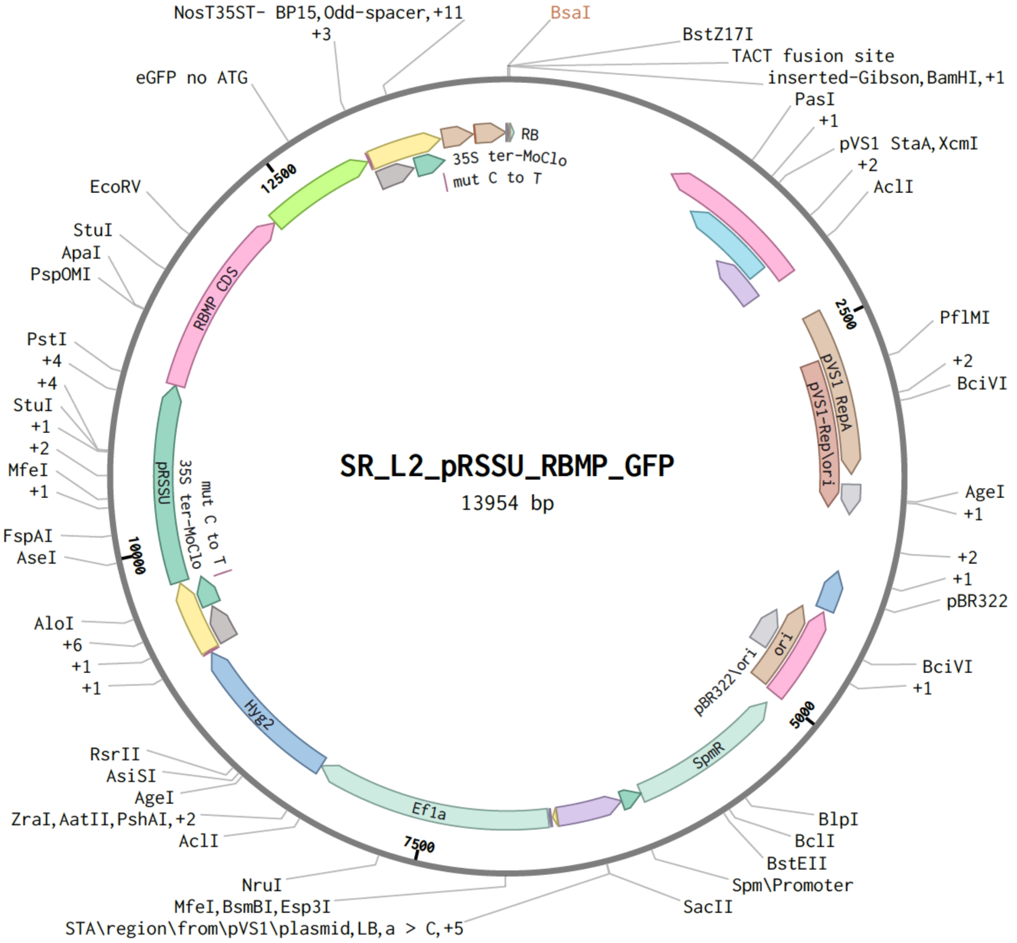


#### >promoter_RSSU

GGAGAGTTCATCTGTCAGCACTTGGACAGACCTTCCGGATCAAAGGTCGCTATTGCTATTCGCAGATCTAAAGTTTGAACGGCATTCACCGAGTGG
TGGATGCTTCAACTGGCTTCTGGCACCGAGCGTAAAAGGACAAATCCAACGTACATATCGCTAAATTTCGTTCCAACGTACATATCGCTAATTTTTC
CTACGGGAGGAATTAATCCTCGATAAACTCGGTGCCTTCTCGAAAGATGGTTTCACCATGGTTCAACTCCATGGCTCACCCCGACCCCACGCTTATC
CCACGGCTTCCAACACTCAGGGTGCTCGACTCCTATTACTCCCCGACCGTCTTCTTCGGAAAATCTTTATGCGCATTCCCCATCCTAACTTGGAGAT
GCGATTGCTTCATCCGCCTTCCTGTTCATCACTTCTCATCACCCCTCATTCGCAAAGAAATATCGTGGCACCTGGGATATGGAAATTTGCCCACTCT
GCCCTAGACTCTGAAAATCAGGGATGAACAAAAGGAATGACCGACAATCTTTGTGGAAAAGCTCTGGACAGGACTAGACTACCAATTGTAACCTGA
AACCTTGCTCTTTGAATGGGGAAATTCCTTACATGAGTCCTAATTGAGAAATCTGTCGCCCTACAAAGTTGCAAATGGCAGCTCTGGCTATGGCAG
CTCTAGCAATGGCCAGGCAAATGGCAGCTCTAGCAATGGCCAGGCAAATGGCAGCTCTAGCTATGGCCAGGCGGGCAAGGCCGTGATCTTCTTCCA
GAGCGAGCTCGGGCATACAGGCGGGGCCAGCATAGAGGGCTCACTATTTCCTCCAGCTGGGAGGCCTAGGGAAGGAAGGTCTGTGCCGCTGGCTAT
GGCGGTCGACTTGTGTGGGCCCTCCGAAGAACGAGCCAAAACTCTGAGCCAGTCGTTGCGCAGCCAATGAGCGGAGCTGTGGATGCTTGAACTGCG
TTTTGTCATTCGATTATGTATTGATTAGCAACTCAAATGATAAGTCGATCTCTCAGGTAATACATAGTCCATGGTTTGAGTTCAAATTAGTCGCTCA
AAAAACGAGGCTTTGGAGAAATCCAGCGAGGTTGGCGATCCTTTCTTATCCTTCCCTGCCAACTCTCTTCGTGTGGCCGTGAACGCTCTCTCGTTG
CCCCTATCCTATTTGAGCCCGGGCGGGCCACTCTCCAATCCAGTCCGCCCTCTCCCGGCCCTCTCCTTTTTCTTCCCCGGTAGCAGGCATACATCTTG
CGCTTCTCCGGTTCCATCCACACTGCAGCAGCA

#### >promoter_Aa_Tip1:1

CACCTGGAAGCATTTCCACACAGCATCTACACCACTACAGGGCTACATGCTCCATAGTGTAGAGCAAACCCTAGTGCAGCTCAGATATTAGATAATG
CTGTCCTTCCCAGAAAGCAAAAGGAATCGACAACTCTGGAAGGCAGACAGCACCAGTTCTGTGCAGCTTCATCCCGAGCTGATGCTGTCCTTCCGA
GAAAGCAAAGGAATCGACACTCCATGGCCGACCGCACAAACACACCATAGATGCTGACCTATCTGGATATGCAACAACCACAACACTTGCGTCTCC
ACCGCTCCAAGTCCAGCCTCCAGGAGCCAGGACTCACGGCCGGGTCGAGTGGGCCCAGCCTGGTCCCGTGGCGCCGGGGCCCGGGCCCGGCTCCCCC
CACTCCCCGGCGCGATTCTTACGCGGCGGACGGATAGGCCCACGACCGTCCTACGTGGCGGGCCCCACGTAAGGACAGCGGTGCCTAGGCTGCACCA
AATGCCGCGCCACCGTCCATGTCGTCAGAATCGGTCAGATATTAGCCCCTATCCCTTCACACGCCAGCGCCACGCCACGCCACGCTGCTGACGCCACC
CCCCTCTCACCATCATCATAGCCTTTCCAGATATCTCCACGCTCCTGCATTACCTGTACATCATTTTAATATGTGCATACATATGTATAATCAATCAGT
AAAGGCAAGGCAGGCTGGATCCCCAGTAGAGCCCATCAATGCCTTGAATCTCGTGTTTCCCGTCAGGTCCTTTCATTCATTCCAAGTCTCCTTCTCA

TACATACATCCCTCCGTGATTTAAACTTTATTAAATACCTACATAGCAATGGAATAAATCTATAATGTGGAAGAGACTCCAAGGTGTGACTGAGGTT
CCACATGTCTGAAAGAAGGAGCATCAAAGTATCTACCTTCCCTTCCATTTGGTTGTATTCCCTTCATCCCCTAAACCCAATGCAAGTAGCAAAAGAG
GGACACGAGAAGGTAGTAGTGAAGGTATTGCATTATTGATTTGTGAAAGCGTTTGATGAAACACAAAGGGCACACAAGAAGCGAATGAATGGATC
CACACATGGAGGGGCAGGCAGCTCGAAAAACAGCAGAGGATGCACATGGCCCAGAGCGGATAGGAAAGCAGGCCAGCCTCACTGGACAGTGGACA
GCCAGGTGCAGCGCACCCTGGCATATTCGAACCGGAAGGGCTGACGTGGGCTCCATTATTTGGGGTGCTGAGTTGGCAAACACGCGCCGGGAAGCT
CTCAAAACCGGCATCTCCATCTCATTCCAACGCCGCAGTCCCATTGCTCACCAACCCCGGCTCCTTTCCCCGCTCTGCTCCTCGGTTTCACGCTCCG
CGCGAGCTCTGCTG

#### >SR_L2_Tip1_EPYC1_eGFP_RSSU_mScarlet

ATACTTGAGACCGGATCCTGACAGGATATATTGGCGGGTAAACCTAAGAGAAAAGAGCGTTTATTAGAATAATCGGATATTTAAAAGGGCGTGAAA
AGGTTTATCCGTTCGTCCATTTGTATGTGCATGCCAACCACAGGGTTCCCCTCGGGATCAAAGTACTTTGATCCAACCCCTCCGCTGCTATAGTGCA
GTCGGCTTCTGACGTTCAGTGCAGCCGTCATCTGAAAACGACATGTCGCACAAGTCCTAAGTTACGCGACAGGCTGCCGCCCTGCCCTTTTCCTGGC
GTTTTCTTGTCGCGTGTTTTAGTCGCATAAAGTAGAATACTTGCGACTAGAACCGGAGACATTACGCCATGAACAAGAGCGCCGCCGCTGGCCTGC
TGGGCTATGCCCGCGTCAGCACCGACGACCAGGACTTGACCAACCAACGGGCCGAACTGCACGCGGCCGGCTGCACCAAGCTGTTTTCCGAGAAGA
TCACCGGCACCAGGCGCGACCGCCCGGAGCTGGCCAGGATGCTTGACCACCTACGCCCTGGCGACGTTGTGACAGTGACCAGGCTAGACCGCCTGGC
CCGCAGCACCCGCGACCTACTGGACATTGCCGAGCGCATCCAGGAGGCCGGCGCGGGCCTGCGTAGCCTGGCAGAGCCGTGGGCCGACACCACCACG
CCGGCCGGCCGCATGGTGTTGACCGTGTTCGCCGGCATTGCCGAGTTCGAGCGTTCCCTAATCATCGACCGCACCCGGAGCGGGCGCGAGGCCGCCA
AGGCCCGAGGCGTGAAGTTTGGCCCCCGCCCTACCCTCACCCCGGCACAGATCGCGCACGCCCGCGAGCTGATCGACCAGGAAGGCCGCACCGTGAA
AGAGGCGGCTGCACTGCTTGGCGTGCATCGCTCGACCCTGTACCGCGCACTTGAGCGCAGCGAGGAAGTGACGCCCACCGAGGCCAGGCGGCGCGG
TGCCTTCCGTGAGGACGCATTGACCGAGGCCGACGCCCTGGCGGCCGCCGAGAATGAACGCCAAGAGGAACAAGCATGAAACCGCACCAGGACGGC
CAGGACGAACCGTTTTTCATTACCGAAGAGATCGAGGCGGAGATGATCGCGGCCGGGTACGTGTTCGAGCCGCCCGCGCACCTCTCAACCGTGCGG
CTGCATGAAATCCTGGCCGGTTTGTCTGATGCCAAGCTGGCGGCCTGGCCGGCCAGCTTGGCCGCTGAAGAAACCGAGCGCCGCCGTCTAAAAAGG
TGATGTGTATTTGAGTAAAACAGCTTGCGTCATGCGGTCGCTGCGTATATGATGCGATGAGTAAATAAACAAATACGCAAGGGGAACGCATGAAGG
TTATCGCTGTACTTAACCAGAAAGGCGGGTCAGGCAAGACGACCATCGCAACCCATCTAGCCCGCGCCCTGCAACTCGCCGGGGCCGATGTTCTGTT
AGTCGATTCCGATCCCCAGGGCAGTGCCCGCGATTGGGCGGCCGTGCGGGAAGATCAACCGCTAACCGTTGTCGGCATCGACCGCCCGACGATTGAC
CGCGACGTGAAGGCCATCGGCCGGCGCGACTTCGTAGTGATCGACGGAGCGCCCCAGGCGGCGGACTTGGCTGTGTCCGCGATCAAGGCAGCCGAC
TTCGTGCTGATTCCGGTGCAGCCAAGCCCTTACGACATATGGGCCACCGCCGACCTGGTGGAGCTGGTTAAGCAGCGCATTGAGGTCACGGATGGA
AGGCTACAAGCGGCCTTTGTCGTGTCGCGGGCGATCAAAGGCACGCGCATCGGCGGTGAGGTTGCCGAGGCGCTGGCCGGGTACGAGCTGCCCATT
CTTGAGTCCCGTATCACGCAGCGCGTGAGCTACCCAGGCACTGCCGCCGCCGGCACAACCGTTCTTGAATCAGAACCCGAGGGCGACGCTGCCCGCG
AGGTCCAGGCGCTGGCCGCTGAAATTAAATCAAAACTCATTTGAGTTAATGAGGTAAAGAGAAAATGAGCAAAAGCACAAACACGCTAAGTGCCG
GCCGTCCGAGCGCACGCAGCAGCAAGGCTGCAACGTTGGCCAGCCTGGCAGACACGCCAGCCATGAAGCGGGTCAACTTTCAGTTGCCGGCGGAGG
ATCACACCAAGCTGAAGATGTACGCGGTACGCCAAGGCAAGACCATTACCGAGCTGCTATCTGAATACATCGCGCAGCTACCAGAGTAAATGAGCAA
ATGAATAAATGAGTAGATGAATTTTAGCGGCTAAAGGAGGCGGCATGGAAAATCAAGAACAACCAGGCACCGACGCCGTGGAATGCCCCATGTGTG
GAGGAACGGGCGGTTGGCCAGGCGTAAGCGGCTGGGTTGCCTGCCGGCCCTGCAATGGCACTGGAACCCCCAAGCCCGAGGAATCGGCGTGAGCGG
TCGCAAACCATCCGGCCCGGTACAAATCGGCGCGGCGCTGGGTGATGACCTGGTGGAGAAGTTGAAGGCCGCGCAGGCCGCCCAGCGGCAACGCAT
CGAGGCAGAAGCACGCCCCGGTGAATCGTGGCAAGCGGCCGCTGATCGAATCCGCAAAGAATCCCGGCAACCGCCGGCAGCCGGTGCGCCGTCGAT
TAGGAAGCCGCCCAAGGGCGACGAGCAACCAGATTTTTTCGTTCCGATGCTCTATGACGTGGGCACCCGCGATAGTCGCAGCATCATGGACGTGGC
CGTTTTCCGTCTGTCGAAGCGTGACCGACGAGCTGGCGAGGTGATCCGCTACGAGCTTCCAGACGGGCACGTAGAGGTTTCCGCAGGGCCGGCCGG
CATGGCCAGTGTGTGGGATTACGACCTGGTACTGATGGCGGTTTCCCATCTAACCGAATCCATGAACCGATACCGGGAAGGGAAGGGAGACAAGCC
CGGCCGCGTGTTCCGTCCACACGTTGCGGACGTACTCAAGTTCTGCCGGCGAGCCGATGGCGGAAAGCAGAAAGACGACCTGGTAGAAACCTGCAT
TCGGTTAAACACCACGCACGTTGCCATGCAGCGTACGAAGAAGGCCAAGAACGGCCGCCTGGTGACGGTATCCGAGGGTGAAGCCTTGATTAGCCG
CTACAAGATCGTAAAGAGCGAAACCGGGCGGCCGGAGTACATCGAGATCGAGCTAGCTGATTGGATGTACCGCGAGATCACAGAAGGCAAGAACCC
GGACGTGCTGACGGTTCACCCCGATTACTTTTTGATCGATCCCGGCATCGGCCGTTTTCTCTACCGCCTGGCACGCCGCGCCGCAGGCAAGGCAGAA
GCCAGATGGTTGTTCAAGACGATCTACGAACGCAGTGGCAGCGCCGGAGAGTTCAAGAAGTTCTGTTTCACCGTGCGCAAGCTGATCGGGTCAAAT
GACCTGCCGGAGTACGATTTGAAGGAGGAGGCGGGGCAGGCTGGCCCGATCCTAGTCATGCGCTACCGCAACCTGATCGAGGGCGAAGCATCCGCC
GGTTCCTAATGTACGGAGCAGATGCTAGGGCAAATTGCCCTAGCAGGGGAAAAAGGTCGAAAAGGACTCTTTCCTGTGGATAGCACGTACATTGGG
AACCCAAAGCCGTACATTGGGAACCGGAACCCGTACATTGGGAACCCAAAGCCGTACATTGGGAACCGGTCACACATGTAAGTGACTGATATAAAA
GAGAAAAAAGGCGATTTTTCCGCCTAAAACTCTTTAAAACTTATTAAAACTCTTAAAACCCGCCTGGCCTGTGCATAACTGTCTGGCCAGCGCACA
GCCGAAGTGCTGCAAAAAGCGCCTACCCTTCGGTCGCTGCGCTCCCTACGCCCCGCCGCTTCGCGTCGGCCTATCGCGGCCGCTGGCCGCTCAAAAA
TGGCTGGCCTACGGCCAGGCAATCTACCAGGGCGCGGACAAGCCGCGCCGTCGCCACTCGACCGCCGGCGCCCACATCAAGGCACCCTGCCTCGCGC
GTTTCGGTGATGACGGTGAAAACCTCTGACACATGCAGCTCCCGGTGACGGTCACAGCTTGTCTGTAAGCGGATGCCGGGAGCAGACAAGCCCGTC
AGGGCGCGTCAGCGGGTGTTGGCGGGTGTCGGGGCGCAGCCATGACCCAGTCACGTAGCGATAGCGGAGTGTATACTGGCTTAACTATGCGGCATC
AGAGCAGATTGTACTGAGAGTGCACCATATGCGGTGTGAAATACCGCACAGATGCGTAAGGAGAAAATACCGCATCAGGCGCTCATCCGCTTCCTCG
CTCACTGACTCGCTGCGCTCGGTCGTTCGGCTGCGGCGAGCGGTATCAGCTCACTCAAAGGCGGTAATACGGTTATCCACAGAATCAGGGGATAACG
CAGGAAAGAACATGTGAGCAAAAGGCCAGCAAAAGGCCAGGAACCGTAAAAAGGCCGCGTTGCTGGCGTTTTTCCATAGGCTCCGCCCCCCTGAC
GAGCATCACAAAAATCGACGCTCAAGTCAGAGGTGGCGAAACCCGACAGGACTATAAAGATACCAGGCGTTTCCCCCTGGAAGCTCCCTCGTGCGC
TCTCCTGTTCCGACCCTGCCGCTTACCGGATACCTGTCCGCCTTTCTCCCTTCGGGAAGCGTGGCGCTTTCTCATAGCTCACGCTGTAGGTATCTCAG
TTCGGTGTAGGTCGTTCGCTCCAAGCTGGGCTGTGTGCACGAACCCCCCGTTCAGCCCGACCGCTGCGCCTTATCCGGTAACTATCGTCTTGAGTCC
AACCCGGTAAGACACGACTTATCGCCACTGGCAGCAGCCACTGGTAACAGGATTAGCAGAGCGAGGTATGTAGGCGGTGCTACAGAGTTCTTGAAG
TGGTGGCCTAACTACGGCTACACTAGAAGGACAGTATTTGGTATCTGCGCTCTGCTGAAGCCAGTTACCTTCGGAAAAAGAGTTGGTAGCTCTTGA
TCCGGCAAACAAACCACCGCTGGTAGCGGTGGTTTTTTTGTTTGCAAGCAGCAGATTACGCGCAGAAAAAAAGGATCTCAAGAAGATCCTTTGAT
CTTTTCTACGGGGTCTGACGCTCAGTGGAACGAAAACTCACGTTAAGGGATTTTGGTCATGCATTCTAGGTGATTATTTGCCGACTACCTTGGTGA
TCTCGCCTTTCACGTAGTGGACAAATTCTTCCAACTGATCTGCGCGCGAGGCCAAGCGATCTTCTTCTTGTCCAAGATAAGCCTGTCTAGCTTCAAG
TATGACGGGCTGATACTGGGCCGGCAGGCGCTCCATTGCCCAGTCGGCAGCGACATCCTTCGGCGCGATTTTGCCGGTTACTGCGCTGTACCAAATG
CGGGACAACGTAAGCACTACATTTCGCTCATCACCAGCCCAGTCGGGCGGCGAGTTCCATAGCGTTAAGGTTTCATTTAGCGCCTCAAATAGATCCT
GTTCAGGAACCGGATCAAAGAGTTCCTCCGCCGCTGGACCTACCAAGGCAACGCTATGTTCTCTTGCTTTTGTCAGCAAGATAGCCAGATCAATGT
CGATCGTGGCTGGCTCGAAGATACCTGCAAGAATGTCATTGCGCTGCCATTCTCCAAATTGCAGTTCGCGCTTAGCTGGATAACGCCACGGAATGAT
GTCGTCGTGCACAACAATGGTGACTTCTACAGCGCGGAGAATCTCGCTCTCTCCAGGGGAAGCCGAAGTTTCCAAAAGGTCGTTGATCAAAGCTCG
CCGCGTTGTTTCATCAAGCCTTACGGTCACCGTAACCAGCAAATCAATATCACTGTGTGGCTTCAGGCCGCCATCCACTGCGGAGCCGTACAAATGT
ACGGCCAGCAACGTCGGTTCGAGATGGCGCTCGATGACGCCAACTACCTCTGATAGTTGAGTCGATACTTCGGCGATCACCGCTTCCCTCATAATGT
TTAACTTTGTTTTAGGGCGACTGCCCTGCTGCGTAACATCGTTGCTGCTCCATAACATCAAACATCGACCCACGGCGTAACGCGCTTGCTGCTTGGA
TGCCCGAGGCATAGACTGTACCCCAAAAAAACAGTCATAACAAGCCATGAAAACCGCCACTGCGCCGTTACCACCGCTGCGTTCGGTCAAGGTTCT
GGACCAGTTGCGTGAGCGCATACGCTACTTGCATTACAGCTTACGAACCGAACAGGCTTATGTCCACTGGGTTCGTGCCTTCATCCGTTTCCACGGT
GTGCGTCACCCGGCAACCTTGGGTAGCAGCGAAGTCGAGGCATTTCTGTCCTGGCTGGAACAGAACTTATTATTTCCTTCCTCTTTTCTACAGTATT

TAAAGATACCCCAAGAAGCTAATTATAACAAGACGAACTCCAATTCACTGTTCCTTGCATTCTAAAACCTTAAATACCAGAAAACAGCTTTTTCAA
AGTTGTTTTCAAAGTTGGCGTATAACATAGTATCGACGGAGCCGATTTTGAAACCGCGGTGATCACAGGCAGCAACGCTCTGTCATCGTTACAATC
AACATGCTACCCTCCGCGAGATCATCCGTGTTTCAAACCCGGCAGCTTAGTTGCCGTTCTTCCGAATAGCATCGGTAACATGAGCAAAGTCTGCCGC
CTTACAACGGCTCTCCCGCTGACGCCGTCCCGGACTGATGGGCTGCCTGTATCGAGTGGTGATTTTGTGCCGAGCTGCCGGTCGGGGAGCTGTTGG
CTGGCTGGTGGCAGGATATATTGTGGTGTAAACATAACGGATCCGGTCTCAGGAGAATGAGGAGCACCGTCCAAACCTGGATAGCCGATATGTGAG
TGTTTGGTTGCCTGTCATTTCTAAAAAGCGTATGTCTCGTAGCGTAGATAGCAAGCGCTGTGAGAAAGGTGTGCTCCTATAGCATGGTAGCATCTGT
CCCATATCTCGTCCGCGAGAAGCCTACATTTCTCGTTGCGTATTAGAGCATGAAAAACCAACGTGAAGTACGTGATTCAGACCTTGACCAGTTCCAG
TTCTCTTCAACAAAAAAACAGTTCTTCAATTCATTTGCTTACATTAAACAATTGTCACACTCTTCTTCCCGGATGACAGCACCAAGACACTTCCCC
TCCACAAAAATCTCCGAGTCATGTGGACCGGCCTGGATGCTCTTAAAGGGGTTGCCATTGTAGCGGGGTGCCTTAACCACGACCTCTTCGATACCTG
TCTGAAGCGGCGTGTGAAATCCTCCAGGAGCTTGTTCAAATTTACAAGCTGCAATGGATGTAAACTCATCCAAAGACATGACTTGCGAGAAGCCAG
AAACGGTCATCAACACTACGAACACCCCTAAACATGGATGGAGAGGCAACATCCTGCATGTGTCTATGTCAGAGGATGCAAAAGTGAAAGACAAAG
GAGAAGGATTGCTCGTGCAATCACTGCAAAGCAAGATGCGAATATCCCTAAAATCCACAGCAGGAAAAGAAAGGAGCCTCAATAAGTCTCTAAAAT
GGGAAGAAAAGAAGCACGGGCCAGCGAGACGGAAATGAGGCGATCCGATGCGATGGTTCCGAGAACTGAAGCGGTGCGTGGGGGGCTAGAAATGA
ATGGGGAACGTGGAGGGCAAACAAAGGCTGTACTTAGCGGCAAATGCAGATGGCATGCATGCCAAGAACAGCAGCAGTCGGCACCGCCAGTTTAG
CGGAGAGCAACAGGCAGCAAAAGGAAGTGGAACAGGTTTAGGGCCGACGCAGTCGGCGCCTGCCCTAAAACGGCAAGGCCCACGGCAGCGGTCGC
GAGGGCGTTGCCTGCCGGTCGTGGAAAGCCGTGGGGCTGTCAACGGAACCTCACGCTCCCAGCCTGCCCTAAGCGGGGAAAAAGAAGATCCACATT
CTAGGGCTTTCCATCTCGGCTCTCAGCCGTCCGATTAAGAATCGCACGGCCTAGCCCCGCCAGCCTAGCCCTCGGCCGCCCCCTTGACAGGTTAAGA
GCTGGGCAAGCGACTCAGTTGTTCCTTCGTCGCTCGCCCTTCTCACGAGCAGCTCGCCTGCCCCGCCGCCGCCTTGCCTGCCTCGCGCCTTTTGTCT
CCAGGTTCCCTTGCCGCGCCTTCTCCTTGGCTTGTGCTTCTGCTTTGCTTGTGCCTCTGCTGGGCTCTACTTGTGTGGTGTTGTGCTTTGGCGCCGC
CGCCTCGCTGCTGCTGTGGAGCTTGGTGTGTGCTGGGGTTTCTGGGGTTTGCATGTGTCAGGGTTTGTAGATCTGTTCTTCGGGGCCGTCTTTCTA
ACGTTTCGTGTGCCGTTGTTTCTGGGGGGTTTGGTGTGGAGCTCGCAGCTTGCAGCTAATGAAAAAGCCTGAACTCACCGCGACGTCTGTCGAGAA
GTTTCTGATCGAAAAGTTCGACAGCGTCTCCGACCTGATGCAGCTCTCGGAGGGCGAAGAATCTCGTGCTTTCAGCTTCGATGTAGGAGGGCGTGG
ATATGTCCTGCGGGTAAATAGCTGCGCCGATGGTTTCTACAAAGATCGTTATGTTTATCGGCACTTTGCATCGGCCGCGCTCCCGATTCCGGAAGTG
CTTGACATTGGGGAGTTTAGCGAGAGCCTGACCTATTGCATCTCCCGCCGTTCACAGGGTGTCACGTTGCAAGACCTGCCTGAAACCGAACTGCCC
GCTGTTCTACAACCGGTCGCGGAGGCTATGGATGCGATCGCTGCGGCCGATCTTAGCCAGACGAGCGGGTTCGGCCCATTCGGACCGCAAGGAATC
GGTCAATACACTACATGGCGTGATTTCATATGCGCGATTGCTGATCCCCATGTGTATCACTGGCAAACTGTGATGGACGACACCGTCAGTGCGTCCG
TCGCGCAGGCTCTCGATGAGCTGATGCTTTGGGCCGAGGACTGCCCCGAAGTCCGGCACCTCGTGCACGCGGATTTCGGCTCCAACAATGTCCTGA
CGGACAATGGCCGCATAACAGCGGTCATTGACTGGAGCGAGGCGATGTTCGGGGATTCCCAATACGAGGTCGCCAACATCTTCTTCTGGAGGCCGT
GGTTGGCTTGTATGGAGCAGCAGACGCGCTACTTCGAGCGGAGGCATCCGGAGCTTGCAGGATCGCCACGACTCCGGGCGTATATGCTCCGCATTG
GTCTTGACCAACTCTATCAGAGCTTGGTTGACGGCAATTTCGATGATGCAGCTTGGGCGCAGGGTCGATGCGACGCAATCGTCCGATCCGGAGCCG
GGACTGTCGGGCGTACACAAATCGCCCGCAGAAGCGCGGCCGTCTGGACCGATGGCTGTGTAGAAGTACTCGCCGATAGTGGAAACCGACGCCCCA
GCACTCGTCCGAGGGCAAAGAAATAGGCTTGAGCTCGAATTTCCCCGATCGTTCAAACATTTGGCAATAAAGTTTCTTAAGATTGAATCCTGTTGC
CGGTCTTGCGATGATTATCATATAATTTCTGTTGAATTACGTTAAGCATGTAATAATTAACATGTAATGCATGACGTTATTTATGAGATGGGTTTTT
ATGATTAGAGTCCCGCAATTATACATTTAATACGCGATAGAAAACAAAATATAGCGCGCAAACTAGGATAAATTATCGCGCGCGGTGTCATCTATGT
TACTAGATCGATCCGTATCGATAGCCTCTAGCTAGAGTCGATCGACAAGCTCGAGTTTCTCCATAATAATGTGTGAGTAGTTCCCAGATAAGGGAAT
TAGGGTTCCTATAGGGTTTCGCTCATGTGTTGAGCATATAAGAAACCCTTAGTATGTATTTGTATTTGTAAAATACTTCTATCAATAAAATTTCTAAT
TCCTAAAATCAAAATCCAGTACTAAAATCCAGATCGCTAGCAAGGAGCACCTGGAAGCATTTCCACACAGCATCTACACCACTACAGGGCTACATGC
TCCATAGTGTAGAGCAAACCCTAGTGCAGCTCAGATATTAGATAATGCTGTCCTTCCCAGAAAGCAAAAGGAATCGACAACTCTGGAAGGCAGACA
GCACCAGTTCTGTGCAGCTTCATCCCGAGCTGATGCTGTCCTTCCGAGAAAGCAAAGGAATCGACACTCCATGGCCGACCGCACAAACACACCATA
GATGCTGACCTATCTGGATATGCAACAACCACAACACTTGCGTCTCCACCGCTCCAAGTCCAGCCTCCAGGAGCCAGGACTCACGGCCGGGTCGAGT
GGGCCCAGCCTGGTCCCGTGGCGCCGGGGCCCGGGCCCGGCTCCCCCACTCCCCGGCGCGATTCTTACGCGGCGGACGGATAGGCCCACGACCGTC
CTACGTGGCGGGCCCCACGTAAGGACAGCGGTGCCTAGGCTGCACCAAATGCCGCGCCACCGTCCATGTCGTCAGAATCGGTCAGATATTAGCCCCT
ATCCCTTCACACGCCAGCGCCACGCCACGCCACGCTGCTGACGCCACCCCCCTCTCACCATCATCATAGCCTTTCCAGATATCTCCACGCTCCTGCAT
TACCTGTACATCATTTTAATATGTGCATACATATGTATAATCAATCAGTAAAGGCAAGGCAGGCTGGATCCCCAGTAGAGCCCATCAATGCCTTGAA
TCTCGTGTTTCCCGTCAGGTCCTTTCATTCATTCCAAGTCTCCTTCTCATACATACATCCCTCCGTGATTTAAACTTTATTAAATACCTACATAGCAA
TGGAATAAATCTATAATGTGGAAGAGACTCCAAGGTGTGACTGAGGTTCCACATGTCTGAAAGAAGGAGCATCAAAGTATCTACCTTCCCTTCCAT
TTGGTTGTATTCCCTTCATCCCCTAAACCCAATGCAAGTAGCAAAAGAGGGACACGAGAAGGTAGTAGTGAAGGTATTGCATTATTGATTTGTGAA
AGCGTTTGATGAAACACAAAGGGCACACAAGAAGCGAATGAATGGATCCACACATGGAGGGGCAGGCAGCTCGAAAAACAGCAGAGGATGCACAT
GGCCCAGAGCGGATAGGAAAGCAGGCCAGCCTCACTGGACAGTGGACAGCCAGGTGCAGCGCACCCTGGCATATTCGAACCGGAAGGGCTGACGTG
GGCTCCATTATTTGGGGTGCTGAGTTGGCAAACACGCGCCGGGAAGCTCTCAAAACCGGCATCTCCATCTCATTCCAACGCCGCAGTCCCATTGCT
CACCAACCCCGGCTCCTTTCCCCGCTCTGCTCCTCGGTTTCACGCTCCGCGCGAGCTCTGCTGCAATGGAAAAGCTGACAGCGGTTATCCCGTCACT
GTTTAAGGAAGGAATGACCTCTAGCACGGGCATTACCGCATCCAGCAGTAAAGCCTCAACCCAGGAAACCTCTTCGAAATCCCGCCGCGATCCTGC
GTCGGCCACTGTCACCCGCCAACAACTGGCCATGGTTGAGCCAGACCTGGTGGCCAACTGGGACGATAACTGGCTCTCAAACCTGCAGCCATTACA
TAAGGATGAAACCTCTAGCAAGAGCCGTCGCGACCCGGCATCAGCGACGATTACTGGGGAACAGCTGGCTATGGTGAAGCCCGAATTGGTTGCAAA
CTGGGATGATAACTGGCTTTCAAACCTTCAGCCCCATGATAAAGATGAGACATCCTCAAAATCCCGTCGCGACCCCGCCTTAGCCACCATCACACCG
GAACAAGACAACTGGCTGTCAAACCTGCAGCCTCACGACAAAGATGAAACCTCAAGCAAATCGCGTCGCGATCCGGCCTCAGCCACTGTCACGCGC
CAACAGCTGGCCAACTGGGATGATAATTGGCTCTCTAACTTGCAGCGTCATCACAAAGACAAGGACGAGACTTCGAGCAAGAGCCGCCGCGACCCG
GCATCAGCCACGGTTACGCATCAACAGCTGGCAATGGTCAAACCGGAGCTGGCCGCCGATTGGGATGACAACTGGTTGTCATCTTTACAGCGTCAC
CACAAAGATAAAGACGAAACTTCTAGCAAAAGTCGTGCTTCGGTGAGCAAGGGCGAGGAGCTGTTCACCGGGGTGGTGCCCATCCTGGTCGAGCT
GGACGGCGACGTAAACGGCCACAAGTTCAGCGTGTCCGGCGAGGGCGAGGGCGATGCCACCTACGGCAAGCTGACCCTGAAGTTCATCTGCACCAC
CGGCAAGCTGCCCGTGCCCTGGCCCACCCTCGTGACCACCCTGACCTACGGCGTGCAGTGCTTCAGCCGCTACCCCGACCACATGAAGCAGCACGAC
TTCTTCAAGTCCGCCATGCCCGAAGGCTACGTCCAGGAGCGCACCATCTTCTTCAAGGACGACGGCAACTACAAGACCCGCGCCGAGGTGAAGTTC
GAGGGCGACACCCTGGTGAACCGCATCGAGCTGAAGGGCATCGACTTCAAGGAGGACGGCAACATCCTGGGGCACAAGCTGGAGTACAACTACAAC
AGCCACAACGTCTATATCATGGCCGACAAGCAGAAGAACGGCATCAAGGTGAACTTCAAGATCCGCCACAACATCGAGGACGGCAGCGTGCAGCTC
GCCGACCACTACCAGCAGAACACCCCCATCGGCGACGGCCCCGTGCTGCTGCCCGACAACCACTACCTGAGCACCCAGTCCGCCCTGAGCAAAGACC
CCAACGAGAAGCGCGATCACATGGTCCTGCTGGAGTTCGTGACCGCCGCCGGGATCACTCTCGGCATGGACGAGCTGTACAAGTAAGCTTGAGCTC
GAATTTCCCCGATCGTTCAAACATTTGGCAATAAAGTTTCTTAAGATTGAATCCTGTTGCCGGTCTTGCGATGATTATCATATAATTTCTGTTGAAT
TACGTTAAGCATGTAATAATTAACATGTAATGCATGACGTTATTTATGAGATGGGTTTTTATGATTAGAGTCCCGCAATTATACATTTAATACGCGAT
AGAAAACAAAATATAGCGCGCAAACTAGGATAAATTATCGCGCGCGGTGTCATCTATGTTACTAGATCGATCCGTATCGATAGCCTCTAGCTAGAGT
CGATCGACAAGCTCGAGTTTCTCCATAATAATGTGTGAGTAGTTCCCAGATAAGGGAATTAGGGTTCCTATAGGGTTTCGCTCATGTGTTGAGCATA
TAAGAAACCCTTAGTATGTATTTGTATTTGTAAAATACTTCTATCAATAAAATTTCTAATTCCTAAAATCAAAATCCAGTACTAAAATCCAGATCGC
TATACAGGAGCACCTGGAAGCATTTCCACACAGCATCTACACCACTACAGGGCTACATGCTCCATAGTGTAGAGCAAACCCTAGTGCAGCTCAGATA
TTAGATAATGCTGTCCTTCCCAGAAAGCAAAAGGAATCGACAACTCTGGAAGGCAGACAGCACCAGTTCTGTGCAGCTTCATCCCGAGCTGATGCT
GTCCTTCCGAGAAAGCAAAGGAATCGACACTCCATGGCCGACCGCACAAACACACCATAGATGCTGACCTATCTGGATATGCAACAACCACAACAC

TTGCGTCTCCACCGCTCCAAGTCCAGCCTCCAGGAGCCAGGACTCACGGCCGGGTCGAGTGGGCCCAGCCTGGTCCCGTGGCGCCGGGGCCCGGGC
CCGGCTCCCCCCACTCCCCGGCGCGATTCTTACGCGGCGGACGGATAGGCCCACGACCGTCCTACGTGGCGGGCCCCACGTAAGGACAGCGGTGCCT
AGGCTGCACCAAATGCCGCGCCACCGTCCATGTCGTCAGAATCGGTCAGATATTAGCCCCTATCCCTTCACACGCCAGCGCCACGCCACGCCACGCT
GCTGACGCCACCCCCCTCTCACCATCATCATAGCCTTTCCAGATATCTCCACGCTCCTGCATTACCTGTACATCATTTTAATATGTGCATACATATGTA
TAATCAATCAGTAAAGGCAAGGCAGGCTGGATCCCCAGTAGAGCCCATCAATGCCTTGAATCTCGTGTTTCCCGTCAGGTCCTTTCATTCATTCCAA
GTCTCCTTCTCATACATACATCCCTCCGTGATTTAAACTTTATTAAATACCTACATAGCAATGGAATAAATCTATAATGTGGAAGAGACTCCAAGGT
GTGACTGAGGTTCCACATGTCTGAAAGAAGGAGCATCAAAGTATCTACCTTCCCTTCCATTTGGTTGTATTCCCTTCATCCCCTAAACCCAATGCAA
GTAGCAAAAGAGGGACACGAGAAGGTAGTAGTGAAGGTATTGCATTATTGATTTGTGAAAGCGTTTGATGAAACACAAAGGGCACACAAGAAGCG
AATGAATGGATCCACACATGGAGGGGCAGGCAGCTCGAAAAACAGCAGAGGATGCACATGGCCCAGAGCGGATAGGAAAGCAGGCCAGCCTCACT
GGACAGTGGACAGCCAGGTGCAGCGCACCCTGGCATATTCGAACCGGAAGGGCTGACGTGGGCTCCATTATTTGGGGTGCTGAGTTGGCAAACACG
CGCCGGGAAGCTCTCAAAACCGGCATCTCCATCTCATTCCAACGCCGCAGTCCCATTGCTCACCAACCCCGGCTCCTTTCCCCGCTCTGCTCCTCGG
TTTCACGCTCCGCGCGAGCTCTGCTGCAATGGCTTCCGTGGTCGCTTCCTCCGCCGTCGTGGCCCCAGCTGCATTCTCTGCCGCTTCCTCCAGCGGC
TCCGCCTCTGAGGCCTCCAGCATGAGGGCCTTCGTCGGCCTCAAGTCCGGCGATGTCTTCTCCAGCAAGACCGCCGACTTGAGCTCCAGGACCGTCA
GCAACGGGAGCCGCGTGAATTGCATGCAGGTGTGGACTCCGGTGGACAACAAGAAGTTCGAAACCCTCTCGTACTTGCCCGCGCTCAGCTCCAATC
AAATCGCTAAGCAAATCGACTTCATGTTGTCGAAGGGCTGGATCCCCTGCCTCGAGTTCGACAAGAGCGCTGAGCTCAAAGTGGTGCGAGGCCAGC
CCGGGTTCTACAACGGCCGGTACTGGACAATGTGGAAGCTTCCCATGTTCGGCTGCAACGACGCTTCATCCGTGTTGAGGGAAGTTGAGAACTGCA
GGAAGGCCTACCCGAACTGCTACATCCGCGTTCTCGGCTTCGACAACCAGAGGCAGGTTCAGTGCTCGGCTTTCATCGTCCACAAGCCGGTGAAGA
ATGCGCTCACTGAGAAGCTGGATCGATCTCTTGCGAAGCCCGCTTACAAGCCCCAGCCCCAAGTCTCGTCGGTGGTGACCTCAAGGAGCGGATGGA
GTAGAGATGTGGAGGTGTACCAACCAAAGGATGACAAGACCGTGGTTGACAAGATCAATTCCCTGCAAGCGGGATTGGATGGTCTGCAGCGGGAGC
TCGCACTACTGAGAGCAAGTGCCGCACATGACAAGATCAAGATTCTTCAAGGGGCATTGGATGGTCTGCAGAGGGAGATCTCAGTACTGAGAGCCA
GTGTGGGTCATACTGTAGCTTCGGTGAGCAAGGGCGAGGCAGTGATCAAGGAGTTCATGCGGTTCAAGGTGCACATGGAGGGCTCCATGAACGGCC
ACGAGTTCGAGATCGAGGGCGAGGGCGAGGGCCGCCCCTACGAGGGCACCCAGACCGCCAAGCTGAAGGTGACCAAGGGTGGCCCCCTGCCCTTCT
CCTGGGACATCCTGTCCCCTCAGTTCATGTACGGCTCCAGGGCCTTCATCAAGCACCCCGCCGACATCCCCGACTACTATAAGCAGTCCTTCCCCGAG
GGCTTCAAGTGGGAGCGCGTGATGAACTTCGAGGACGGCGGCGCCGTGACCGTGACCCAGGACACCTCCCTGGAGGACGGCACCCTGATCTACAAG
GTGAAGCTCCGCGGCACCAACTTCCCTCCTGACGGCCCCGTAATGCAGAAGAAGACAATGGGCTGGGAAGCGTCCACCGAGCGGTTGTACCCCGAG
GACGGCGTGCTGAAGGGCGACATTAAGATGGCCCTGCGCCTGAAGGACGGCGGCCGCTACCTGGCGGACTTCAAGACCACCTACAAGGCCAAGAAG
CCCGTGCAGATGCCCGGCGCCTACAACGTCGACCGCAAGTTGGACATCACCTCCCACAACGAGGACTACACCGTGGTGGAACAGTACGAACGCTCCG
AGGGCCGCCACTCCACCGGCGGCATGGACGAGCTGTACAAGTAAGCTTGAGCTCGAATTTCCCCGATCGTTCAAACATTTGGCAATAAAGTTTCTT
AAGATTGAATCCTGTTGCCGGTCTTGCGATGATTATCATATAATTTCTGTTGAATTACGTTAAGCATGTAATAATTAACATGTAATGCATGACGTTA
TTTATGAGATGGGTTTTTATGATTAGAGTCCCGCAATTATACATTTAATACGCGATAGAAAACAAAATATAGCGCGCAAACTAGGATAAATTATCGC
GCGCGGTGTCATCTATGTTACTAGATCGATCCGTATCGATAGCCTCTAGCTAGAGTCGATCGACAAGCTCGAGTTTCTCCATAATAATGTGTGAGTAG
TTCCCAGATAAGGGAATTAGGGTTCCTATAGGGTTTCGCTCATGTGTTGAGCATATAAGAAACCCTTAGTATGTATTTGTATTTGTAAAATACTTCT
ATCAATAAAATTTCTAATTCCTAAAATCAAAATCCAGTACTAAAATCCAGATCGCTACAGAGGAGTCTGAGCAGGACAACTCGCGTAGTGAGAGTT
ACATGTTCGTTGGGTTCTTCCGACACGGACCTGAGTTGGCCAACGTCCCACCTGAGGTCTGTGCCCCGGTGATGAGAAGTGTGCATCTCGTTCTTG
CAGCTCGTCAGTACTTTCAGAATCATGGCGTGCATGGTAGAATGACCCTTATAACGGACTTCGACATGGCAATCGCTAGGT

#### >SR_L2_Tip1_RSSU_eGFP

CTGATCGGGTCAAATGACCTGCCGGAGTACGATTTGAAGGAGGAGGCGGGGCAGGCTGGCCCGATCCTAGTCATGCGCTACCGCAACCTGATCGAG
GGCGAAGCATCCGCCGGTTCCTAATGTACGGAGCAGATGCTAGGGCAAATTGCCCTAGCAGGGGAAAAAGGTCGAAAAGGACTCTTTCCTGTGGAT
AGCACGTACATTGGGAACCCAAAGCCGTACATTGGGAACCGGAACCCGTACATTGGGAACCCAAAGCCGTACATTGGGAACCGGTCACACATGTAA
GTGACTGATATAAAAGAGAAAAAAGGCGATTTTTCCGCCTAAAACTCTTTAAAACTTATTAAAACTCTTAAAACCCGCCTGGCCTGTGCATAACTG
TCTGGCCAGCGCACAGCCGAAGTGCTGCAAAAAGCGCCTACCCTTCGGTCGCTGCGCTCCCTACGCCCCGCCGCTTCGCGTCGGCCTATCGCGGCCG
CTGGCCGCTCAAAAATGGCTGGCCTACGGCCAGGCAATCTACCAGGGCGCGGACAAGCCGCGCCGTCGCCACTCGACCGCCGGCGCCCACATCAAG
GCACCCTGCCTCGCGCGTTTCGGTGATGACGGTGAAAACCTCTGACACATGCAGCTCCCGGTGACGGTCACAGCTTGTCTGTAAGCGGATGCCGGG
AGCAGACAAGCCCGTCAGGGCGCGTCAGCGGGTGTTGGCGGGTGTCGGGGCGCAGCCATGACCCAGTCACGTAGCGATAGCGGAGTGTATACTGGC
TTAACTATGCGGCATCAGAGCAGATTGTACTGAGAGTGCACCATATGCGGTGTGAAATACCGCACAGATGCGTAAGGAGAAAATACCGCATCAGGC
GCTCATCCGCTTCCTCGCTCACTGACTCGCTGCGCTCGGTCGTTCGGCTGCGGCGAGCGGTATCAGCTCACTCAAAGGCGGTAATACGGTTATCCAC
AGAATCAGGGGATAACGCAGGAAAGAACATGTGAGCAAAAGGCCAGCAAAAGGCCAGGAACCGTAAAAAGGCCGCGTTGCTGGCGTTTTTCCATA
GGCTCCGCCCCCCTGACGAGCATCACAAAAATCGACGCTCAAGTCAGAGGTGGCGAAACCCGACAGGACTATAAAGATACCAGGCGTTTCCCCCTG
GAAGCTCCCTCGTGCGCTCTCCTGTTCCGACCCTGCCGCTTACCGGATACCTGTCCGCCTTTCTCCCTTCGGGAAGCGTGGCGCTTTCTCATAGCTC
ACGCTGTAGGTATCTCAGTTCGGTGTAGGTCGTTCGCTCCAAGCTGGGCTGTGTGCACGAACCCCCCGTTCAGCCCGACCGCTGCGCCTTATCCGGT
AACTATCGTCTTGAGTCCAACCCGGTAAGACACGACTTATCGCCACTGGCAGCAGCCACTGGTAACAGGATTAGCAGAGCGAGGTATGTAGGCGGTG
CTACAGAGTTCTTGAAGTGGTGGCCTAACTACGGCTACACTAGAAGGACAGTATTTGGTATCTGCGCTCTGCTGAAGCCAGTTACCTTCGGAAAAA
GAGTTGGTAGCTCTTGATCCGGCAAACAAACCACCGCTGGTAGCGGTGGTTTTTTTGTTTGCAAGCAGCAGATTACGCGCAGAAAAAAAGGATCTC
AAGAAGATCCTTTGATCTTTTCTACGGGGTCTGACGCTCAGTGGAACGAAAACTCACGTTAAGGGATTTTGGTCATGCATTCTAGGTGATTATTTG
CCGACTACCTTGGTGATCTCGCCTTTCACGTAGTGGACAAATTCTTCCAACTGATCTGCGCGCGAGGCCAAGCGATCTTCTTCTTGTCCAAGATAAG
CCTGTCTAGCTTCAAGTATGACGGGCTGATACTGGGCCGGCAGGCGCTCCATTGCCCAGTCGGCAGCGACATCCTTCGGCGCGATTTTGCCGGTTAC
TGCGCTGTACCAAATGCGGGACAACGTAAGCACTACATTTCGCTCATCACCAGCCCAGTCGGGCGGCGAGTTCCATAGCGTTAAGGTTTCATTTAGC
GCCTCAAATAGATCCTGTTCAGGAACCGGATCAAAGAGTTCCTCCGCCGCTGGACCTACCAAGGCAACGCTATGTTCTCTTGCTTTTGTCAGCAAG
ATAGCCAGATCAATGTCGATCGTGGCTGGCTCGAAGATACCTGCAAGAATGTCATTGCGCTGCCATTCTCCAAATTGCAGTTCGCGCTTAGCTGGAT
AACGCCACGGAATGATGTCGTCGTGCACAACAATGGTGACTTCTACAGCGCGGAGAATCTCGCTCTCTCCAGGGGAAGCCGAAGTTTCCAAAAGGT
CGTTGATCAAAGCTCGCCGCGTTGTTTCATCAAGCCTTACGGTCACCGTAACCAGCAAATCAATATCACTGTGTGGCTTCAGGCCGCCATCCACTGC
GGAGCCGTACAAATGTACGGCCAGCAACGTCGGTTCGAGATGGCGCTCGATGACGCCAACTACCTCTGATAGTTGAGTCGATACTTCGGCGATCACC
GCTTCCCTCATAATGTTTAACTTTGTTTTAGGGCGACTGCCCTGCTGCGTAACATCGTTGCTGCTCCATAACATCAAACATCGACCCACGGCGTAAC
GCGCTTGCTGCTTGGATGCCCGAGGCATAGACTGTACCCCAAAAAAACAGTCATAACAAGCCATGAAAACCGCCACTGCGCCGTTACCACCGCTGC
GTTCGGTCAAGGTTCTGGACCAGTTGCGTGAGCGCATACGCTACTTGCATTACAGCTTACGAACCGAACAGGCTTATGTCCACTGGGTTCGTGCCTT
CATCCGTTTCCACGGTGTGCGTCACCCGGCAACCTTGGGTAGCAGCGAAGTCGAGGCATTTCTGTCCTGGCTGGAACAGAACTTATTATTTCCTTCC
TCTTTTCTACAGTATTTAAAGATACCCCAAGAAGCTAATTATAACAAGACGAACTCCAATTCACTGTTCCTTGCATTCTAAAACCTTAAATACCAGA
AAACAGCTTTTTCAAAGTTGTTTTCAAAGTTGGCGTATAACATAGTATCGACGGAGCCGATTTTGAAACCGCGGTGATCACAGGCAGCAACGCTCT
GTCATCGTTACAATCAACATGCTACCCTCCGCGAGATCATCCGTGTTTCAAACCCGGCAGCTTAGTTGCCGTTCTTCCGAATAGCATCGGTAACATG
AGCAAAGTCTGCCGCCTTACAACGGCTCTCCCGCTGACGCCGTCCCGGACTGATGGGCTGCCTGTATCGAGTGGTGATTTTGTGCCGAGCTGCCGGT
CGGGGAGCTGTTGGCTGGCTGGTGGCAGGATATATTGTGGTGTAAACATAACGGATCCGGTCTCAGGAGAATGAGGAGCACCGTCCAAACCTGGAT
AGCCGATATGTGAGTGTTTGGTTGCCTGTCATTTCTAAAAAGCGTATGTCTCGTAGCGTAGATAGCAAGCGCTGTGAGAAAGGTGTGCTCCTATAGC
ATGGTAGCATCTGTCCCATATCTCGTCCGCGAGAAGCCTACATTTCTCGTTGCGTATTAGAGCATGAAAAACCAACGTGAAGTACGTGATTCAGACC

TTGACCAGTTCCAGTTCTCTTCAACAAAAAAACAGTTCTTCAATTCATTTGCTTACATTAAACAATTGTCACACTCTTCTTCCCGGATGACAGCAC
CAAGACACTTCCCCTCCACAAAAATCTCCGAGTCATGTGGACCGGCCTGGATGCTCTTAAAGGGGTTGCCATTGTAGCGGGGTGCCTTAACCACGA
CCTCTTCGATACCTGTCTGAAGCGGCGTGTGAAATCCTCCAGGAGCTTGTTCAAATTTACAAGCTGCAATGGATGTAAACTCATCCAAAGACATGA
CTTGCGAGAAGCCAGAAACGGTCATCAACACTACGAACACCCCTAAACATGGATGGAGAGGCAACATCCTGCATGTGTCTATGTCAGAGGATGCAA
AAGTGAAAGACAAAGGAGAAGGATTGCTCGTGCAATCACTGCAAAGCAAGATGCGAATATCCCTAAAATCCACAGCAGGAAAAGAAAGGAGCCTC
AATAAGTCTCTAAAATGGGAAGAAAAGAAGCACGGGCCAGCGAGACGGAAATGAGGCGATCCGATGCGATGGTTCCGAGAACTGAAGCGGTGCGT
GGGGGGCTAGAAATGAATGGGGAACGTGGAGGGCAAACAAAGGCTGTACTTAGCGGCAAATGCAGATGGCATGCATGCCAAGAACAGCAGCAGTC
GGCACCGCCAGTTTAGCGGAGAGCAACAGGCAGCAAAAGGAAGTGGAACAGGTTTAGGGCCGACGCAGTCGGCGCCTGCCCTAAAACGGCAAGGC
CCACGGCAGCGGTCGCGAGGGCGTTGCCTGCCGGTCGTGGAAAGCCGTGGGGCTGTCAACGGAACCTCACGCTCCCAGCCTGCCCTAAGCGGGGAA
AAAGAAGATCCACATTCTAGGGCTTTCCATCTCGGCTCTCAGCCGTCCGATTAAGAATCGCACGGCCTAGCCCCGCCAGCCTAGCCCTCGGCCGCCC
CCTTGACAGGTTAAGAGCTGGGCAAGCGACTCAGTTGTTCCTTCGTCGCTCGCCCTTCTCACGAGCAGCTCGCCTGCCCCGCCGCCGCCTTGCCTGC
CTCGCGCCTTTTGTCTCCAGGTTCCCTTGCCGCGCCTTCTCCTTGGCTTGTGCTTCTGCTTTGCTTGTGCCTCTGCTGGGCTCTACTTGTGTGGTGT
TGTGCTTTGGCGCCGCCGCCTCGCTGCTGCTGTGGAGCTTGGTGTGTGCTGGGGTTTCTGGGGTTTGCATGTGTCAGGGTTTGTAGATCTGTTCTT
CGGGGCCGTCTTTCTAACGTTTCGTGTGCCGTTGTTTCTGGGGGGTTTGGTGTGGAGCTCGCAGCTTGCAGCTAATGAAAAAGCCTGAACTCACCG
CGACGTCTGTCGAGAAGTTTCTGATCGAAAAGTTCGACAGCGTCTCCGACCTGATGCAGCTCTCGGAGGGCGAAGAATCTCGTGCTTTCAGCTTCG
ATGTAGGAGGGCGTGGATATGTCCTGCGGGTAAATAGCTGCGCCGATGGTTTCTACAAAGATCGTTATGTTTATCGGCACTTTGCATCGGCCGCGCT
CCCGATTCCGGAAGTGCTTGACATTGGGGAGTTTAGCGAGAGCCTGACCTATTGCATCTCCCGCCGTTCACAGGGTGTCACGTTGCAAGACCTGCC
TGAAACCGAACTGCCCGCTGTTCTACAACCGGTCGCGGAGGCTATGGATGCGATCGCTGCGGCCGATCTTAGCCAGACGAGCGGGTTCGGCCCATTC
GGACCGCAAGGAATCGGTCAATACACTACATGGCGTGATTTCATATGCGCGATTGCTGATCCCCATGTGTATCACTGGCAAACTGTGATGGACGACA
CCGTCAGTGCGTCCGTCGCGCAGGCTCTCGATGAGCTGATGCTTTGGGCCGAGGACTGCCCCGAAGTCCGGCACCTCGTGCACGCGGATTTCGGCTC
CAACAATGTCCTGACGGACAATGGCCGCATAACAGCGGTCATTGACTGGAGCGAGGCGATGTTCGGGGATTCCCAATACGAGGTCGCCAACATCTT
CTTCTGGAGGCCGTGGTTGGCTTGTATGGAGCAGCAGACGCGCTACTTCGAGCGGAGGCATCCGGAGCTTGCAGGATCGCCACGACTCCGGGCGTA
TATGCTCCGCATTGGTCTTGACCAACTCTATCAGAGCTTGGTTGACGGCAATTTCGATGATGCAGCTTGGGCGCAGGGTCGATGCGACGCAATCGT
CCGATCCGGAGCCGGGACTGTCGGGCGTACACAAATCGCCCGCAGAAGCGCGGCCGTCTGGACCGATGGCTGTGTAGAAGTACTCGCCGATAGTGG
AAACCGACGCCCCAGCACTCGTCCGAGGGCAAAGAAATAGGCTTGAGCTCGAATTTCCCCGATCGTTCAAACATTTGGCAATAAAGTTTCTTAAGA
TTGAATCCTGTTGCCGGTCTTGCGATGATTATCATATAATTTCTGTTGAATTACGTTAAGCATGTAATAATTAACATGTAATGCATGACGTTATTTAT
GAGATGGGTTTTTATGATTAGAGTCCCGCAATTATACATTTAATACGCGATAGAAAACAAAATATAGCGCGCAAACTAGGATAAATTATCGCGCGCG
GTGTCATCTATGTTACTAGATCGATCCGTATCGATAGCCTCTAGCTAGAGTCGATCGACAAGCTCGAGTTTCTCCATAATAATGTGTGAGTAGTTCCC
AGATAAGGGAATTAGGGTTCCTATAGGGTTTCGCTCATGTGTTGAGCATATAAGAAACCCTTAGTATGTATTTGTATTTGTAAAATACTTCTATCAA
TAAAATTTCTAATTCCTAAAATCAAAATCCAGTACTAAAATCCAGATCGCTAGCAAGGAGTCTGAGCAGGACAACTCGCGTAGTGAGAGTTACATG
TTCGTTGGGTTCTTCCGACACGGACCTGAGTTGGCCAACGTCCCACCTGAGGTCTGTGCCCCGGTGATGAGAAGTGTGCATCTCGTTCTTGCAGCT
CGTCAGTACTTTCAGAATCATGGCGTGCATGGTAGAATGACCCTTATAACGGACTTCGACATGGCAATCGCTATACAGGAGCACCTGGAAGCATTTC
CACACAGCATCTACACCACTACAGGGCTACATGCTCCATAGTGTAGAGCAAACCCTAGTGCAGCTCAGATATTAGATAATGCTGTCCTTCCCAGAAA
GCAAAAGGAATCGACAACTCTGGAAGGCAGACAGCACCAGTTCTGTGCAGCTTCATCCCGAGCTGATGCTGTCCTTCCGAGAAAGCAAAGGAATC
GACACTCCATGGCCGACCGCACAAACACACCATAGATGCTGACCTATCTGGATATGCAACAACCACAACACTTGCGTCTCCACCGCTCCAAGTCCAG
CCTCCAGGAGCCAGGACTCACGGCCGGGTCGAGTGGGCCCAGCCTGGTCCCGTGGCGCCGGGGCCCGGGCCCGGCTCCCCCACTCCCCGGCGCGAT
TCTTACGCGGCGGACGGATAGGCCCACGACCGTCCTACGTGGCGGGCCCCACGTAAGGACAGCGGTGCCTAGGCTGCACCAAATGCCGCGCCACCGT
CCATGTCGTCAGAATCGGTCAGATATTAGCCCCTATCCCTTCACACGCCAGCGCCACGCCACGCCACGCTGCTGACGCCACCCCCCTCTCACCATCAT
CATAGCCTTTCCAGATATCTCCACGCTCCTGCATTACCTGTACATCATTTTAATATGTGCATACATATGTATAATCAATCAGTAAAGGCAAGGCAGGC
TGGATCCCCAGTAGAGCCCATCAATGCCTTGAATCTCGTGTTTCCCGTCAGGTCCTTTCATTCATTCCAAGTCTCCTTCTCATACATACATCCCTCCG
TGATTTAAACTTTATTAAATACCTACATAGCAATGGAATAAATCTATAATGTGGAAGAGACTCCAAGGTGTGACTGAGGTTCCACATGTCTGAAAG
AAGGAGCATCAAAGTATCTACCTTCCCTTCCATTTGGTTGTATTCCCTTCATCCCCTAAACCCAATGCAAGTAGCAAAAGAGGGACACGAGAAGGTA
GTAGTGAAGGTATTGCATTATTGATTTGTGAAAGCGTTTGATGAAACACAAAGGGCACACAAGAAGCGAATGAATGGATCCACACATGGAGGGGC
AGGCAGCTCGAAAAACAGCAGAGGATGCACATGGCCCAGAGCGGATAGGAAAGCAGGCCAGCCTCACTGGACAGTGGACAGCCAGGTGCAGCGCA
CCCTGGCATATTCGAACCGGAAGGGCTGACGTGGGCTCCATTATTTGGGGTGCTGAGTTGGCAAACACGCGCCGGGAAGCTCTCAAAACCGGCATC
TCCATCTCATTCCAACGCCGCAGTCCCATTGCTCACCAACCCCGGCTCCTTTCCCCGCTCTGCTCCTCGGTTTCACGCTCCGCGCGAGCTCTGCTGC
AATGGCTTCCGTGGTCGCTTCCTCCGCCGTCGTGGCCCCAGCTGCATTCTCTGCCGCTTCCTCCAGCGGCTCCGCCTCTGAGGCCTCCAGCATGAGG
GCCTTCGTCGGCCTCAAGTCCGGCGATGGTCTTCTCCAGCAAGACCGCCGACTTGAGCTCCAGGACCGTCAGCAACGGGAGCCGCGTGAATTGCATG
CAGGTGTGGACTCCGGTGGACAACAAGAAGTTCGAAACCCTCTCGTACTTGCCCGCGCTCAGCTCCAATCAAATCGCTAAGCAAATCGACTTCATG
TTGTCGAAGGGCTGGATCCCCTGCCTCGAGTTCGACAAGAGCGCTGAGCTCAAAGTGGTGCGAGGCCAGCCCGGGTTCTACAACGGCCGGTACTGG
ACAATGTGGAAGCTTCCCATGTTCGGCTGCAACGACGCTTCATCCGTGTTGAGGGAAGTTGAGAACTGCAGGAAGGCCTACCCGAACTGCTACATC
CGCGTTCTCGGCTTCGACAACCAGAGGCAGGTTCAGTGCTCGGCTTTCATCGTCCACAAGCCGGTGAAGAATGCGCTCACTGAGAAGCTGGATCGA
TCTCTTGCGAAGCCCGCTTACAAGCCCCAGCCCCAAGTCTCGTCGGTGGTGACCTCAAGGAGCGGATGGAGTAGAGATGTGGAGGTGTACCAACCA
AAGGATGACAAGACCGTGGTTGACAAGATCAATTCCCTGCAAGCGGGATTGGATGGTCTGCAGCGGGAGCTCGCACTACTGAGAGCAAGTGCCGCA
CATGACAAGATCAAGATTCTTCAAGGGGCATTGGATGGTCTGCAGAGGGAGATCTCAGTACTGAGAGCCAGTGTGGGTCATACTGTAGCTTCGGTG
AGCAAGGGCGAGGAGCTGTTCACCGGGGTGGTGCCCATCCTGGTCGAGCTGGACGGCGACGTAAACGGCCACAAGTTCAGCGTGTCCGGCGAGGGC
GAGGGCGATGCCACCTACGGCAAGCTGACCCTGAAGTTCATCTGCACCACCGGCAAGCTGCCCGTGCCCTGGCCCACCCTCGTGACCACCCTGACCT
ACGGCGTGCAGTGCTTCAGCCGCTACCCCGACCACATGAAGCAGCACGACTTCTTCAAGTCCGCCATGCCCGAAGGCTACGTCCAGGAGCGCACCAT
CTTCTTCAAGGACGACGGCAACTACAAGACCCGCGCCGAGGTGAAGTTCGAGGGCGACACCCTGGTGAACCGCATCGAGCTGAAGGGCATCGACTT
CAAGGAGGACGGCAACATCCTGGGGCACAAGCTGGAGTACAACTACAACAGCCACAACGTCTATATCATGGCCGACAAGCAGAAGAACGGCATCAA
GGTGAACTTCAAGATCCGCCACAACATCGAGGACGGCAGCGTGCAGCTCGCCGACCACTACCAGCAGAACACCCCCATCGGCGACGGCCCCGTGCT
GCTGCCCGACAACCACTACCTGAGCACCCAGTCCGCCCTGAGCAAAGACCCCAACGAGAAGCGCGATCACATGGTCCTGCTGGAGTTCGTGACCGCC
GCCGGGATCACTCTCGGCATGGACGAGCTGTACAAGTAAGCTTGAGCTCGAATTTCCCCGATCGTTCAAACATTTGGCAATAAAGTTTCTTAAGAT
TGAATCCTGTTGCCGGTCTTGCGATGATTATCATATAATTTCTGTTGAATTACGTTAAGCATGTAATAATTAACATGTAATGCATGACGTTATTTATG
AGATGGGTTTTTATGATTAGAGTCCCGCAATTATACATTTAATACGCGATAGAAAACAAAATATAGCGCGCAAACTAGGATAAATTATCGCGCGCGG
TGTCATCTATGTTACTAGATCGATCCGTATCGATAGCCTCTAGCTAGAGTCGATCGACAAGCTCGAGTTTCTCCATAATAATGTGTGAGTAGTTCCCA
GATAAGGGAATTAGGGTTCCTATAGGGTTTCGCTCATGTGTTGAGCATATAAGAAACCCTTAGTATGTATTTGTATTTGTAAAATACTTCTATCAAT
AAAATTTCTAATTCCTAAAATCAAAATCCAGTACTAAAATCCAGATCGCTACAGAGGAGTCTGAGCAGGACAACTCGCGTAGTGAGAGTTACATGT
TCGTTGGGTTCTTCCGACACGGACCTGAGTTGGCCAACGTCCCACCTGAGGTCTGTGCCCCGGTGATGAGAAGTGTGCATCTCGTTCTTGCAGCTC
GTCAGTACTTTCAGAATCATGGCGTGCATGGTAGAATGACCCTTATAACGGACTTCGACATGGCAATCGCTAGGTATACTTGAGACCGGATCCTGAC
AGGATATATTGGCGGGTAAACCTAAGAGAAAAGAGCGTTTATTAGAATAATCGGATATTTAAAAGGGCGTGAAAAGGTTTATCCGTTCGTCCATTT
GTATGTGCATGCCAACCACAGGGTTCCCCTCGGGATCAAAGTACTTTGATCCAACCCCTCCGCTGCTATAGTGCAGTCGGCTTCTGACGTTCAGTGC
AGCCGTCATCTGAAAACGACATGTCGCACAAGTCCTAAGTTACGCGACAGGCTGCCGCCCTGCCCTTTTCCTGGCGTTTTCTTGTCGCGTGTTTTAG
TCGCATAAAGTAGAATACTTGCGACTAGAACCGGAGACATTACGCCATGAACAAGAGCGCCGCCGCTGGCCTGCTGGGCTATGCCCGCGTCAGCACC

GACGACCAGGACTTGACCAACCAACGGGCCGAACTGCACGCGGCCGGCTGCACCAAGCTGTTTTCCGAGAAGATCACCGGCACCAGGCGCGACCGC
CCGGAGCTGGCCAGGATGCTTGACCACCTACGCCCTGGCGACGTTGTGACAGTGACCAGGCTAGACCGCCTGGCCCGCAGCACCCGCGACCTACTGG
ACATTGCCGAGCGCATCCAGGAGGCCGGCGCGGGCCTGCGTAGCCTGGCAGAGCCGTGGGCCGACACCACCACGCCGGCCGGCCGCATGGTGTTGA
CCGTGTTCGCCGGCATTGCCGAGTTCGAGCGTTCCCTAATCATCGACCGCACCCGGAGCGGGCGCGAGGCCGCCAAGGCCCGAGGCGTGAAGTTTG
GCCCCCGCCCTACCCTCACCCCGGCACAGATCGCGCACGCCCGCGAGCTGATCGACCAGGAAGGCCGCACCGTGAAAGAGGCGGCTGCACTGCTTGG
CGTGCATCGCTCGACCCTGTACCGCGCACTTGAGCGCAGCGAGGAAGTGACGCCCACCGAGGCCAGGCGGCGCGGTGCCTTCCGTGAGGACGCATT
GACCGAGGCCGACGCCCTGGCGGCCGCCGAGAATGAACGCCAAGAGGAACAAGCATGAAACCGCACCAGGACGGCCAGGACGAACCGTTTTTCATT
ACCGAAGAGATCGAGGCGGAGATGATCGCGGCCGGGTACGTGTTCGAGCCGCCCGCGCACCTCTCAACCGTGCGGCTGCATGAAATCCTGGCCGGT
TTGTCTGATGCCAAGCTGGCGGCCTGGCCGGCCAGCTTGGCCGCTGAAGAAACCGAGCGCCGCCGTCTAAAAAGGTGATGTGTATTTGAGTAAAAC
AGCTTGCGTCATGCGGTCGCTGCGTATATGATGCGATGAGTAAATAAACAAATACGCAAGGGGAACGCATGAAGGTTATCGCTGTACTTAACCAGA
AAGGCGGGTCAGGCAAGACGACCATCGCAACCCATCTAGCCCGCGCCCTGCAACTCGCCGGGGCCGATGTTCTGTTAGTCGATTCCGATCCCCAGGG
CAGTGCCCGCGATTGGGCGGCCGTGCGGGAAGATCAACCGCTAACCGTTGTCGGCATCGACCGCCCGACGATTGACCGCGACGTGAAGGCCATCGG
CCGGCGCGACTTCGTAGTGATCGACGGAGCGCCCCAGGCGGCGGACTTGGCTGTGTCCGCGATCAAGGCAGCCGACTTCGTGCTGATTCCGGTGCA
GCCAAGCCCTTACGACATATGGGCCACCGCCGACCTGGTGGAGCTGGTTAAGCAGCGCATTGAGGTCACGGATGGAAGGCTACAAGCGGCCTTTGT
CGTGTCGCGGGCGATCAAAGGCACGCGCATCGGCGGTGAGGTTGCCGAGGCGCTGGCCGGGTACGAGCTGCCCATTCTTGAGTCCCGTATCACGCA
GCGCGTGAGCTACCCAGGCACTGCCGCCGCCGGCACAACCGTTCTTGAATCAGAACCCGAGGGCGACGCTGCCCGCGAGGTCCAGGCGCTGGCCGC
TGAAATTAAATCAAAACTCATTTGAGTTAATGAGGTAAAGAGAAAATGAGCAAAAGCACAAACACGCTAAGTGCCGGCCGTCCGAGCGCACGCAG
CAGCAAGGCTGCAACGTTGGCCAGCCTGGCAGACACGCCAGCCATGAAGCGGGTCAACTTTCAGTTGCCGGCGGAGGATCACACCAAGCTGAAGAT
GTACGCGGTACGCCAAGGCAAGACCATTACCGAGCTGCTATCTGAATACATCGCGCAGCTACCAGAGTAAATGAGCAAATGAATAAATGAGTAGATG
AATTTTAGCGGCTAAAGGAGGCGGCATGGAAAATCAAGAACAACCAGGCACCGACGCCGTGGAATGCCCCATGTGTGGAGGAACGGGCGGTTGGC
CAGGCGTAAGCGGCTGGGTTGCCTGCCGGCCCTGCAATGGCACTGGAACCCCCAAGCCCGAGGAATCGGCGTGAGCGGTCGCAAACCATCCGGCCC
GGTACAAATCGGCGCGGCGCTGGGTGATGACCTGGTGGAGAAGTTGAAGGCCGCGCAGGCCGCCCAGCGGCAACGCATCGAGGCAGAAGCACGCCC
CGGTGAATCGTGGCAAGCGGCCGCTGATCGAATCCGCAAAGAATCCCGGCAACCGCCGGCAGCCGGTGCGCCGTCGATTAGGAAGCCGCCCAAGGG
CGACGAGCAACCAGATTTTTTCGTTCCGATGCTCTATGACGTGGGCACCCGCGATAGTCGCAGCATCATGGACGTGGCCGTTTTCCGTCTGTCGAAG
CGTGACCGACGAGCTGGCGAGGTGATCCGCTACGAGCTTCCAGACGGGCACGTAGAGGTTTCCGCAGGGCCGGCCGGCATGGCCAGTGTGTGGGAT
TACGACCTGGTACTGATGGCGGTTTCCCATCTAACCGAATCCATGAACCGATACCGGGAAGGGAAGGGAGACAAGCCCGGCCGCGTGTTCCGTCCAC
ACGTTGCGGACGTACTCAAGTTCTGCCGGCGAGCCGATGGCGGAAAGCAGAAAGACGACCTGGTAGAAACCTGCATTCGGTTAAACACCACGCACG
TTGCCATGCAGCGTACGAAGAAGGCCAAGAACGGCCGCCTGGTGACGGTATCCGAGGGTGAAGCCTTGATTAGCCGCTACAAGATCGTAAAGAGCG
AAACCGGGCGGCCGGAGTACATCGAGATCGAGCTAGCTGATTGGATGTACCGCGAGATCACAGAAGGCAAGAACCCGGACGTGCTGACGGTTCACC
CCGATTACTTTTTGATCGATCCCGGCATCGGCCGTTTTCTCTACCGCCTGGCACGCCGCGCCGCAGGCAAGGCAGAAGCCAGATGGTTGTTCAAGAC
GATCTACGAACGCAGTGGCAGCGCCGGAGAGTTCAAGAAGTTCTGTTTCACCGTGCGCAAG

#### >SR_L2_Tip1_LCIB_mVenus

ATACTTGAGACCGGATCCTGACAGGATATATTGGCGGGTAAACCTAAGAGAAAAGAGCGTTTATTAGAATAATCGGATATTTAAAAGGGCGTGAAA
AGGTTTATCCGTTCGTCCATTTGTATGTGCATGCCAACCACAGGGTTCCCCTCGGGATCAAAGTACTTTGATCCAACCCCTCCGCTGCTATAGTGCA
GTCGGCTTCTGACGTTCAGTGCAGCCGTCATCTGAAAACGACATGTCGCACAAGTCCTAAGTTACGCGACAGGCTGCCGCCCTGCCCTTTTCCTGGC
GTTTTCTTGTCGCGTGTTTTAGTCGCATAAAGTAGAATACTTGCGACTAGAACCGGAGACATTACGCCATGAACAAGAGCGCCGCCGCTGGCCTGC
TGGGCTATGCCCGCGTCAGCACCGACGACCAGGACTTGACCAACCAACGGGCCGAACTGCACGCGGCCGGCTGCACCAAGCTGTTTTCCGAGAAGA
TCACCGGCACCAGGCGCGACCGCCCGGAGCTGGCCAGGATGCTTGACCACCTACGCCCTGGCGACGTTGTGACAGTGACCAGGCTAGACCGCCTGGC
CCGCAGCACCCGCGACCTACTGGACATTGCCGAGCGCATCCAGGAGGCCGGCGCGGGCCTGCGTAGCCTGGCAGAGCCGTGGGCCGACACCACCACG
CCGGCCGGCCGCATGGTGTTGACCGTGTTCGCCGGCATTGCCGAGTTCGAGCGTTCCCTAATCATCGACCGCACCCGGAGCGGGCGCGAGGCCGCCA
AGGCCCGAGGCGTGAAGTTTGGCCCCCGCCCTACCCTCACCCCGGCACAGATCGCGCACGCCCGCGAGCTGATCGACCAGGAAGGCCGCACCGTGAA
AGAGGCGGCTGCACTGCTTGGCGTGCATCGCTCGACCCTGTACCGCGCACTTGAGCGCAGCGAGGAAGTGACGCCCACCGAGGCCAGGCGGCGCGG
TGCCTTCCGTGAGGACGCATTGACCGAGGCCGACGCCCTGGCGGCCGCCGAGAATGAACGCCAAGAGGAACAAGCATGAAACCGCACCAGGACGGC
CAGGACGAACCGTTTTTCATTACCGAAGAGATCGAGGCGGAGATGATCGCGGCCGGGTACGTGTTCGAGCCGCCCGCGCACCTCTCAACCGTGCGG
CTGCATGAAATCCTGGCCGGTTTGTCTGATGCCAAGCTGGCGGCCTGGCCGGCCAGCTTGGCCGCTGAAGAAACCGAGCGCCGCCGTCTAAAAAGG
TGATGTGTATTTGAGTAAAACAGCTTGCGTCATGCGGTCGCTGCGTATATGATGCGATGAGTAAATAAACAAATACGCAAGGGGAACGCATGAAGG
TTATCGCTGTACTTAACCAGAAAGGCGGGTCAGGCAAGACGACCATCGCAACCCATCTAGCCCGCGCCCTGCAACTCGCCGGGGCCGATGTTCTGTT
AGTCGATTCCGATCCCCAGGGCAGTGCCCGCGATTGGGCGGCCGTGCGGGAAGATCAACCGCTAACCGTTGTCGGCATCGACCGCCCGACGATTGAC
CGCGACGTGAAGGCCATCGGCCGGCGCGACTTCGTAGTGATCGACGGAGCGCCCCAGGCGGCGGACTTGGCTGTGTCCGCGATCAAGGCAGCCGAC
TTCGTGCTGATTCCGGTGCAGCCAAGCCCTTACGACATATGGGCCACCGCCGACCTGGTGGAGCTGGTTAAGCAGCGCATTGAGGTCACGGATGGA
AGGCTACAAGCGGCCTTTGTCGTGTCGCGGGCGATCAAAGGCACGCGCATCGGCGGTGAGGTTGCCGAGGCGCTGGCCGGGTACGAGCTGCCCATT
CTTGAGTCCCGTATCACGCAGCGCGTGAGCTACCCAGGCACTGCCGCCGCCGGCACAACCGTTCTTGAATCAGAACCCGAGGGCGACGCTGCCCGCG
AGGTCCAGGCGCTGGCCGCTGAAATTAAATCAAAACTCATTTGAGTTAATGAGGTAAAGAGAAAATGAGCAAAAGCACAAACACGCTAAGTGCCG
GCCGTCCGAGCGCACGCAGCAGCAAGGCTGCAACGTTGGCCAGCCTGGCAGACACGCCAGCCATGAAGCGGGTCAACTTTCAGTTGCCGGCGGAGG
ATCACACCAAGCTGAAGATGTACGCGGTACGCCAAGGCAAGACCATTACCGAGCTGCTATCTGAATACATCGCGCAGCTACCAGAGTAAATGAGCAA
ATGAATAAATGAGTAGATGAATTTTAGCGGCTAAAGGAGGCGGCATGGAAAATCAAGAACAACCAGGCACCGACGCCGTGGAATGCCCCATGTGTG
GAGGAACGGGCGGTTGGCCAGGCGTAAGCGGCTGGGTTGCCTGCCGGCCCTGCAATGGCACTGGAACCCCCAAGCCCGAGGAATCGGCGTGAGCGG
TCGCAAACCATCCGGCCCGGTACAAATCGGCGCGGCGCTGGGTGATGACCTGGTGGAGAAGTTGAAGGCCGCGCAGGCCGCCCAGCGGCAACGCAT
CGAGGCAGAAGCACGCCCCGGTGAATCGTGGCAAGCGGCCGCTGATCGAATCCGCAAAGAATCCCGGCAACCGCCGGCAGCCGGTGCGCCGTCGAT
TAGGAAGCCGCCCAAGGGCGACGAGCAACCAGATTTTTTCGTTCCGATGCTCTATGACGTGGGCACCCGCGATAGTCGCAGCATCATGGACGTGGC
CGTTTTCCGTCTGTCGAAGCGTGACCGACGAGCTGGCGAGGTGATCCGCTACGAGCTTCCAGACGGGCACGTAGAGGTTTCCGCAGGGCCGGCCGG
CATGGCCAGTGTGTGGGATTACGACCTGGTACTGATGGCGGTTTCCCATCTAACCGAATCCATGAACCGATACCGGGAAGGGAAGGGAGACAAGCC
CGGCCGCGTGTTCCGTCCACACGTTGCGGACGTACTCAAGTTCTGCCGGCGAGCCGATGGCGGAAAGCAGAAAGACGACCTGGTAGAAACCTGCAT
TCGGTTAAACACCACGCACGTTGCCATGCAGCGTACGAAGAAGGCCAAGAACGGCCGCCTGGTGACGGTATCCGAGGGTGAAGCCTTGATTAGCCG
CTACAAGATCGTAAAGAGCGAAACCGGGCGGCCGGAGTACATCGAGATCGAGCTAGCTGATTGGATGTACCGCGAGATCACAGAAGGCAAGAACCC
GGACGTGCTGACGGTTCACCCCGATTACTTTTTGATCGATCCCGGCATCGGCCGTTTTCTCTACCGCCTGGCACGCCGCGCCGCAGGCAAGGCAGAA
GCCAGATGGTTGTTCAAGACGATCTACGAACGCAGTGGCAGCGCCGGAGAGTTCAAGAAGTTCTGTTTCACCGTGCGCAAGCTGATCGGGTCAAAT
GACCTGCCGGAGTACGATTTGAAGGAGGAGGCGGGGCAGGCTGGCCCGATCCTAGTCATGCGCTACCGCAACCTGATCGAGGGCGAAGCATCCGCC
GGTTCCTAATGTACGGAGCAGATGCTAGGGCAAATTGCCCTAGCAGGGGAAAAAGGTCGAAAAGGACTCTTTCCTGTGGATAGCACGTACATTGGG
AACCCAAAGCCGTACATTGGGAACCGGAACCCGTACATTGGGAACCCAAAGCCGTACATTGGGAACCGGTCACACATGTAAGTGACTGATATAAAA
GAGAAAAAAGGCGATTTTTCCGCCTAAAACTCTTTAAAACTTATTAAAACTCTTAAAACCCGCCTGGCCTGTGCATAACTGTCTGGCCAGCGCACA
GCCGAAGTGCTGCAAAAAGCGCCTACCCTTCGGTCGCTGCGCTCCCTACGCCCCGCCGCTTCGCGTCGGCCTATCGCGGCCGCTGGCCGCTCAAAAA
TGGCTGGCCTACGGCCAGGCAATCTACCAGGGCGCGGACAAGCCGCGCCGTCGCCACTCGACCGCCGGCGCCCACATCAAGGCACCCTGCCTCGCGC

GTTTCGGTGATGACGGTGAAAACCTCTGACACATGCAGCTCCCGGTGACGGTCACAGCTTGTCTGTAAGCGGATGCCGGGAGCAGACAAGCCCGTC
AGGGCGCGTCAGCGGGTGTTGGCGGGTGTCGGGGCGCAGCCATGACCCAGTCACGTAGCGATAGCGGAGTGTATACTGGCTTAACTATGCGGCATC
AGAGCAGATTGTACTGAGAGTGCACCATATGCGGTGTGAAATACCGCACAGATGCGTAAGGAGAAAATACCGCATCAGGCGCTCATCCGCTTCCTCG
CTCACTGACTCGCTGCGCTCGGTCGTTCGGCTGCGGCGAGCGGTATCAGCTCACTCAAAGGCGGTAATACGGTTATCCACAGAATCAGGGGATAACG
CAGGAAAGAACATGTGAGCAAAAGGCCAGCAAAAGGCCAGGAACCGTAAAAAGGCCGCGTTGCTGGCGTTTTTCCATAGGCTCCGCCCCCCTGAC
GAGCATCACAAAAATCGACGCTCAAGTCAGAGGTGGCGAAACCCGACAGGACTATAAAGATACCAGGCGTTTCCCCCTGGAAGCTCCCTCGTGCGC
TCTCCTGTTCCGACCCTGCCGCTTACCGGATACCTGTCCGCCTTTCTCCCTTCGGGAAGCGTGGCGCTTTCTCATAGCTCACGCTGTAGGTATCTCAG
TTCGGTGTAGGTCGTTCGCTCCAAGCTGGGCTGTGTGCACGAACCCCCCGTTCAGCCCGACCGCTGCGCCTTATCCGGTAACTATCGTCTTGAGTCC
AACCCGGTAAGACACGACTTATCGCCACTGGCAGCAGCCACTGGTAACAGGATTAGCAGAGCGAGGTATGTAGGCGGTGCTACAGAGTTCTTGAAG
TGGTGGCCTAACTACGGCTACACTAGAAGGACAGTATTTGGTATCTGCGCTCTGCTGAAGCCAGTTACCTTCGGAAAAAGAGTTGGTAGCTCTTGA
TCCGGCAAACAAACCACCGCTGGTAGCGGTGGTTTTTTTGTTTGCAAGCAGCAGATTACGCGCAGAAAAAAAGGATCTCAAGAAGATCCTTTGAT
CTTTTCTACGGGGTCTGACGCTCAGTGGAACGAAAACTCACGTTAAGGGATTTTGGTCATGCATTCTAGGTGATTATTTGCCGACTACCTTGGTGA
TCTCGCCTTTCACGTAGTGGACAAATTCTTCCAACTGATCTGCGCGCGAGGCCAAGCGATCTTCTTCTTGTCCAAGATAAGCCTGTCTAGCTTCAAG
TATGACGGGCTGATACTGGGCCGGCAGGCGCTCCATTGCCCAGTCGGCAGCGACATCCTTCGGCGCGATTTTGCCGGTTACTGCGCTGTACCAAATG
CGGGACAACGTAAGCACTACATTTCGCTCATCACCAGCCCAGTCGGGCGGCGAGTTCCATAGCGTTAAGGTTTCATTTAGCGCCTCAAATAGATCCT
GTTCAGGAACCGGATCAAAGAGTTCCTCCGCCGCTGGACCTACCAAGGCAACGCTATGTTCTCTTGCTTTTGTCAGCAAGATAGCCAGATCAATGT
CGATCGTGGCTGGCTCGAAGATACCTGCAAGAATGTCATTGCGCTGCCATTCTCCAAATTGCAGTTCGCGCTTAGCTGGATAACGCCACGGAATGAT
GTCGTCGTGCACAACAATGGTGACTTCTACAGCGCGGAGAATCTCGCTCTCTCCAGGGGAAGCCGAAGTTTCCAAAAGGTCGTTGATCAAAGCTCG
CCGCGTTGTTTCATCAAGCCTTACGGTCACCGTAACCAGCAAATCAATATCACTGTGTGGCTTCAGGCCGCCATCCACTGCGGAGCCGTACAAATGT
ACGGCCAGCAACGTCGGTTCGAGATGGCGCTCGATGACGCCAACTACCTCTGATAGTTGAGTCGATACTTCGGCGATCACCGCTTCCCTCATAATGT
TTAACTTTGTTTTAGGGCGACTGCCCTGCTGCGTAACATCGTTGCTGCTCCATAACATCAAACATCGACCCACGGCGTAACGCGCTTGCTGCTTGGA
TGCCCGAGGCATAGACTGTACCCCAAAAAAACAGTCATAACAAGCCATGAAAACCGCCACTGCGCCGTTACCACCGCTGCGTTCGGTCAAGGTTCT
GGACCAGTTGCGTGAGCGCATACGCTACTTGCATTACAGCTTACGAACCGAACAGGCTTATGTCCACTGGGTTCGTGCCTTCATCCGTTTCCACGGT
GTGCGTCACCCGGCAACCTTGGGTAGCAGCGAAGTCGAGGCATTTCTGTCCTGGCTGGAACAGAACTTATTATTTCCTTCCTCTTTTCTACAGTATT
TAAAGATACCCCAAGAAGCTAATTATAACAAGACGAACTCCAATTCACTGTTCCTTGCATTCTAAAACCTTAAATACCAGAAAACAGCTTTTTCAA
AGTTGTTTTCAAAGTTGGCGTATAACATAGTATCGACGGAGCCGATTTTGAAACCGCGGTGATCACAGGCAGCAACGCTCTGTCATCGTTACAATC
AACATGCTACCCTCCGCGAGATCATCCGTGTTTCAAACCCGGCAGCTTAGTTGCCGTTCTTCCGAATAGCATCGGTAACATGAGCAAAGTCTGCCGC
CTTACAACGGCTCTCCCGCTGACGCCGTCCCGGACTGATGGGCTGCCTGTATCGAGTGGTGATTTTGTGCCGAGCTGCCGGTCGGGGAGCTGTTGG
CTGGCTGGTGGCAGGATATATTGTGGTGTAAACATAACGGATCCGGTCTCAGGAGAATGAGGAGCACCGTCCAAACCTGGATAGCCGATATGTGAG
TGTTTGGTTGCCTGTCATTTCTAAAAAGCGTATGTCTCGTAGCGTAGATAGCAAGCGCTGTGAGAAAGGTGTGCTCCTATAGCATGGTAGCATCTGT
CCCATATCTCGTCCGCGAGAAGCCTACATTTCTCGTTGCGTATTAGAGCATGAAAAACCAACGTGAAGTACGTGATTCAGACCTTGACCAGTTCCAG
TTCTCTTCAACAAAAAAACAGTTCTTCAATTCATTTGCTTACATTAAACAATTGTCACACTCTTCTTCCCGGATGACAGCACCAAGACACTTCCCC
TCCACAAAAATCTCCGAGTCATGTGGACCGGCCTGGATGCTCTTAAAGGGGTTGCCATTGTAGCGGGGTGCCTTAACCACGACCTCTTCGATACCTG
TCTGAAGCGGCGTGTGAAATCCTCCAGGAGCTTGTTCAAATTTACAAGCTGCAATGGATGTAAACTCATCCAAAGACATGACTTGCGAGAAGCCAG
AAACGGTCATCAACACTACGAACACCCCTAAACATGGATGGAGAGGCAACATCCTGCATGTGTCTATGTCAGAGGATGCAAAAGTGAAAGACAAAG
GAGAAGGATTGCTCGTGCAATCACTGCAAAGCAAGATGCGAATATCCCTAAAATCCACAGCAGGAAAAGAAAGGAGCCTCAATAAGTCTCTAAAAT
GGGAAGAAAAGAAGCACGGGCCAGCGAGACGGAAATGAGGCGATCCGATGCGATGGTTCCGAGAACTGAAGCGGTGCGTGGGGGGCTAGAAATGA
ATGGGGAACGTGGAGGGCAAACAAAGGCTGTACTTAGCGGCAAATGCAGATGGCATGCATGCCAAGAACAGCAGCAGTCGGCACCGCCAGTTTAG
CGGAGAGCAACAGGCAGCAAAAGGAAGTGGAACAGGTTTAGGGCCGACGCAGTCGGCGCCTGCCCTAAAACGGCAAGGCCCACGGCAGCGGTCGC
GAGGGCGTTGCCTGCCGGTCGTGGAAAGCCGTGGGGCTGTCAACGGAACCTCACGCTCCCAGCCTGCCCTAAGCGGGGAAAAAGAAGATCCACATT
CTAGGGCTTTCCATCTCGGCTCTCAGCCGTCCGATTAAGAATCGCACGGCCTAGCCCCGCCAGCCTAGCCCTCGGCCGCCCCCTTGACAGGTTAAGA
GCTGGGCAAGCGACTCAGTTGTTCCTTCGTCGCTCGCCCTTCTCACGAGCAGCTCGCCTGCCCCGCCGCCGCCTTGCCTGCCTCGCGCCTTTTGTCT
CCAGGTTCCCTTGCCGCGCCTTCTCCTTGGCTTGTGCTTCTGCTTTGCTTGTGCCTCTGCTGGGCTCTACTTGTGTGGTGTTGTGCTTTGGCGCCGC
CGCCTCGCTGCTGCTGTGGAGCTTGGTGTGTGCTGGGGTTTCTGGGGTTTGCATGTGTCAGGGTTTGTAGATCTGTTCTTCGGGGCCGTCTTTCTA
ACGTTTCGTGTGCCGTTGTTTCTGGGGGGTTTGGTGTGGAGCTCGCAGCTTGCAGCTAATGAAAAAGCCTGAACTCACCGCGACGTCTGTCGAGAA
GTTTCTGATCGAAAAGTTCGACAGCGTCTCCGACCTGATGCAGCTCTCGGAGGGCGAAGAATCTCGTGCTTTCAGCTTCGATGTAGGAGGGCGTGG
ATATGTCCTGCGGGTAAATAGCTGCGCCGATGGTTTCTACAAAGATCGTTATGTTTATCGGCACTTTGCATCGGCCGCGCTCCCGATTCCGGAAGTG
CTTGACATTGGGGAGTTTAGCGAGAGCCTGACCTATTGCATCTCCCGCCGTTCACAGGGTGTCACGTTGCAAGACCTGCCTGAAACCGAACTGCCC
GCTGTTCTACAACCGGTCGCGGAGGCTATGGATGCGATCGCTGCGGCCGATCTTAGCCAGACGAGCGGGTTCGGCCCATTCGGACCGCAAGGAATC
GGTCAATACACTACATGGCGTGATTTCATATGCGCGATTGCTGATCCCCATGTGTATCACTGGCAAACTGTGATGGACGACACCGTCAGTGCGTCCG
TCGCGCAGGCTCTCGATGAGCTGATGCTTTGGGCCGAGGACTGCCCCGAAGTCCGGCACCTCGTGCACGCGGATTTCGGCTCCAACAATGTCCTGA
CGGACAATGGCCGCATAACAGCGGTCATTGACTGGAGCGAGGCGATGTTCGGGGATTCCCAATACGAGGTCGCCAACATCTTCTTCTGGAGGCCGT
GGTTGGCTTGTATGGAGCAGCAGACGCGCTACTTCGAGCGGAGGCATCCGGAGCTTGCAGGATCGCCACGACTCCGGGCGTATATGCTCCGCATTG
GTCTTGACCAACTCTATCAGAGCTTGGTTGACGGCAATTTCGATGATGCAGCTTGGGCGCAGGGTCGATGCGACGCAATCGTCCGATCCGGAGCCG
GGACTGTCGGGCGTACACAAATCGCCCGCAGAAGCGCGGCCGTCTGGACCGATGGCTGTGTAGAAGTACTCGCCGATAGTGGAAACCGACGCCCCA
GCACTCGTCCGAGGGCAAAGAAATAGGCTTGAGCTCGAATTTCCCCGATCGTTCAAACATTTGGCAATAAAGTTTCTTAAGATTGAATCCTGTTGC
CGGTCTTGCGATGATTATCATATAATTTCTGTTGAATTACGTTAAGCATGTAATAATTAACATGTAATGCATGACGTTATTTATGAGATGGGTTTTT
ATGATTAGAGTCCCGCAATTATACATTTAATACGCGATAGAAAACAAAATATAGCGCGCAAACTAGGATAAATTATCGCGCGCGGTGTCATCTATGT
TACTAGATCGATCCGTATCGATAGCCTCTAGCTAGAGTCGATCGACAAGCTCGAGTTTCTCCATAATAATGTGTGAGTAGTTCCCAGATAAGGGAAT
TAGGGTTCCTATAGGGTTTCGCTCATGTGTTGAGCATATAAGAAACCCTTAGTATGTATTTGTATTTGTAAAATACTTCTATCAATAAAATTTCTAAT
TCCTAAAATCAAAATCCAGTACTAAAATCCAGATCGCTAGCAAGGAGTCTGAGCAGGACAACTCGCGTAGTGAGAGTTACATGTTCGTTGGGTTCT
TCCGACACGGACCTGAGTTGGCCAACGTCCCACCTGAGGTCTGTGCCCCGGTGATGAGAAGTGTGCATCTCGTTCTTGCAGCTCGTCAGTACTTTC
AGAATCATGGCGTGCATGGTAGAATGACCCTTATAACGGACTTCGACATGGCAATCGCTATACAGGAGTCTGAGCAGGACAACTCGCGTAGTGAGA
GTTACATGTTCGTTGGGTTCTTCCGACACGGACCTGAGTTGGCCAACGTCCCACCTGAGGTCTGTGCCCCGGTGATGAGAAGTGTGCATCTCGTTC
TTGCAGCTCGTCAGTACTTTCAGAATCATGGCGTGCATGGTAGAATGACCCTTATAACGGACTTCGACATGGCAATCGCTACAGAGGAGCACCTGG
AAGCATTTCCACACAGCATCTACACCACTACAGGGCTACATGCTCCATAGTGTAGAGCAAACCCTAGTGCAGCTCAGATATTAGATAATGCTGTCCT
TCCCAGAAAGCAAAAGGAATCGACAACTCTGGAAGGCAGACAGCACCAGTTCTGTGCAGCTTCATCCCGAGCTGATGCTGTCCTTCCGAGAAAGCA
AAGGAATCGACACTCCATGGCCGACCGCACAAACACACCATAGATGCTGACCTATCTGGATATGCAACAACCACAACACTTGCGTCTCCACCGCTCC
AAGTCCAGCCTCCAGGAGCCAGGACTCACGGCCGGGTCGAGTGGGCCCAGCCTGGTCCCGTGGCGCCGGGGCCCGGGCCCGGCTCCCCCACTCCCC
GGCGCGATTCTTACGCGGCGGACGGATAGGCCCACGACCGTCCTACGTGGCGGGCCCCACGTAAGGACAGCGGTGCCTAGGCTGCACCAAATGCCGC
GCCACCGTCCATGTCGTCAGAATCGGTCAGATATTAGCCCCTATCCCTTCACACGCCAGCGCCACGCCACGCCACGCTGCTGACGCCACCCCCCTCTC
ACCATCATCATAGCCTTTCCAGATATCTCCACGCTCCTGCATTACCTGTACATCATTTTAATATGTGCATACATATGTATAATCAATCAGTAAAGGCA
AGGCAGGCTGGATCCCCAGTAGAGCCCATCAATGCCTTGAATCTCGTGTTTCCCGTCAGGTCCTTTCATTCATTCCAAGTCTCCTTCTCATACATAC
ATCCCTCCGTGATTTAAACTTTATTAAATACCTACATAGCAATGGAATAAATCTATAATGTGGAAGAGACTCCAAGGTGTGACTGAGGTTCCACATG

TCTGAAAGAAGGAGCATCAAAGTATCTACCTTCCCTTCCATTTGGTTGTATTCCCTTCATCCCCTAAACCCAATGCAAGTAGCAAAAGAGGGACACG
AGAAGGTAGTAGTGAAGGTATTGCATTATTGATTTGTGAAAGCGTTTGATGAAACACAAAGGGCACACAAGAAGCGAATGAATGGATCCACACAT
GGAGGGGCAGGCAGCTCGAAAAACAGCAGAGGATGCACATGGCCCAGAGCGGATAGGAAAGCAGGCCAGCCTCACTGGACAGTGGACAGCCAGGT
GCAGCGCACCCTGGCATATTCGAACCGGAAGGGCTGACGTGGGCTCCATTATTTGGGGTGCTGAGTTGGCAAACACGCGCCGGGAAGCTCTCAAAA
CCGGCATCTCCATCTCATTCCAACGCCGCAGTCCCATTGCTCACCAACCCCGGCTCCTTTCCCCGCTCTGCTCCTCGGTTTCACGCTCCGCGCGAGCT
CTGCTGCAATGGCAGCCGGATTGTGCAGCATGTCGGGCCTTGCTCTCTCCGCCGACGTTCGCAGGCTCTCCCCTGCCCTCCCTGCCCGCGCGGTTGA
TCCCCTGCGCCTCTCAGGCCCCCGCAGGCTCCATGTAGTGTCAGCCTCCGCGTCTGCGAACTCGCAGCCCGCTGGAGAGGAGCTGAAGCTGCAGGCG
CAGCCGCAGCAGAGCTATGCTGCCTCTGTCCGGGAAACCATGCCCTTCGCTTCCGCTTCCACTTCCGTGGAAGCACCTCCTCCCTCTGCACCCGTAG
AGGGCTCTGGGAAGTCGCCTGTCTTCCTGTACGTGGCAGCGGCCCTGGCTGCTGTGGGAATCTTCGCTGCTCTGAAGACCCGGGGCGGCAGCGCAG
CCCCAGCTCCTGCTACTCCGGCTGCCCCGCCGGCTCCCCTCACCAAGCTGCAGAAGCCCCGCCGGGAAATCGCTCCTGCTGCGTTGAAGAAGATCTC
CACCGACTTCTCTGTTGCTGTTGTGGACAACAAGTTCGTGAAAAGCGTGGGCTCAGCTCTTGCTGGCCTTGGCTTCACCAAGGAGAACACCATTGC
GTTGGTTAGTACTTGCAGAGATGAGATCACAAGGCCACTGATCGGCCTTGTGGAAGAGGTCTACGGGGACTCCTTCAACATCGCTGGTCTTGGCGG
TGCTATCGTTTGTGGGAAAGTTGGGTTCAAGGCTGGCATTGGTCACTCGCCAGATGCCAAGCAGGACAGAGAGAAGTATGTGTTCTTCGCCTTCCC
TCACGTTGGTGTTGATGAGCAAGGAAACGAAGGTCTTGTGAAGAGAGAGGGCCAAGCCAAGCCCTCAACTGCCTGTGCAGCTCTCCTCGCAGTCA
AGGGCGAGATTGAACGGGGTGCCACTCCCCATTCAGGACCTCTTCCCCTTGATGACATGGAGTTCGGCCTGCTCAAAGCCAAGCTTCTGAACTGGA
GCGCTGCAAAGAACAAGACCAAAGTGAGTCTTATCGACGTGACGAAGGCAGCAGAACAGGTCATCAGGGAGGACATCGATGATTTGATCTCTAAA
GTGGTTGACCCGAAAAAGGCCGACTATGCAGTTGTGACTGGCATTCAGATCCACGCTGCCTGCTCCTGCGGTAGTGGTAGCTGGACACCCGACCAC
CTTGTGGATTACATAGCCCCTGGTACCATGTATGCTGTTGTAAATGGACAACGGTACCTTCTGAAGACGGATGGCAAGAACTTCTACACGGAAAAG
GCAACTGACCCAGTTCTTACACCCGCTGCTTCGGTGAGCAAGGGCGAGGAGCTGTTCACCGGGGTGGTGCCCATCCTGGTCGAGCTGGACGGCGAC
GTAAACGGCCACAAGTTCAGCGTGTCCGGCGAGGGCGAGGGCGATGCCACCTACGGCAAGCTGACCCTGAAGCTGATCTGCACCACCGGCAAGCTG
CCCGTGCCCTGGCCCACCCTCGTGACCACCCTGGGCTACGGCCTGCAGTGCTTCGCCCGCTACCCCGACCACATGAAGCAGCACGACTTCTTCAAGT
CCGCCATGCCCGAAGGCTACGTCCAGGAGCGCACCATCTTCTTCAAGGACGACGGCAACTACAAGACCCGCGCCGAGGTGAAGTTCGAGGGCGACA
CCCTGGTGAACCGCATCGAGCTGAAGGGCATCGACTTCAAGGAGGACGGCAACATCCTGGGGCACAAGCTGGAGTACAACTACAACAGCCACAACG
TCTATATCACCGCCGACAAGCAGAAGAACGGCATCAAGGCCAACTTCAAGATCCGCCACAACATCGAGGACGGCGGCGTGCAGCTCGCCGACCACT
ACCAGCAGAACACCCCCATCGGCGACGGCCCCGTGCTGCTGCCCGACAACCACTACCTGAGCTACCAGTCCAAGCTGAGCAAAGACCCCAACGAGA
AGCGCGATCACATGGTCCTGCTGGAGTTCGTGACCGCCGCCGGGATCACTCTCGGCATGGACGAGCTGTACAAGTAAGCTTGAGCTCGAATTTCCCC
GATCGTTCAAACATTTGGCAATAAAGTTTCTTAAGATTGAATCCTGTTGCCGGTCTTGCGATGATTATCATATAATTTCTGTTGAATTACGTTAAGC
ATGTAATAATTAACATGTAATGCATGACGTTATTTATGAGATGGGTTTTTATGATTAGAGTCCCGCAATTATACATTTAATACGCGATAGAAAACAA
AATATAGCGCGCAAACTAGGATAAATTATCGCGCGCGGTGTCATCTATGTTACTAGATCGATCCGTATCGATAGCCTCTAGCTAGAGTCGATCGACAA
GCTCGAGTTTCTCCATAATAATGTGTGAGTAGTTCCCAGATAAGGGAATTAGGGTTCCTATAGGGTTTCGCTCATGTGTTGAGCATATAAGAAACCC
TTAGTATGTATTTGTATTTGTAAAATACTTCTATCAATAAAATTTCTAATTCCTAAAATCAAAATCCAGTACTAAAATCCAGATCGCTAGGT

#### >SR_L2_pRSSU_CAH3_egFP

ATACTTGAGACCGGATCCTGACAGGATATATTGGCGGGTAAACCTAAGAGAAAAGAGCGTTTATTAGAATAATCGGATATTTAAAAGGGCGTGAAA
AGGTTTATCCGTTCGTCCATTTGTATGTGCATGCCAACCACAGGGTTCCCCTCGGGATCAAAGTACTTTGATCCAACCCCTCCGCTGCTATAGTGCA
GTCGGCTTCTGACGTTCAGTGCAGCCGTCATCTGAAAACGACATGTCGCACAAGTCCTAAGTTACGCGACAGGCTGCCGCCCTGCCCTTTTCCTGGC
GTTTTCTTGTCGCGTGTTTTAGTCGCATAAAGTAGAATACTTGCGACTAGAACCGGAGACATTACGCCATGAACAAGAGCGCCGCCGCTGGCCTGC
TGGGCTATGCCCGCGTCAGCACCGACGACCAGGACTTGACCAACCAACGGGCCGAACTGCACGCGGCCGGCTGCACCAAGCTGTTTTCCGAGAAGA
TCACCGGCACCAGGCGCGACCGCCCGGAGCTGGCCAGGATGCTTGACCACCTACGCCCTGGCGACGTTGTGACAGTGACCAGGCTAGACCGCCTGGC
CCGCAGCACCCGCGACCTACTGGACATTGCCGAGCGCATCCAGGAGGCCGGCGCGGGCCTGCGTAGCCTGGCAGAGCCGTGGGCCGACACCACCACG
CCGGCCGGCCGCATGGTGTTGACCGTGTTCGCCGGCATTGCCGAGTTCGAGCGTTCCCTAATCATCGACCGCACCCGGAGCGGGCGCGAGGCCGCCA
AGGCCCGAGGCGTGAAGTTTGGCCCCCGCCCTACCCTCACCCCGGCACAGATCGCGCACGCCCGCGAGCTGATCGACCAGGAAGGCCGCACCGTGAA
AGAGGCGGCTGCACTGCTTGGCGTGCATCGCTCGACCCTGTACCGCGCACTTGAGCGCAGCGAGGAAGTGACGCCCACCGAGGCCAGGCGGCGCGG
TGCCTTCCGTGAGGACGCATTGACCGAGGCCGACGCCCTGGCGGCCGCCGAGAATGAACGCCAAGAGGAACAAGCATGAAACCGCACCAGGACGGC
CAGGACGAACCGTTTTTCATTACCGAAGAGATCGAGGCGGAGATGATCGCGGCCGGGTACGTGTTCGAGCCGCCCGCGCACCTCTCAACCGTGCGG
CTGCATGAAATCCTGGCCGGTTTGTCTGATGCCAAGCTGGCGGCCTGGCCGGCCAGCTTGGCCGCTGAAGAAACCGAGCGCCGCCGTCTAAAAAGG
TGATGTGTATTTGAGTAAAACAGCTTGCGTCATGCGGTCGCTGCGTATATGATGCGATGAGTAAATAAACAAATACGCAAGGGGAACGCATGAAGG
TTATCGCTGTACTTAACCAGAAAGGCGGGTCAGGCAAGACGACCATCGCAACCCATCTAGCCCGCGCCCTGCAACTCGCCGGGGCCGATGTTCTGTT
AGTCGATTCCGATCCCCAGGGCAGTGCCCGCGATTGGGCGGCCGTGCGGGAAGATCAACCGCTAACCGTTGTCGGCATCGACCGCCCGACGATTGAC
CGCGACGTGAAGGCCATCGGCCGGCGCGACTTCGTAGTGATCGACGGAGCGCCCCAGGCGGCGGACTTGGCTGTGTCCGCGATCAAGGCAGCCGAC
TTCGTGCTGATTCCGGTGCAGCCAAGCCCTTACGACATATGGGCCACCGCCGACCTGGTGGAGCTGGTTAAGCAGCGCATTGAGGTCACGGATGGA
AGGCTACAAGCGGCCTTTGTCGTGTCGCGGGCGATCAAAGGCACGCGCATCGGCGGTGAGGTTGCCGAGGCGCTGGCCGGGTACGAGCTGCCCATT
CTTGAGTCCCGTATCACGCAGCGCGTGAGCTACCCAGGCACTGCCGCCGCCGGCACAACCGTTCTTGAATCAGAACCCGAGGGCGACGCTGCCCGCG
AGGTCCAGGCGCTGGCCGCTGAAATTAAATCAAAACTCATTTGAGTTAATGAGGTAAAGAGAAAATGAGCAAAAGCACAAACACGCTAAGTGCCG
GCCGTCCGAGCGCACGCAGCAGCAAGGCTGCAACGTTGGCCAGCCTGGCAGACACGCCAGCCATGAAGCGGGTCAACTTTCAGTTGCCGGCGGAGG
ATCACACCAAGCTGAAGATGTACGCGGTACGCCAAGGCAAGACCATTACCGAGCTGCTATCTGAATACATCGCGCAGCTACCAGAGTAAATGAGCAA
ATGAATAAATGAGTAGATGAATTTTAGCGGCTAAAGGAGGCGGCATGGAAAATCAAGAACAACCAGGCACCGACGCCGTGGAATGCCCCATGTGTG
GAGGAACGGGCGGTTGGCCAGGCGTAAGCGGCTGGGTTGCCTGCCGGCCCTGCAATGGCACTGGAACCCCCAAGCCCGAGGAATCGGCGTGAGCGG
TCGCAAACCATCCGGCCCGGTACAAATCGGCGCGGCGCTGGGTGATGACCTGGTGGAGAAGTTGAAGGCCGCGCAGGCCGCCCAGCGGCAACGCAT
CGAGGCAGAAGCACGCCCCGGTGAATCGTGGCAAGCGGCCGCTGATCGAATCCGCAAAGAATCCCGGCAACCGCCGGCAGCCGGTGCGCCGTCGAT
TAGGAAGCCGCCCAAGGGCGACGAGCAACCAGATTTTTTCGTTCCGATGCTCTATGACGTGGGCACCCGCGATAGTCGCAGCATCATGGACGTGGC
CGTTTTCCGTCTGTCGAAGCGTGACCGACGAGCTGGCGAGGTGATCCGCTACGAGCTTCCAGACGGGCACGTAGAGGTTTCCGCAGGGCCGGCCGG
CATGGCCAGTGTGTGGGATTACGACCTGGTACTGATGGCGGTTTCCCATCTAACCGAATCCATGAACCGATACCGGGAAGGGAAGGGAGACAAGCC
CGGCCGCGTGTTCCGTCCACACGTTGCGGACGTACTCAAGTTCTGCCGGCGAGCCGATGGCGGAAAGCAGAAAGACGACCTGGTAGAAACCTGCAT
TCGGTTAAACACCACGCACGTTGCCATGCAGCGTACGAAGAAGGCCAAGAACGGCCGCCTGGTGACGGTATCCGAGGGTGAAGCCTTGATTAGCCG
CTACAAGATCGTAAAGAGCGAAACCGGGCGGCCGGAGTACATCGAGATCGAGCTAGCTGATTGGATGTACCGCGAGATCACAGAAGGCAAGAACCC
GGACGTGCTGACGGTTCACCCCGATTACTTTTTGATCGATCCCGGCATCGGCCGTTTTCTCTACCGCCTGGCACGCCGCGCCGCAGGCAAGGCAGAA
GCCAGATGGTTGTTCAAGACGATCTACGAACGCAGTGGCAGCGCCGGAGAGTTCAAGAAGTTCTGTTTCACCGTGCGCAAGCTGATCGGGTCAAAT
GACCTGCCGGAGTACGATTTGAAGGAGGAGGCGGGGCAGGCTGGCCCGATCCTAGTCATGCGCTACCGCAACCTGATCGAGGGCGAAGCATCCGCC
GGTTCCTAATGTACGGAGCAGATGCTAGGGCAAATTGCCCTAGCAGGGGAAAAAGGTCGAAAAGGACTCTTTCCTGTGGATAGCACGTACATTGGG
AACCCAAAGCCGTACATTGGGAACCGGAACCCGTACATTGGGAACCCAAAGCCGTACATTGGGAACCGGTCACACATGTAAGTGACTGATATAAAA
GAGAAAAAAGGCGATTTTTCCGCCTAAAACTCTTTAAAACTTATTAAAACTCTTAAAACCCGCCTGGCCTGTGCATAACTGTCTGGCCAGCGCACA
GCCGAAGTGCTGCAAAAAGCGCCTACCCTTCGGTCGCTGCGCTCCCTACGCCCCGCCGCTTCGCGTCGGCCTATCGCGGCCGCTGGCCGCTCAAAAA
TGGCTGGCCTACGGCCAGGCAATCTACCAGGGCGCGGACAAGCCGCGCCGTCGCCACTCGACCGCCGGCGCCCACATCAAGGCACCCTGCCTCGCGC

GTTTCGGTGATGACGGTGAAAACCTCTGACACATGCAGCTCCCGGTGACGGTCACAGCTTGTCTGTAAGCGGATGCCGGGAGCAGACAAGCCCGTC
AGGGCGCGTCAGCGGGTGTTGGCGGGTGTCGGGGCGCAGCCATGACCCAGTCACGTAGCGATAGCGGAGTGTATACTGGCTTAACTATGCGGCATC
AGAGCAGATTGTACTGAGAGTGCACCATATGCGGTGTGAAATACCGCACAGATGCGTAAGGAGAAAATACCGCATCAGGCGCTCATCCGCTTCCTCG
CTCACTGACTCGCTGCGCTCGGTCGTTCGGCTGCGGCGAGCGGTATCAGCTCACTCAAAGGCGGTAATACGGTTATCCACAGAATCAGGGGATAACG
CAGGAAAGAACATGTGAGCAAAAGGCCAGCAAAAGGCCAGGAACCGTAAAAAGGCCGCGTTGCTGGCGTTTTTCCATAGGCTCCGCCCCCCTGAC
GAGCATCACAAAAATCGACGCTCAAGTCAGAGGTGGCGAAACCCGACAGGACTATAAAGATACCAGGCGTTTCCCCCTGGAAGCTCCCTCGTGCGC
TCTCCTGTTCCGACCCTGCCGCTTACCGGATACCTGTCCGCCTTTCTCCCTTCGGGAAGCGTGGCGCTTTCTCATAGCTCACGCTGTAGGTATCTCAG
TTCGGTGTAGGTCGTTCGCTCCAAGCTGGGCTGTGTGCACGAACCCCCCGTTCAGCCCGACCGCTGCGCCTTATCCGGTAACTATCGTCTTGAGTCC
AACCCGGTAAGACACGACTTATCGCCACTGGCAGCAGCCACTGGTAACAGGATTAGCAGAGCGAGGTATGTAGGCGGTGCTACAGAGTTCTTGAAG
TGGTGGCCTAACTACGGCTACACTAGAAGGACAGTATTTGGTATCTGCGCTCTGCTGAAGCCAGTTACCTTCGGAAAAAGAGTTGGTAGCTCTTGA
TCCGGCAAACAAACCACCGCTGGTAGCGGTGGTTTTTTTGTTTGCAAGCAGCAGATTACGCGCAGAAAAAAAGGATCTCAAGAAGATCCTTTGAT
CTTTTCTACGGGGTCTGACGCTCAGTGGAACGAAAACTCACGTTAAGGGATTTTGGTCATGCATTCTAGGTGATTATTTGCCGACTACCTTGGTGA
TCTCGCCTTTCACGTAGTGGACAAATTCTTCCAACTGATCTGCGCGCGAGGCCAAGCGATCTTCTTCTTGTCCAAGATAAGCCTGTCTAGCTTCAAG
TATGACGGGCTGATACTGGGCCGGCAGGCGCTCCATTGCCCAGTCGGCAGCGACATCCTTCGGCGCGATTTTGCCGGTTACTGCGCTGTACCAAATG
CGGGACAACGTAAGCACTACATTTCGCTCATCACCAGCCCAGTCGGGCGGCGAGTTCCATAGCGTTAAGGTTTCATTTAGCGCCTCAAATAGATCCT
GTTCAGGAACCGGATCAAAGAGTTCCTCCGCCGCTGGACCTACCAAGGCAACGCTATGTTCTCTTGCTTTTGTCAGCAAGATAGCCAGATCAATGT
CGATCGTGGCTGGCTCGAAGATACCTGCAAGAATGTCATTGCGCTGCCATTCTCCAAATTGCAGTTCGCGCTTAGCTGGATAACGCCACGGAATGAT
GTCGTCGTGCACAACAATGGTGACTTCTACAGCGCGGAGAATCTCGCTCTCTCCAGGGGAAGCCGAAGTTTCCAAAAGGTCGTTGATCAAAGCTCG
CCGCGTTGTTTCATCAAGCCTTACGGTCACCGTAACCAGCAAATCAATATCACTGTGTGGCTTCAGGCCGCCATCCACTGCGGAGCCGTACAAATGT
ACGGCCAGCAACGTCGGTTCGAGATGGCGCTCGATGACGCCAACTACCTCTGATAGTTGAGTCGATACTTCGGCGATCACCGCTTCCCTCATAATGT
TTAACTTTGTTTTAGGGCGACTGCCCTGCTGCGTAACATCGTTGCTGCTCCATAACATCAAACATCGACCCACGGCGTAACGCGCTTGCTGCTTGGA
TGCCCGAGGCATAGACTGTACCCCAAAAAAACAGTCATAACAAGCCATGAAAACCGCCACTGCGCCGTTACCACCGCTGCGTTCGGTCAAGGTTCT
GGACCAGTTGCGTGAGCGCATACGCTACTTGCATTACAGCTTACGAACCGAACAGGCTTATGTCCACTGGGTTCGTGCCTTCATCCGTTTCCACGGT
GTGCGTCACCCGGCAACCTTGGGTAGCAGCGAAGTCGAGGCATTTCTGTCCTGGCTGGAACAGAACTTATTATTTCCTTCCTCTTTTCTACAGTATT
TAAAGATACCCCAAGAAGCTAATTATAACAAGACGAACTCCAATTCACTGTTCCTTGCATTCTAAAACCTTAAATACCAGAAAACAGCTTTTTCAA
AGTTGTTTTCAAAGTTGGCGTATAACATAGTATCGACGGAGCCGATTTTGAAACCGCGGTGATCACAGGCAGCAACGCTCTGTCATCGTTACAATC
AACATGCTACCCTCCGCGAGATCATCCGTGTTTCAAACCCGGCAGCTTAGTTGCCGTTCTTCCGAATAGCATCGGTAACATGAGCAAAGTCTGCCGC
CTTACAACGGCTCTCCCGCTGACGCCGTCCCGGACTGATGGGCTGCCTGTATCGAGTGGTGATTTTGTGCCGAGCTGCCGGTCGGGGAGCTGTTGG
CTGGCTGGTGGCAGGATATATTGTGGTGTAAACATAACGGATCCGGTCTCAGGAGAATGAGGAGCACCGTCCAAACCTGGATAGCCGATATGTGAG
TGTTTGGTTGCCTGTCATTTCTAAAAAGCGTATGTCTCGTAGCGTAGATAGCAAGCGCTGTGAGAAAGGTGTGCTCCTATAGCATGGTAGCATCTGT
CCCATATCTCGTCCGCGAGAAGCCTACATTTCTCGTTGCGTATTAGAGCATGAAAAACCAACGTGAAGTACGTGATTCAGACCTTGACCAGTTCCAG
TTCTCTTCAACAAAAAAACAGTTCTTCAATTCATTTGCTTACATTAAACAATTGTCACACTCTTCTTCCCGGATGACAGCACCAAGACACTTCCCC
TCCACAAAAATCTCCGAGTCATGTGGACCGGCCTGGATGCTCTTAAAGGGGTTGCCATTGTAGCGGGGTGCCTTAACCACGACCTCTTCGATACCTG
TCTGAAGCGGCGTGTGAAATCCTCCAGGAGCTTGTTCAAATTTACAAGCTGCAATGGATGTAAACTCATCCAAAGACATGACTTGCGAGAAGCCAG
AAACGGTCATCAACACTACGAACACCCCTAAACATGGATGGAGAGGCAACATCCTGCATGTGTCTATGTCAGAGGATGCAAAAGTGAAAGACAAAG
GAGAAGGATTGCTCGTGCAATCACTGCAAAGCAAGATGCGAATATCCCTAAAATCCACAGCAGGAAAAGAAAGGAGCCTCAATAAGTCTCTAAAAT
GGGAAGAAAAGAAGCACGGGCCAGCGAGACGGAAATGAGGCGATCCGATGCGATGGTTCCGAGAACTGAAGCGGTGCGTGGGGGGCTAGAAATGA
ATGGGGAACGTGGAGGGCAAACAAAGGCTGTACTTAGCGGCAAATGCAGATGGCATGCATGCCAAGAACAGCAGCAGTCGGCACCGCCAGTTTAG
CGGAGAGCAACAGGCAGCAAAAGGAAGTGGAACAGGTTTAGGGCCGACGCAGTCGGCGCCTGCCCTAAAACGGCAAGGCCCACGGCAGCGGTCGC
GAGGGCGTTGCCTGCCGGTCGTGGAAAGCCGTGGGGCTGTCAACGGAACCTCACGCTCCCAGCCTGCCCTAAGCGGGGAAAAAGAAGATCCACATT
CTAGGGCTTTCCATCTCGGCTCTCAGCCGTCCGATTAAGAATCGCACGGCCTAGCCCCGCCAGCCTAGCCCTCGGCCGCCCCCTTGACAGGTTAAGA
GCTGGGCAAGCGACTCAGTTGTTCCTTCGTCGCTCGCCCTTCTCACGAGCAGCTCGCCTGCCCCGCCGCCGCCTTGCCTGCCTCGCGCCTTTTGTCT
CCAGGTTCCCTTGCCGCGCCTTCTCCTTGGCTTGTGCTTCTGCTTTGCTTGTGCCTCTGCTGGGCTCTACTTGTGTGGTGTTGTGCTTTGGCGCCGC
CGCCTCGCTGCTGCTGTGGAGCTTGGTGTGTGCTGGGGTTTCTGGGGTTTGCATGTGTCAGGGTTTGTAGATCTGTTCTTCGGGGCCGTCTTTCTA
ACGTTTCGTGTGCCGTTGTTTCTGGGGGGTTTGGTGTGGAGCTCGCAGCTTGCAGCTAATGAAAAAGCCTGAACTCACCGCGACGTCTGTCGAGAA
GTTTCTGATCGAAAAGTTCGACAGCGTCTCCGACCTGATGCAGCTCTCGGAGGGCGAAGAATCTCGTGCTTTCAGCTTCGATGTAGGAGGGCGTGG
ATATGTCCTGCGGGTAAATAGCTGCGCCGATGGTTTCTACAAAGATCGTTATGTTTATCGGCACTTTGCATCGGCCGCGCTCCCGATTCCGGAAGTG
CTTGACATTGGGGAGTTTAGCGAGAGCCTGACCTATTGCATCTCCCGCCGTTCACAGGGTGTCACGTTGCAAGACCTGCCTGAAACCGAACTGCCC
GCTGTTCTACAACCGGTCGCGGAGGCTATGGATGCGATCGCTGCGGCCGATCTTAGCCAGACGAGCGGGTTCGGCCCATTCGGACCGCAAGGAATC
GGTCAATACACTACATGGCGTGATTTCATATGCGCGATTGCTGATCCCCATGTGTATCACTGGCAAACTGTGATGGACGACACCGTCAGTGCGTCCG
TCGCGCAGGCTCTCGATGAGCTGATGCTTTGGGCCGAGGACTGCCCCGAAGTCCGGCACCTCGTGCACGCGGATTTCGGCTCCAACAATGTCCTGA
CGGACAATGGCCGCATAACAGCGGTCATTGACTGGAGCGAGGCGATGTTCGGGGATTCCCAATACGAGGTCGCCAACATCTTCTTCTGGAGGCCGT
GGTTGGCTTGTATGGAGCAGCAGACGCGCTACTTCGAGCGGAGGCATCCGGAGCTTGCAGGATCGCCACGACTCCGGGCGTATATGCTCCGCATTG
GTCTTGACCAACTCTATCAGAGCTTGGTTGACGGCAATTTCGATGATGCAGCTTGGGCGCAGGGTCGATGCGACGCAATCGTCCGATCCGGAGCCG
GGACTGTCGGGCGTACACAAATCGCCCGCAGAAGCGCGGCCGTCTGGACCGATGGCTGTGTAGAAGTACTCGCCGATAGTGGAAACCGACGCCCCA
GCACTCGTCCGAGGGCAAAGAAATAGGCTTGAGCTCGAATTTCCCCGATCGTTCAAACATTTGGCAATAAAGTTTCTTAAGATTGAATCCTGTTGC
CGGTCTTGCGATGATTATCATATAATTTCTGTTGAATTACGTTAAGCATGTAATAATTAACATGTAATGCATGACGTTATTTATGAGATGGGTTTTT
ATGATTAGAGTCCCGCAATTATACATTTAATACGCGATAGAAAACAAAATATAGCGCGCAAACTAGGATAAATTATCGCGCGCGGTGTCATCTATGT
TACTAGATCGATCCGTATCGATAGCCTCTAGCTAGAGTCGATCGACAAGCTCGAGTTTCTCCATAATAATGTGTGAGTAGTTCCCAGATAAGGGAAT
TAGGGTTCCTATAGGGTTTCGCTCATGTGTTGAGCATATAAGAAACCCTTAGTATGTATTTGTATTTGTAAAATACTTCTATCAATAAAATTTCTAAT
TCCTAAAATCAAAATCCAGTACTAAAATCCAGATCGCTAGCAAGGAGAGTTCATCTGTCAGCACTTGGACAGACCTTCCGGATCAAAGGTCGCTAT
TGCTATTCGCAGATCTAAAGTTTGAACGGCATTCACCGAGTGGTGGATGCTTCAACTGGCTTCTGGCACCGAGCGTAAAAGGACAAATCCAACGTA
CATATCGCTAAATTTCGTTCCAACGTACATATCGCTAATTTTTCCTACGGGAGGAATTAATCCTCGATAAACTCGGTGCCTTCTCGAAAGATGGTTT
CACCATGGTTCAACTCCATGGCTCACCCCGACCCCACGCTTATCCCACGGCTTCCAACACTCAGGGTGCTCGACTCCTATTACTCCCCGACCGTCTTC
TTCGGAAAATCTTTATGCGCATTCCCCATCCTAACTTGGAGATGCGATTGCTTCATCCGCCTTCCTGTTCATCACTTCTCATCACCCCTCATTCGCA
AAGAAATATCGTGGCACCTGGGATATGGAAATTTGCCCACTCTGCCCTAGACTCTGAAAATCAGGGATGAACAAAAGGAATGACCGACAATCTTTG
TGGAAAAGCTCTGGACAGGACTAGACTACCAATTGTAACCTGAAACCTTGCTCTTTGAATGGGGAAATTCCTTACATGAGTCCTAATTGAGAAATC
TGTCGCCCTACAAAGTTGCAAATGGCAGCTCTGGCTATGGCAGCTCTAGCAATGGCCAGGCAAATGGCAGCTCTAGCAATGGCCAGGCAAATGGCA
GCTCTAGCTATGGCCAGGCGGGCAAGGCCGTGATCTTCTTCCAGAGCGAGCTCGGGCATACAGGCGGGGCCAGCATAGAGGGCTCACTATTTCCTCC
AGCTGGGAGGCCTAGGGAAGGAAGGTCTGTGCCGCTGGCTATGGCGGTCGACTTGTGTGGGCCCTCCGAAGAACGAGCCAAAACTCTGAGCCAGTC
GTTGCGCAGCCAATGAGCGGAGCTGTGGATGCTTGAACTGCGTTTTGTCATTCGATTATGTATTGATTAGCAACTCAAATGATAAGTCGATCTCTC
AGGTAATACATAGTCCATGGTTTGAGTTCAAATTAGTCGCTCAAAAAACGAGGCTTTGGAGAAATCCAGCGAGGTTGGCGATCCTTTCTTATCCTT
CCCTGCCAACTCTCTTCGTGTGGCCGTGAACGCTCTCTCGTTGCCCCTATCCTATTTGAGCCCGGGCGGGCCACTCTCCAATCCAGTCCGCCCTCTCC
CGGCCCTCTCCTTTTTCTTCCCCGGTAGCAGGCATACATCTTGCGCTTCTCCGGTTCCATCCACACTGCAGCAGCAATGGGGAGCCTGCGGGTGCTG

GTGCCCAGCACATTGCTGGGAAGCAACGGCCAAGCGGAGTCCTGGAGGGCCGATTTAGGCACTGCCCGGCCTGCGCTTCCTCGCAGGGGCGGGAAG
GGGAATGCGGGGGGAGTGAGATGCTGCGTGCGGGAGGCGGGGCCGGCGGACAGGGAAGAAGGTGAGGGCTCCAAGAGCAGCCTAATTGCGCGCGA
TGGCTTGGGGAGGAGAGGGATGCTGTCCGCGGCTATGCTGGCCACCATGACGTGCGTGTGCACCGAGGGGGCGGAGGCCGCAGTGGCGCGGCACGC
GGCCGTGGAGCAGAGGCGTCGGGAGGCCCTGCAGCTGCAGACCCTGCAGGGGGCCAAGGCGCGCGGGGGATCTCGGATCAAAAAGCTCGGGGGCA
AGGCTGCCGTCGGCACCGCCGAGTGGGAGTACGGCAATTCCTGCGGGCCGGGCGAATGGGGCTCTGTGTGCGCCGCGGGCCGCGTGCAGTCGCCGG
TGAACTTCGAGATGAACAAGGTGCAGGAGAAGGGGGAGAAGGCCATGAGTGATCTCATGTTCGACTACGGCCCCGTCCAGCCCACCTTCCTCAATA
CGGGACACGGTACAATGCAGGTAAATTTTCCAGGTGGTCGCAATAAGCTAAAGATTGGCGACAGGGTGCTGGACCTGCTCCAGTTTCACTTCCACA
CGCCGTCTGAGCATTCCTTCAACGGGGTGCACCACACCATGGAGGCTCACTTGGTGCACGGTGACCCTAAAACCAAGTCGCTGGCAGTGGTCGGGG
TGCTGCTCGATGCCAAGAGCCGCGTCAGCGCCAACAAGGCTTTGCAAGCTGCACTGGAGTACTCGCCCAAGGAGCACTATAAGACAGCGCGAGGAC
CGGATAACTTCAGTCTATCGCCCTCCCTCCTGCTCCCGTATGCAGGAAGAATTCGGAAGAAGCGGGGATATATGTACTACCAGGGCTCTCTCACGAC
TCCTCCTTGCTCCGAGGGTGTGGAGTGGTATGTCATGGAAACTCCGGTGAGCATCTCCTACGCGCAGGTGGTGGAGTTCATGCTATACGTTGGAGAC
TCCAGAACTTTAGCCTTGAACACCAGGCCCGTCCAGCCCCTGGGCCAGCGGAGGGTATACAGGGGCCCCATGCAGGCGGCTTCGGTGAGCAAGGGC
GAGGAGCTGTTCACCGGGGTGGTGCCCATCCTGGTCGAGCTGGACGGCGACGTAAACGGCCACAAGTTCAGCGTGTCCGGCGAGGGCGAGGGCGAT
GCCACCTACGGCAAGCTGACCCTGAAGTTCATCTGCACCACCGGCAAGCTGCCCGTGCCCTGGCCCACCCTCGTGACCACCCTGACCTACGGCGTGC
AGTGCTTCAGCCGCTACCCCGACCACATGAAGCAGCACGACTTCTTCAAGTCCGCCATGCCCGAAGGCTACGTCCAGGAGCGCACCATCTTCTTCAA
GGACGACGGCAACTACAAGACCCGCGCCGAGGTGAAGTTCGAGGGCGACACCCTGGTGAACCGCATCGAGCTGAAGGGCATCGACTTCAAGGAGG
ACGGCAACATCCTGGGGCACAAGCTGGAGTACAACTACAACAGCCACAACGTCTATATCATGGCCGACAAGCAGAAGAACGGCATCAAGGTGAACT
TCAAGATCCGCCACAACATCGAGGACGGCAGCGTGCAGCTCGCCGACCACTACCAGCAGAACACCCCCATCGGCGACGGCCCCGTGCTGCTGCCCGA
CAACCACTACCTGAGCACCCAGTCCGCCCTGAGCAAAGACCCCAACGAGAAGCGCGATCACATGGTCCTGCTGGAGTTCGTGACCGCCGCCGGGATC
ACTCTCGGCATGGACGAGCTGTACAAGTAAGCTTGAGCTCGAATTTCCCCGATCGTTCAAACATTTGGCAATAAAGTTTCTTAAGATTGAATCCTG
TTGCCGGTCTTGCGATGATTATCATATAATTTCTGTTGAATTACGTTAAGCATGTAATAATTAACATGTAATGCATGACGTTATTTATGAGATGGGT
TTTTATGATTAGAGTCCCGCAATTATACATTTAATACGCGATAGAAAACAAAATATAGCGCGCAAACTAGGATAAATTATCGCGCGCGGTGTCATCT
ATGTTACTAGATCGATCCGTATCGATAGCCTCTAGCTAGAGTCGATCGACAAGCTCGAGTTTCTCCATAATAATGTGTGAGTAGTTCCCAGATAAGG
GAATTAGGGTTCCTATAGGGTTTCGCTCATGTGTTGAGCATATAAGAAACCCTTAGTATGTATTTGTATTTGTAAAATACTTCTATCAATAAAATTT
CTAATTCCTAAAATCAAAATCCAGTACTAAAATCCAGATCGCTATACAGGAGTCTGAGCAGGACAACTCGCGTAGTGAGAGTTACATGTTCGTTGGG
TTCTTCCGACACGGACCTGAGTTGGCCAACGTCCCACCTGAGGTCTGTGCCCCGGTGATGAGAAGTGTGCATCTCGTTCTTGCAGCTCGTCAGTACT
TTCAGAATCATGGCGTGCATGGTAGAATGACCCTTATAACGGACTTCGACATGGCAATCGCTACAGAGGAGTCTGAGCAGGACAACTCGCGTAGTG
AGAGTTACATGTTCGTTGGGTTCTTCCGACACGGACCTGAGTTGGCCAACGTCCCACCTGAGGTCTGTGCCCCGGTGATGAGAAGTGTGCATCTCG
TTCTTGCAGCTCGTCAGTACTTTCAGAATCATGGCGTGCATGGTAGAATGACCCTTATAACGGACTTCGACATGGCAATCGCTAGGT

#### >SR_L2_pRSSU_RBMP1_eGFP

ATACTTGAGACCGGATCCTGACAGGATATATTGGCGGGTAAACCTAAGAGAAAAGAGCGTTTATTAGAATAATCGGATATTTAAAAGGGCGTGAAA
AGGTTTATCCGTTCGTCCATTTGTATGTGCATGCCAACCACAGGGTTCCCCTCGGGATCAAAGTACTTTGATCCAACCCCTCCGCTGCTATAGTGCA
GTCGGCTTCTGACGTTCAGTGCAGCCGTCATCTGAAAACGACATGTCGCACAAGTCCTAAGTTACGCGACAGGCTGCCGCCCTGCCCTTTTCCTGGC
GTTTTCTTGTCGCGTGTTTTAGTCGCATAAAGTAGAATACTTGCGACTAGAACCGGAGACATTACGCCATGAACAAGAGCGCCGCCGCTGGCCTGC
TGGGCTATGCCCGCGTCAGCACCGACGACCAGGACTTGACCAACCAACGGGCCGAACTGCACGCGGCCGGCTGCACCAAGCTGTTTTCCGAGAAGA
TCACCGGCACCAGGCGCGACCGCCCGGAGCTGGCCAGGATGCTTGACCACCTACGCCCTGGCGACGTTGTGACAGTGACCAGGCTAGACCGCCTGGC
CCGCAGCACCCGCGACCTACTGGACATTGCCGAGCGCATCCAGGAGGCCGGCGCGGGCCTGCGTAGCCTGGCAGAGCCGTGGGCCGACACCACCACG
CCGGCCGGCCGCATGGTGTTGACCGTGTTCGCCGGCATTGCCGAGTTCGAGCGTTCCCTAATCATCGACCGCACCCGGAGCGGGCGCGAGGCCGCCA
AGGCCCGAGGCGTGAAGTTTGGCCCCCGCCCTACCCTCACCCCGGCACAGATCGCGCACGCCCGCGAGCTGATCGACCAGGAAGGCCGCACCGTGAA
AGAGGCGGCTGCACTGCTTGGCGTGCATCGCTCGACCCTGTACCGCGCACTTGAGCGCAGCGAGGAAGTGACGCCCACCGAGGCCAGGCGGCGCGG
TGCCTTCCGTGAGGACGCATTGACCGAGGCCGACGCCCTGGCGGCCGCCGAGAATGAACGCCAAGAGGAACAAGCATGAAACCGCACCAGGACGGC
CAGGACGAACCGTTTTTCATTACCGAAGAGATCGAGGCGGAGATGATCGCGGCCGGGTACGTGTTCGAGCCGCCCGCGCACCTCTCAACCGTGCGG
CTGCATGAAATCCTGGCCGGTTTGTCTGATGCCAAGCTGGCGGCCTGGCCGGCCAGCTTGGCCGCTGAAGAAACCGAGCGCCGCCGTCTAAAAAGG
TGATGTGTATTTGAGTAAAACAGCTTGCGTCATGCGGTCGCTGCGTATATGATGCGATGAGTAAATAAACAAATACGCAAGGGGAACGCATGAAGG
TTATCGCTGTACTTAACCAGAAAGGCGGGTCAGGCAAGACGACCATCGCAACCCATCTAGCCCGCGCCCTGCAACTCGCCGGGGCCGATGTTCTGTT
AGTCGATTCCGATCCCCAGGGCAGTGCCCGCGATTGGGCGGCCGTGCGGGAAGATCAACCGCTAACCGTTGTCGGCATCGACCGCCCGACGATTGAC
CGCGACGTGAAGGCCATCGGCCGGCGCGACTTCGTAGTGATCGACGGAGCGCCCCAGGCGGCGGACTTGGCTGTGTCCGCGATCAAGGCAGCCGAC
TTCGTGCTGATTCCGGTGCAGCCAAGCCCTTACGACATATGGGCCACCGCCGACCTGGTGGAGCTGGTTAAGCAGCGCATTGAGGTCACGGATGGA
AGGCTACAAGCGGCCTTTGTCGTGTCGCGGGCGATCAAAGGCACGCGCATCGGCGGTGAGGTTGCCGAGGCGCTGGCCGGGTACGAGCTGCCCATT
CTTGAGTCCCGTATCACGCAGCGCGTGAGCTACCCAGGCACTGCCGCCGCCGGCACAACCGTTCTTGAATCAGAACCCGAGGGCGACGCTGCCCGCG
AGGTCCAGGCGCTGGCCGCTGAAATTAAATCAAAACTCATTTGAGTTAATGAGGTAAAGAGAAAATGAGCAAAAGCACAAACACGCTAAGTGCCG
GCCGTCCGAGCGCACGCAGCAGCAAGGCTGCAACGTTGGCCAGCCTGGCAGACACGCCAGCCATGAAGCGGGTCAACTTTCAGTTGCCGGCGGAGG
ATCACACCAAGCTGAAGATGTACGCGGTACGCCAAGGCAAGACCATTACCGAGCTGCTATCTGAATACATCGCGCAGCTACCAGAGTAAATGAGCAA
ATGAATAAATGAGTAGATGAATTTTAGCGGCTAAAGGAGGCGGCATGGAAAATCAAGAACAACCAGGCACCGACGCCGTGGAATGCCCCATGTGTG
GAGGAACGGGCGGTTGGCCAGGCGTAAGCGGCTGGGTTGCCTGCCGGCCCTGCAATGGCACTGGAACCCCCAAGCCCGAGGAATCGGCGTGAGCGG
TCGCAAACCATCCGGCCCGGTACAAATCGGCGCGGCGCTGGGTGATGACCTGGTGGAGAAGTTGAAGGCCGCGCAGGCCGCCCAGCGGCAACGCAT
CGAGGCAGAAGCACGCCCCGGTGAATCGTGGCAAGCGGCCGCTGATCGAATCCGCAAAGAATCCCGGCAACCGCCGGCAGCCGGTGCGCCGTCGAT
TAGGAAGCCGCCCAAGGGCGACGAGCAACCAGATTTTTTCGTTCCGATGCTCTATGACGTGGGCACCCGCGATAGTCGCAGCATCATGGACGTGGC
CGTTTTCCGTCTGTCGAAGCGTGACCGACGAGCTGGCGAGGTGATCCGCTACGAGCTTCCAGACGGGCACGTAGAGGTTTCCGCAGGGCCGGCCGG
CATGGCCAGTGTGTGGGATTACGACCTGGTACTGATGGCGGTTTCCCATCTAACCGAATCCATGAACCGATACCGGGAAGGGAAGGGAGACAAGCC
CGGCCGCGTGTTCCGTCCACACGTTGCGGACGTACTCAAGTTCTGCCGGCGAGCCGATGGCGGAAAGCAGAAAGACGACCTGGTAGAAACCTGCAT
TCGGTTAAACACCACGCACGTTGCCATGCAGCGTACGAAGAAGGCCAAGAACGGCCGCCTGGTGACGGTATCCGAGGGTGAAGCCTTGATTAGCCG
CTACAAGATCGTAAAGAGCGAAACCGGGCGGCCGGAGTACATCGAGATCGAGCTAGCTGATTGGATGTACCGCGAGATCACAGAAGGCAAGAACCC
GGACGTGCTGACGGTTCACCCCGATTACTTTTTGATCGATCCCGGCATCGGCCGTTTTCTCTACCGCCTGGCACGCCGCGCCGCAGGCAAGGCAGAA
GCCAGATGGTTGTTCAAGACGATCTACGAACGCAGTGGCAGCGCCGGAGAGTTCAAGAAGTTCTGTTTCACCGTGCGCAAGCTGATCGGGTCAAAT
GACCTGCCGGAGTACGATTTGAAGGAGGAGGCGGGGCAGGCTGGCCCGATCCTAGTCATGCGCTACCGCAACCTGATCGAGGGCGAAGCATCCGCC
GGTTCCTAATGTACGGAGCAGATGCTAGGGCAAATTGCCCTAGCAGGGGAAAAAGGTCGAAAAGGACTCTTTCCTGTGGATAGCACGTACATTGGG
AACCCAAAGCCGTACATTGGGAACCGGAACCCGTACATTGGGAACCCAAAGCCGTACATTGGGAACCGGTCACACATGTAAGTGACTGATATAAAA
GAGAAAAAAGGCGATTTTTCCGCCTAAAACTCTTTAAAACTTATTAAAACTCTTAAAACCCGCCTGGCCTGTGCATAACTGTCTGGCCAGCGCACA
GCCGAAGTGCTGCAAAAAGCGCCTACCCTTCGGTCGCTGCGCTCCCTACGCCCCGCCGCTTCGCGTCGGCCTATCGCGGCCGCTGGCCGCTCAAAAA
TGGCTGGCCTACGGCCAGGCAATCTACCAGGGCGCGGACAAGCCGCGCCGTCGCCACTCGACCGCCGGCGCCCACATCAAGGCACCCTGCCTCGCGC
GTTTCGGTGATGACGGTGAAAACCTCTGACACATGCAGCTCCCGGTGACGGTCACAGCTTGTCTGTAAGCGGATGCCGGGAGCAGACAAGCCCGTC
AGGGCGCGTCAGCGGGTGTTGGCGGGTGTCGGGGCGCAGCCATGACCCAGTCACGTAGCGATAGCGGAGTGTATACTGGCTTAACTATGCGGCATC

AGAGCAGATTGTACTGAGAGTGCACCATATGCGGTGTGAAATACCGCACAGATGCGTAAGGAGAAAATACCGCATCAGGCGCTCATCCGCTTCCTCG
CTCACTGACTCGCTGCGCTCGGTCGTTCGGCTGCGGCGAGCGGTATCAGCTCACTCAAAGGCGGTAATACGGTTATCCACAGAATCAGGGGATAACG
CAGGAAAGAACATGTGAGCAAAAGGCCAGCAAAAGGCCAGGAACCGTAAAAAGGCCGCGTTGCTGGCGTTTTTCCATAGGCTCCGCCCCCCTGAC
GAGCATCACAAAAATCGACGCTCAAGTCAGAGGTGGCGAAACCCGACAGGACTATAAAGATACCAGGCGTTTCCCCCTGGAAGCTCCCTCGTGCGC
TCTCCTGTTCCGACCCTGCCGCTTACCGGATACCTGTCCGCCTTTCTCCCTTCGGGAAGCGTGGCGCTTTCTCATAGCTCACGCTGTAGGTATCTCAG
TTCGGTGTAGGTCGTTCGCTCCAAGCTGGGCTGTGTGCACGAACCCCCCGTTCAGCCCGACCGCTGCGCCTTATCCGGTAACTATCGTCTTGAGTCC
AACCCGGTAAGACACGACTTATCGCCACTGGCAGCAGCCACTGGTAACAGGATTAGCAGAGCGAGGTATGTAGGCGGTGCTACAGAGTTCTTGAAG
TGGTGGCCTAACTACGGCTACACTAGAAGGACAGTATTTGGTATCTGCGCTCTGCTGAAGCCAGTTACCTTCGGAAAAAGAGTTGGTAGCTCTTGA
TCCGGCAAACAAACCACCGCTGGTAGCGGTGGTTTTTTTGTTTGCAAGCAGCAGATTACGCGCAGAAAAAAAGGATCTCAAGAAGATCCTTTGAT
CTTTTCTACGGGGTCTGACGCTCAGTGGAACGAAAACTCACGTTAAGGGATTTTGGTCATGCATTCTAGGTGATTATTTGCCGACTACCTTGGTGA
TCTCGCCTTTCACGTAGTGGACAAATTCTTCCAACTGATCTGCGCGCGAGGCCAAGCGATCTTCTTCTTGTCCAAGATAAGCCTGTCTAGCTTCAAG
TATGACGGGCTGATACTGGGCCGGCAGGCGCTCCATTGCCCAGTCGGCAGCGACATCCTTCGGCGCGATTTTGCCGGTTACTGCGCTGTACCAAATG
CGGGACAACGTAAGCACTACATTTCGCTCATCACCAGCCCAGTCGGGCGGCGAGTTCCATAGCGTTAAGGTTTCATTTAGCGCCTCAAATAGATCCT
GTTCAGGAACCGGATCAAAGAGTTCCTCCGCCGCTGGACCTACCAAGGCAACGCTATGTTCTCTTGCTTTTGTCAGCAAGATAGCCAGATCAATGT
CGATCGTGGCTGGCTCGAAGATACCTGCAAGAATGTCATTGCGCTGCCATTCTCCAAATTGCAGTTCGCGCTTAGCTGGATAACGCCACGGAATGAT
GTCGTCGTGCACAACAATGGTGACTTCTACAGCGCGGAGAATCTCGCTCTCTCCAGGGGAAGCCGAAGTTTCCAAAAGGTCGTTGATCAAAGCTCG
CCGCGTTGTTTCATCAAGCCTTACGGTCACCGTAACCAGCAAATCAATATCACTGTGTGGCTTCAGGCCGCCATCCACTGCGGAGCCGTACAAATGT
ACGGCCAGCAACGTCGGTTCGAGATGGCGCTCGATGACGCCAACTACCTCTGATAGTTGAGTCGATACTTCGGCGATCACCGCTTCCCTCATAATGT
TTAACTTTGTTTTAGGGCGACTGCCCTGCTGCGTAACATCGTTGCTGCTCCATAACATCAAACATCGACCCACGGCGTAACGCGCTTGCTGCTTGGA
TGCCCGAGGCATAGACTGTACCCCAAAAAAACAGTCATAACAAGCCATGAAAACCGCCACTGCGCCGTTACCACCGCTGCGTTCGGTCAAGGTTCT
GGACCAGTTGCGTGAGCGCATACGCTACTTGCATTACAGCTTACGAACCGAACAGGCTTATGTCCACTGGGTTCGTGCCTTCATCCGTTTCCACGGT
GTGCGTCACCCGGCAACCTTGGGTAGCAGCGAAGTCGAGGCATTTCTGTCCTGGCTGGAACAGAACTTATTATTTCCTTCCTCTTTTCTACAGTATT
TAAAGATACCCCAAGAAGCTAATTATAACAAGACGAACTCCAATTCACTGTTCCTTGCATTCTAAAACCTTAAATACCAGAAAACAGCTTTTTCAA
AGTTGTTTTCAAAGTTGGCGTATAACATAGTATCGACGGAGCCGATTTTGAAACCGCGGTGATCACAGGCAGCAACGCTCTGTCATCGTTACAATC
AACATGCTACCCTCCGCGAGATCATCCGTGTTTCAAACCCGGCAGCTTAGTTGCCGTTCTTCCGAATAGCATCGGTAACATGAGCAAAGTCTGCCGC
CTTACAACGGCTCTCCCGCTGACGCCGTCCCGGACTGATGGGCTGCCTGTATCGAGTGGTGATTTTGTGCCGAGCTGCCGGTCGGGGAGCTGTTGG
CTGGCTGGTGGCAGGATATATTGTGGTGTAAACATAACGGATCCGGTCTCAGGAGAATGAGGAGCACCGTCCAAACCTGGATAGCCGATATGTGAG
TGTTTGGTTGCCTGTCATTTCTAAAAAGCGTATGTCTCGTAGCGTAGATAGCAAGCGCTGTGAGAAAGGTGTGCTCCTATAGCATGGTAGCATCTGT
CCCATATCTCGTCCGCGAGAAGCCTACATTTCTCGTTGCGTATTAGAGCATGAAAAACCAACGTGAAGTACGTGATTCAGACCTTGACCAGTTCCAG
TTCTCTTCAACAAAAAAACAGTTCTTCAATTCATTTGCTTACATTAAACAATTGTCACACTCTTCTTCCCGGATGACAGCACCAAGACACTTCCCC
TCCACAAAAATCTCCGAGTCATGTGGACCGGCCTGGATGCTCTTAAAGGGGTTGCCATTGTAGCGGGGTGCCTTAACCACGACCTCTTCGATACCTG
TCTGAAGCGGCGTGTGAAATCCTCCAGGAGCTTGTTCAAATTTACAAGCTGCAATGGATGTAAACTCATCCAAAGACATGACTTGCGAGAAGCCAG
AAACGGTCATCAACACTACGAACACCCCTAAACATGGATGGAGAGGCAACATCCTGCATGTGTCTATGTCAGAGGATGCAAAAGTGAAAGACAAAG
GAGAAGGATTGCTCGTGCAATCACTGCAAAGCAAGATGCGAATATCCCTAAAATCCACAGCAGGAAAAGAAAGGAGCCTCAATAAGTCTCTAAAAT
GGGAAGAAAAGAAGCACGGGCCAGCGAGACGGAAATGAGGCGATCCGATGCGATGGTTCCGAGAACTGAAGCGGTGCGTGGGGGGCTAGAAATGA
ATGGGGAACGTGGAGGGCAAACAAAGGCTGTACTTAGCGGCAAATGCAGATGGCATGCATGCCAAGAACAGCAGCAGTCGGCACCGCCAGTTTAG
CGGAGAGCAACAGGCAGCAAAAGGAAGTGGAACAGGTTTAGGGCCGACGCAGTCGGCGCCTGCCCTAAAACGGCAAGGCCCACGGCAGCGGTCGC
GAGGGCGTTGCCTGCCGGTCGTGGAAAGCCGTGGGGCTGTCAACGGAACCTCACGCTCCCAGCCTGCCCTAAGCGGGGAAAAAGAAGATCCACATT
CTAGGGCTTTCCATCTCGGCTCTCAGCCGTCCGATTAAGAATCGCACGGCCTAGCCCCGCCAGCCTAGCCCTCGGCCGCCCCCTTGACAGGTTAAGA
GCTGGGCAAGCGACTCAGTTGTTCCTTCGTCGCTCGCCCTTCTCACGAGCAGCTCGCCTGCCCCGCCGCCGCCTTGCCTGCCTCGCGCCTTTTGTCT
CCAGGTTCCCTTGCCGCGCCTTCTCCTTGGCTTGTGCTTCTGCTTTGCTTGTGCCTCTGCTGGGCTCTACTTGTGTGGTGTTGTGCTTTGGCGCCGC
CGCCTCGCTGCTGCTGTGGAGCTTGGTGTGTGCTGGGGTTTCTGGGGTTTGCATGTGTCAGGGTTTGTAGATCTGTTCTTCGGGGCCGTCTTTCTA
ACGTTTCGTGTGCCGTTGTTTCTGGGGGGTTTGGTGTGGAGCTCGCAGCTTGCAGCTAATGAAAAAGCCTGAACTCACCGCGACGTCTGTCGAGAA
GTTTCTGATCGAAAAGTTCGACAGCGTCTCCGACCTGATGCAGCTCTCGGAGGGCGAAGAATCTCGTGCTTTCAGCTTCGATGTAGGAGGGCGTGG
ATATGTCCTGCGGGTAAATAGCTGCGCCGATGGTTTCTACAAAGATCGTTATGTTTATCGGCACTTTGCATCGGCCGCGCTCCCGATTCCGGAAGTG
CTTGACATTGGGGAGTTTAGCGAGAGCCTGACCTATTGCATCTCCCGCCGTTCACAGGGTGTCACGTTGCAAGACCTGCCTGAAACCGAACTGCCC
GCTGTTCTACAACCGGTCGCGGAGGCTATGGATGCGATCGCTGCGGCCGATCTTAGCCAGACGAGCGGGTTCGGCCCATTCGGACCGCAAGGAATC
GGTCAATACACTACATGGCGTGATTTCATATGCGCGATTGCTGATCCCCATGTGTATCACTGGCAAACTGTGATGGACGACACCGTCAGTGCGTCCG
TCGCGCAGGCTCTCGATGAGCTGATGCTTTGGGCCGAGGACTGCCCCGAAGTCCGGCACCTCGTGCACGCGGATTTCGGCTCCAACAATGTCCTGA
CGGACAATGGCCGCATAACAGCGGTCATTGACTGGAGCGAGGCGATGTTCGGGGATTCCCAATACGAGGTCGCCAACATCTTCTTCTGGAGGCCGT
GGTTGGCTTGTATGGAGCAGCAGACGCGCTACTTCGAGCGGAGGCATCCGGAGCTTGCAGGATCGCCACGACTCCGGGCGTATATGCTCCGCATTG
GTCTTGACCAACTCTATCAGAGCTTGGTTGACGGCAATTTCGATGATGCAGCTTGGGCGCAGGGTCGATGCGACGCAATCGTCCGATCCGGAGCCG
GGACTGTCGGGCGTACACAAATCGCCCGCAGAAGCGCGGCCGTCTGGACCGATGGCTGTGTAGAAGTACTCGCCGATAGTGGAAACCGACGCCCCA
GCACTCGTCCGAGGGCAAAGAAATAGGCTTGAGCTCGAATTTCCCCGATCGTTCAAACATTTGGCAATAAAGTTTCTTAAGATTGAATCCTGTTGC
CGGTCTTGCGATGATTATCATATAATTTCTGTTGAATTACGTTAAGCATGTAATAATTAACATGTAATGCATGACGTTATTTATGAGATGGGTTTTT
ATGATTAGAGTCCCGCAATTATACATTTAATACGCGATAGAAAACAAAATATAGCGCGCAAACTAGGATAAATTATCGCGCGCGGTGTCATCTATGT
TACTAGATCGATCCGTATCGATAGCCTCTAGCTAGAGTCGATCGACAAGCTCGAGTTTCTCCATAATAATGTGTGAGTAGTTCCCAGATAAGGGAAT
TAGGGTTCCTATAGGGTTTCGCTCATGTGTTGAGCATATAAGAAACCCTTAGTATGTATTTGTATTTGTAAAATACTTCTATCAATAAAATTTCTAAT
TCCTAAAATCAAAATCCAGTACTAAAATCCAGATCGCTAGCAAGGAGAGTTCATCTGTCAGCACTTGGACAGACCTTCCGGATCAAAGGTCGCTAT
TGCTATTCGCAGATCTAAAGTTTGAACGGCATTCACCGAGTGGTGGATGCTTCAACTGGCTTCTGGCACCGAGCGTAAAAGGACAAATCCAACGTA
CATATCGCTAAATTTCGTTCCAACGTACATATCGCTAATTTTTCCTACGGGAGGAATTAATCCTCGATAAACTCGGTGCCTTCTCGAAAGATGGTTT
CACCATGGTTCAACTCCATGGCTCACCCCGACCCCACGCTTATCCCACGGCTTCCAACACTCAGGGTGCTCGACTCCTATTACTCCCCGACCGTCTTC
TTCGGAAAATCTTTATGCGCATTCCCCATCCTAACTTGGAGATGCGATTGCTTCATCCGCCTTCCTGTTCATCACTTCTCATCACCCCTCATTCGCA
AAGAAATATCGTGGCACCTGGGATATGGAAATTTGCCCACTCTGCCCTAGACTCTGAAAATCAGGGATGAACAAAAGGAATGACCGACAATCTTTG
TGGAAAAGCTCTGGACAGGACTAGACTACCAATTGTAACCTGAAACCTTGCTCTTTGAATGGGGAAATTCCTTACATGAGTCCTAATTGAGAAATC
TGTCGCCCTACAAAGTTGCAAATGGCAGCTCTGGCTATGGCAGCTCTAGCAATGGCCAGGCAAATGGCAGCTCTAGCAATGGCCAGGCAAATGGCA
GCTCTAGCTATGGCCAGGCGGGCAAGGCCGTGATCTTCTTCCAGAGCGAGCTCGGGCATACAGGCGGGGCCAGCATAGAGGGCTCACTATTTCCTCC
AGCTGGGAGGCCTAGGGAAGGAAGGTCTGTGCCGCTGGCTATGGCGGTCGACTTGTGTGGGCCCTCCGAAGAACGAGCCAAAACTCTGAGCCAGTC
GTTGCGCAGCCAATGAGCGGAGCTGTGGATGCTTGAACTGCGTTTTGTCATTCGATTATGTATTGATTAGCAACTCAAATGATAAGTCGATCTCTC
AGGTAATACATAGTCCATGGTTTGAGTTCAAATTAGTCGCTCAAAAAACGAGGCTTTGGAGAAATCCAGCGAGGTTGGCGATCCTTTCTTATCCTT
CCCTGCCAACTCTCTTCGTGTGGCCGTGAACGCTCTCTCGTTGCCCCTATCCTATTTGAGCCCGGGCGGGCCACTCTCCAATCCAGTCCGCCCTCTCC
CGGCCCTCTCCTTTTTCTTCCCCGGTAGCAGGCATACATCTTGCGCTTCTCCGGTTCCATCCACACTGCAGCAGCAATGGCCGCAACAGCGGGGCTT
GCGCTTCCTTGCAGGTCTGCCCTGGATTTGGGCGGGTCGGGGTCGGCTTCGGGCGCAAGGCAGGTCAAGGCGGGCTCCGTGGCCTTCCTCGGGGAG
GCGGAGGGTCTATGGCGGGCCAAGGTGTCTGTTGCGGGCGTTGTGTGGAGGAGGAAGGTGGGCAGTAAAGTTCCTGTTTCCCGGTCGTATTTCAAT

CCTGTGGAGGCGGACCAGCAGAAGGAGGAGGCGAGGCTGTACAGGCGGACGGTCTACGACCACGACCAGTGGAGGAGGCACCGGAGCAGCACCCG
CCATGCCCGCCACATTCTCTCCATCGCCTCCTCCAGGGTGATACTCTCGCTGGGGCCCCCGGTGTACAGCCTCACCGTCTTCTCTGCCGCCGTTGCGC
TCTACAACGAGATTGTGTCGGACAAGTTCTTCCCGCTGCCGGAATGGCTGCCCATTCTGCACATCTCGCCGCTGCCCATCCAGTTGTCGGCTCCGGC
TCTGGCCCTCCTGCTGGTCTTCCGGACCAACTCCTCCTACGGGAGGTTCGACGAGGCCCGGAAGATCTGGGGGTCCAATGTGAACAGGACCAGGGA
TATTGCTCGGCAGGCCTTGTCCTGGATCGATGCTCCTGAAGACTCCGGCAAGCTGAAGGAGTTTCTGCGCTATGTGATTGCGTACCCGTACTGCCTT
AAGGATCACCTTGTCCAGGAGGATGATCTGCGGAAGGAGCTTGCTCCTATTCTCCAGAAGAACGAGCTGGACAAGTTGCTGGCCTCCAATCACCGA
CCTAACCATGCTGTGCAGATGATGTCGGACACTCTTGCCAAATGCCAGTTGTCGGATATCAACAGGTCTTCCATGGATCACAACCTGACCCAGTTCC
ACGACAACACGGGGGCGTGTGAACGTATCTTCAAGACCCCAATTCCTGTGTCCTACACACGGCTTACATCCCGTTTCCTGGTTCTGTGGCACATTGT
TATGCCTTTCTGCCTCTGGGATTCTTGCCGGCTGCTGCTGATCCCCGCAACCTTCGCTAGCGCAGCGGCCTTGATGTGCATTGACGAGGTTGGTGTT
CTGATAGAAGAACCGTTCTCAATTCTCGCCCTGCCAGCTCTATGCCAGGGCATGCACAAGGGTGTTGAGGGCCTTCTCCTGTCCCATGCTGACTGGC
CTAGCCTCAGATCTGCCGGTGAGCGAGTTCCAGTTGGAGTGCAGACCGTGAATGGCGTTAAGGCTTCGGTGAGCAAGGGCGAGGAGCTGTTCACCG
GGGTGGTGCCCATCCTGGTCGAGCTGGACGGCGACGTAAACGGCCACAAGTTCAGCGTGTCCGGCGAGGGCGAGGGCGATGCCACCTACGGCAAGC
TGACCCTGAAGTTCATCTGCACCACCGGCAAGCTGCCCGTGCCCTGGCCCACCCTCGTGACCACCCTGACCTACGGCGTGCAGTGCTTCAGCCGCTA
CCCCGACCACATGAAGCAGCACGACTTCTTCAAGTCCGCCATGCCCGAAGGCTACGTCCAGGAGCGCACCATCTTCTTCAAGGACGACGGCAACTAC
AAGACCCGCGCCGAGGTGAAGTTCGAGGGCGACACCCTGGTGAACCGCATCGAGCTGAAGGGCATCGACTTCAAGGAGGACGGCAACATCCTGGG
GCACAAGCTGGAGTACAACTACAACAGCCACAACGTCTATATCATGGCCGACAAGCAGAAGAACGGCATCAAGGTGAACTTCAAGATCCGCCACAA
CATCGAGGACGGCAGCGTGCAGCTCGCCGACCACTACCAGCAGAACACCCCCATCGGCGACGGCCCCGTGCTGCTGCCCGACAACCACTACCTGAGC
ACCCAGTCCGCCCTGAGCAAAGACCCCAACGAGAAGCGCGATCACATGGTCCTGCTGGAGTTCGTGACCGCCGCCGGGATCACTCTCGGCATGGAC
GAGCTGTACAAGTAAGCTTGAGCTCGAATTTCCCCGATCGTTCAAACATTTGGCAATAAAGTTTCTTAAGATTGAATCCTGTTGCCGGTCTTGCGA
TGATTATCATATAATTTCTGTTGAATTACGTTAAGCATGTAATAATTAACATGTAATGCATGACGTTATTTATGAGATGGGTTTTTATGATTAGAGTC
CCGCAATTATACATTTAATACGCGATAGAAAACAAAATATAGCGCGCAAACTAGGATAAATTATCGCGCGCGGTGTCATCTATGTTACTAGATCGATC
CGTATCGATAGCCTCTAGCTAGAGTCGATCGACAAGCTCGAGTTTCTCCATAATAATGTGTGAGTAGTTCCCAGATAAGGGAATTAGGGTTCCTATA
GGGTTTCGCTCATGTGTTGAGCATATAAGAAACCCTTAGTATGTATTTGTATTTGTAAAATACTTCTATCAATAAAATTTCTAATTCCTAAAATCAA
AATCCAGTACTAAAATCCAGATCGCTATACAGGAGTCTGAGCAGGACAACTCGCGTAGTGAGAGTTACATGTTCGTTGGGTTCTTCCGACACGGACC
TGAGTTGGCCAACGTCCCACCTGAGGTCTGTGCCCCGGTGATGAGAAGTGTGCATCTCGTTCTTGCAGCTCGTCAGTACTTTCAGAATCATGGCGT
GCATGGTAGAATGACCCTTATAACGGACTTCGACATGGCAATCGCTACAGAGGAGTCTGAGCAGGACAACTCGCGTAGTGAGAGTTACATGTTCGT
TGGGTTCTTCCGACACGGACCTGAGTTGGCCAACGTCCCACCTGAGGTCTGTGCCCCGGTGATGAGAAGTGTGCATCTCGTTCTTGCAGCTCGTCA
GTACTTTCAGAATCATGGCGTGCATGGTAGAATGACCCTTATAACGGACTTCGACATGGCAATCGCTAGGT
